# Chromosomal curing drives an arms race between bacterial transformation and prophage

**DOI:** 10.1101/2024.08.13.607808

**Authors:** Min Jung Kwun, Alexandru V. Ion, Katinka J. Apagyi, Nicholas J. Croucher

**Affiliations:** MRC Centre for Global Infectious Disease Analysis, Department of Infectious Disease Epidemiology, School of Public Health, Imperial College London, London, W12 0BZ, UK

## Abstract

Transformation occurs when bacteria import exogenous DNA via the competence machinery and integrate it into their genome through homologous recombination (HR). This process may provide an evolutionary advantage to cells through enabling “chromosomal curing”: the deletion of integrative mobile genetic elements (MGEs). However, many such MGEs are sensitive to RecA-DNA filaments, triggering activation of a lifecycle that may enable them to evade HR-mediated deletion. Despite >40% of isolates containing prophage integrated at a site that inhibits transformation, three representative prophage were identified in naturally-competent pneumococci to test this hypothesis. These encompassed representatives with C1-type and ImmAR-type regulatory systems, found in almost all pneumococcal prophage. All three prophage were deleted by HR with an efficiency similar to the transfer of base substitutions. Mutations that impaired a C1-regulated prophage increased this deletion rate, reflecting this element being activated by RecA-DNA filaments imported during transformation, likely preferentially killing cells that induce competence. ImmAR-regulated prophage instead responded to transient stimuli by excising as deletion-resistant pseudolysogens, only driving cell lysis in response to sustained stimuli. This was likely a consequence of these prophage reacting to multiple signals, as they differed in their response to both RecA and the DNA-binding protein and competence repressor DprA. One prophage constitutively elevated host DprA levels, thereby reducing transformation by preventing induction of the competence machinery. Hence these data are consistent with an evolutionary arms race between prophage and the competence machinery, resulting in bacterial diversification though HR being impeded by MGEs preventing their own elimination from the chromosome.

## Introduction

The population of viruses that infect bacteria comprises bacteriophage, the most abundant biological entities worldwide (Wommack and Colwell 2000; Chevallereau et al. 2022), and prophage, dormant sequences found in most bacterial genomes (Silveira et al. 2021). Both population genomic and metagenomic data has revealed these viruses to be highly diverse (Moura de Sousa et al. 2021). Despite this variation, the induction of prophage to generate phage by the genotoxic streptomycete-produced antibiotic mitomycin C (MMC) is common across the global microbiome, including samples from animal and human microbiotas (Minot et al. 2011; Kim and Bae 2018), soil (Williamson et al. 2007; Roy et al. 2020), freshwater (Maurice et al. 2011), seawater (Paul 2008), and subseafloor sediments (Engelhardt et al. 2011). Correspondingly, MMC induction of prophage and other types of mobile genetic element (MGE) (Auchtung et al. 2007; Bose et al. 2008) are observed across most bacterial phyla (Knowles et al. 2017; Silveira et al. 2021) and archaea (Chen et al. 2020; Fusco et al. 2020). Hence the capacity to respond to MMC is disseminated across a genetically diverse set of MGEs found within the full taxonomic and ecological range of prokaryotic hosts.

This broad distribution is the result of convergent evolution of multiple regulatory mechanisms (Sauer et al. 1982; McLeod et al. 2005; Bose et al. 2008; Chen et al. 2020; Fusco et al. 2020) that detect the RecA-coated single-stranded DNA (ssDNA) lesions generated after cells attempt to replicate DNA crosslinked by MMC (Payne-Dwyer et al. 2022). In many bacteria, these RecA-ssDNA structures catalyse the autodigestion of a LexA transcriptional repressor, activating the DNA repair genes of the SOS response (Erill et al. 2007). These structures also activate the LexA-like C1 repressor protein of *Escherichia coli* phage λ, the archetypal MMC-inducible MGE (Ptashne et al. 1982). *E. coli* prophage 186 is instead activated by LexA cleavage derepressing expression of Tum, which inhibits a repressor (Shearwin et al. 1998). Similarly, the CTXϕ prophage of *Vibrio cholerae* and GIL01 prophage of *Bacillus thuringiensis* are directly repressed by LexA binding the prophage in conjunction with a phage protein (McLeod et al. 2005; Fornelos et al. 2015). By contrast, prophage SSV1 of the archaeon *Saccharolobus shibatae* is proposed to be repressed by RadA, the archaeal orthologue of RecA, binding the prophage until it is removed through a higher-affinity interaction with ssDNA (Fusco et al. 2020). The sensitivity of other MGEs to MMC depends on regulatory machinery that structurally resembles the IrrE/PprI-DdrO regulators (de Groot et al. 2019) that control DNA repair genes in the *Deinococcaceae* and *Thermaceae* families (Ludanyi et al. 2014; Sadowska-Bartosz and Bartosz 2023). These use a bipartite system that features a ImmR transcriptional repressor (Auchtung et al. 2007) that can be degraded or displaced by an MGE-encoded ImmA protease (Bose et al. 2008; de Groot et al. 2019).

Yet prophage remain MMC-sensitive even in the bacterium *Streptococcus pneumoniae* (Martin et al. 1995) (the pneumococcus), which lacks LexA (Erill et al. 2007), a canonical SOS response (Claverys et al. 2000), or any well-characterised IrrE/PprI-DdrO regulators (Sadowska-Bartosz and Bartosz 2023). Pneumococci themselves have been found to respond to MMC by inducing competence for natural transformation (Prudhomme et al. 2006), enabling the bacteria to import exogenous DNA for integration into the chromosome through homologous recombination (HR) (Johnston et al. 2014). Although pneumococci frequently contain prophage (N. Croucher, Coupland, et al. 2014; Brueggemann et al. 2017) and other integrative MGEs (Croucher et al. 2009; N. Croucher, Coupland, et al. 2014; Martínez-Rubio et al. 2017; D’Aeth et al. 2021), both β-lactam resistance (Dowson et al. 1993) and vaccine evasion (N. Croucher, Chewapreecha, et al. 2014) are instead facilitated through transformation. This process is enabled by expression of the competence machinery, the activation of which is coordinated by competence stimulating peptide (CSP), a quorum sensing pheromone (Håvarstein et al. 1995). This signal induces competence in a fraction of cells, as the overall population engages in “bet hedging” (Veening et al. 2008; Kwun et al. 2023; Prudhomme et al. 2024). The competent subpopulation first induces early competence genes, such as *comX*, which encodes a competence-specific RNA polymerase sigma factor (Straume et al. 2015). ComX then binds combox sequence motifs, driving the expression of late competence genes that encode the machinery necessary for DNA import and HR (Straume et al. 2015), including a ComY (alternatively named ComG) pilus, able to contact exogenous DNA (Sandra et al. 2019); a ComE pore complex, which enables the import of single-stranded DNA (ssDNA) fragments, and ssDNA-binding proteins, such as DprA (Johnston et al. 2023) and SsbB (Attaiech et al. 2011), which protect and store intracellular ssDNA. However, pneumococcal competence is only transient, as after ∼20 minutes is it shut down by DprA (Mirouze et al. 2013). Despite this extensive regulation, the primary evolutionary advantage of transformation remains controversial (Redfield 1993; Johnston et al. 2014).

One hypothesis proposes the main function of transformation is “chromosomal curing”, the elimination of deleterious MGEs from the genome through HR (Croucher et al. 2016). This is supported by the high efficiency with which non-essential sequences can be removed from the pneumococcal chromosome by transformation, which is not matched by a comparable ability to insert equivalently-sized loci (Apagyi et al. 2018). Hence transformation is asymmetric, implying it has the potential to cure cells of MGEs integrated into their chromosome, with little risk of inserting an additional element. This is the theoretical basis for the intragenomic conflict between prophage and the competence machinery (Croucher et al. 2016).

Yet integrative MGEs can only be deleted by HR when they are inserted into the chromosome at their *attB* (bacterial attachment) site, flanked by regions with similarity to the imported DNA. However, following activation, most prophage excise from the genome as extrachromosomal circular forms (McLeod et al. 2005; Mäntynen et al. 2021) that cannot be removed by HR. In the C1-regulated phage λ, this excision is traditionally regarded as the first stage of a rapid, irreversible transition from a lysogenic to lytic life cycle (Ptashne et al. 1982). In other phage, the phage may be detected in a non-replicating circularised form, termed a pseudolysogen (Łoś and Węgrzyn 2012; Mäntynen et al. 2021). The low copy number renders such a form unstable (Ogunseitan et al. 1990; Ripp and Miller 1997; Sakaguchi et al. 2005; Lood and Collin 2011; Cenens et al. 2013), meaning pseudolysogeny is typically regarded as a stalled phase associated with starvation conditions (Wommack and Colwell 2000; Łoś and Węgrzyn 2012; Mäntynen et al. 2021). Nevertheless, it has the potential to affect the efficiency of deletion by HR.

In this work, we test whether transformation is an efficient mechanism for deleting prophage despite their sensitivity to the ssDNA the competence machinery imports (Fig. 1A). Our analyses identify the key interfaces in the evolutionary arms race between prophage and their hosts, and how this affects overall pneumococcal epidemiology.

**Figure 1.**
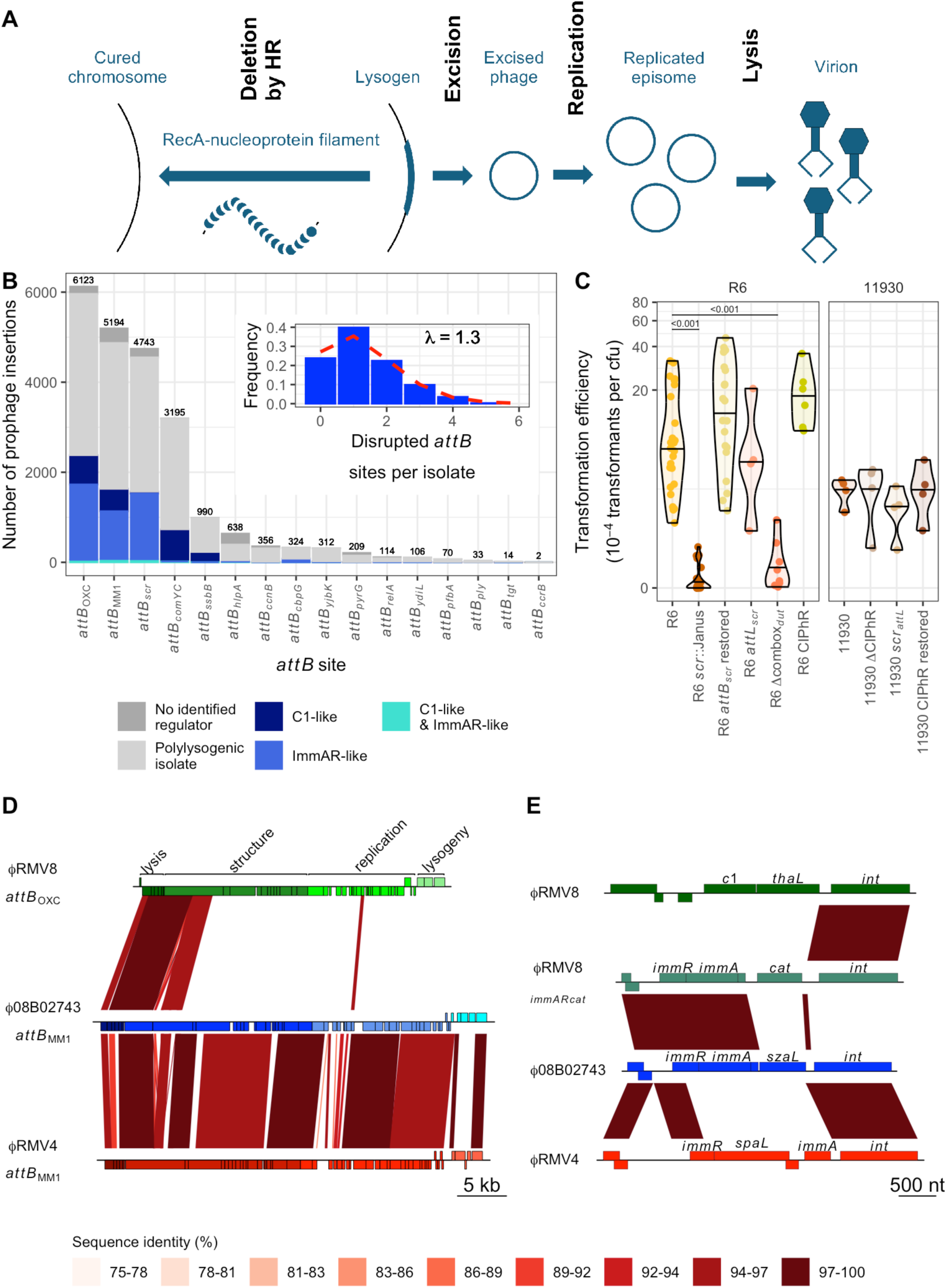
Genomic characterisation of pneumococcal prophage. **(A)** Summary of the intragenomic conflict between cell-driven HR and prophage activation. **(B)** Frequency of prophage insertion at the 16 identified pneumococcal *attB* sites across the 20,047 isolates of the Global Pneumococcal Sequence project collection. Bars are coloured by the associated prophage regulatory system, which was only inferred in monolysogenic isolates; otherwise, they are coloured grey, as it was not feasible to reliably link regulators and *attB* sites in draft assemblies of polylysogenic isolates. The inset shows the number of disrupted *attB* sites in a set of isolates corrected for the effects of population structure by randomly selecting one genome from each Global Pneumococcal Sequence Cluster in the isolate collection. **(C)** Effect of modifications at the *attB_scr_* site on transformation efficiency. Each point represents an individual transformation experiment using donor DNA containing a rifampicin resistance marker. The distribution of transformation efficiencies is summarised by a violin plot for each recipient genotype, with a horizontal line at the median value. The left panel compares mutants against the CIPhR-negative parental R6 genotype; the right panel compares mutants against their CIPhR-positive parental 11930 isolate. Statistical comparisons used a two-tailed Wilcoxon rank sum test, subject to a Holm-Bonferroni correction for multiple testing within each panel. Significant differences are shown by the horizontal bars at the top of the plot, annotated with the associated *p* values. **(D)** Alignment of the prophage, each labelled with its name and *attB* site. Protein coding sequences (CDS) are shown as boxes above the line, if encoded on the top strand, or below the line, if encoded on the bottom strand. Their colour indicates whether they were associated with lysogeny regulation, phage replication, virion structure or host cell lysis. The red bands linking the sequences mark regions of sequence similarity identified by BLASTN. **(E)** Alignment of lysogeny regulation loci. CDSs are annotated, and similarity between sequences indicated by bands, as in panel (A).

## Results

### Many pneumococcal genomes contain prophage inhibiting transformation

Analysis of 20,047 published Global Pneumococcal Sequencing (GPS) project genomes (Gladstone et al. 2019) (Text S1, Fig. S1) estimated 14,113 were infected by 22,682 prophage-like sequences (Fig. 1B, S2-S3), with the number per cell being Poisson distributed (Fig. 1B). These were identified inserting at 17 *attB* sites, at which 16 were not targeted by other MGEs (Fig. 1B, S4-S6; Table S2). However, the majority of insertions (19,255; 84.9%) occurred at the four most common *attB* sites (Romero et al. 2009; Garriss and Henriques-Normark 2020). Insertions at the most frequently-affected *attB* site, *attB*_OXC_, result in a modification of *ccnC* (Furi et al. 2019) that enhances the activity (Kwun et al. 2022) of the competence-suppressing non-coding RNA it encodes (Halfmann et al. 2007). Similarly, the fourth most commonly-targeted site disrupts the *comYC* competence pilus gene, necessary for transformation (Croucher et al. 2011; Croucher et al. 2014). Among the rarer *attB* sites identified in this analysis, two were highly likely to inhibit transformation. One inserted between the promoter and translation start site of *yjbK*, a positive regulator of competence (Kwun et al. 2023). The other, previously identified in *Streptococcus mitis*, disrupts the *ssbB* gene (Denapaite et al. 2010), required for efficient recombination when high concentrations of donor DNA are available (Attaiech et al. 2011).

A phylogeny of the integrases targeting all the *attB* sites identified seven clades of closely-related sequences (Fig. S7). This tree was consistent with each competence-inhibiting *attB* site being targeted by an independently-evolved integrase, which has subsequently diversified to target rarer *attB* sites not predicted to limit DNA uptake. The exceptions were *attB*_MM1_ (Romero et al. 2009), which duplicates the stop codon of *whiA* without affecting the encoded protein’s sequence, and *attB_scr_*, within the *scr* gene encoding the small cytoplasmic RNA (scRNA), which is necessary for efficient transformation (Lin et al. 2024). Although there is variation in the annotated length of pneumococcal scRNA genes, qRT-PCR experiments concurred with the recent inference that it is 79 nt in size (Fig. S8) (Lin et al. 2024). This positioned the *attB_scr_* site at the 3’ end of the gene, overlapping a pyrimidine-rich region that, in conjunction with the adjacent combox motif, drives the competence-associated transcription of a downstream operon containing the *dut* and *radA* genes (Peterson et al. 2004). Prophage integrations at *attB_scr_* duplicated 22 bp, copying the 3’ end of *scr* and the pyrimidine-rich sequences on the *attR* and *attL* sides, respectively. As both the scRNA and *radA* genes affect competence (Burghout et al. 2007; Lin et al. 2024), mutations affecting the loci on either side of *attB_scr_* were constructed using the selectable and counter-selectable Janus cassette in the highly-transformable laboratory strain *S. pneumoniae* R6 (Li et al. 2014). Deleting the combox (mutant Δcombox*_dut_*) caused a ∼20-fold reduction in transformation efficiency (Fig. 1C). Disrupting the *scr* gene (mutant *scr*::Janus) resulted in a similar phenotype (Fig. 1C), as well as a substantial growth defect (Fig. S9). Restoring the cellular sequence (mutant *scr_attB_* restored) reversed both changes, as did introducing a ∼100 bp sequence corresponding to the *attL* junction generated by prophage insertions at this site (mutant *attL_scr_*). Hence, in contrast to some other pneumococcal *attB* sites, the *attB_scr_* integration site maintains the integrity of both flanking loci affected by prophage integration, each of which are required for efficient activation of competence.

The *attB_scr_* site contained an insertion in 23.7% of the analysed pneumococcal genomes, although 68.0% of these were not full-length prophage, but instead a ∼6.2 kb prophage-related sequence (Croucher et al. 2014) (Fig. S1). The archetype of these elements, identified in an isolate belonging to the PMEN1 lineage (Croucher et al. 2009), encodes a truncated lytic amidase, suggesting it is incapable of lysing its host cell; a regulatory protein, and an integrase gene interrupted by a frameshift mutation in a polyadenine tract. Despite this disruptive mutation, PCR amplification of DNA from a PMEN1 isolate demonstrated that the element was able to excise from the chromosome, although quantitative assays indicated this only occurred in <0.01% of cells, even after exposure to MMC (Fig. S7). As its only apparent cargo is a gene of unknown function, this cryptic element was termed a Competence-associated loci Intervening Prophage Remnant (CIPhR). To determine whether this element affected the transformation efficiency of the host, it was introduced into *S. pneumoniae* R6 (mutant R6 CIPhR). This mutant exhibited a similar growth profile and transformation efficiency to the unmodified genotype, indicating the integration does not affect the functioning of the *scr* gene or *dut* combox (Fig. 1C). Similarly, no change in transformation efficiency was observed when CIPhR was removed from the PMEN1 representative 11930, a natural host genotype, and either replaced with the wild-type *scr* locus, or the modified *attL_scr_* sequence (Fig. 1C). Therefore insertions at *attB_scr_* appear to precisely avoid disrupting the cell’s ability to induce competence, whereas almost half of pneumococcal genomes (8,843 isolates; 44.1%; Fig. S3) contained MGE insertions at *attB* sites that are expected to inhibit transformation. This suggests prophage may employ different strategies to evade deletion by HR.

### Pneumococcal prophage are primarily regulated by two types of system

Re-analysis of a previously-defined set of 538 prophage-derived sequences (Croucher et al. 2013) found full-length elements contained one of two mutually-exclusive regulatory systems: a C1-like protein, or an ImmAR-like system (Text S1, Fig. S10). A search of the GPS isolates found 9,841 (49.1%) contained an ImmAR-type regulator, and 7,402 (36.9%) contained a C1-type regulator. The distribution of these systems across *attB* sites in monolysogenic isolates was heterogeneous (𝜒^2^ test, *p* < 10^-16^; Text S1, Fig. 1B). The ImmAR-type systems were more prevalent at the three most commonly-infected sites (*att*_OXC_, *att*_MM1_ and *attB_scr_*); correspondingly, CIPhR elements encoded this type. C1-type systems dominated at the sites that disrupted protein-coding genes that were part of the competence machinery (*att_comYC_* and *att_ssbB_*). This suggested that the ImmAR-type and C1-type regulators differed in their interaction with cellular transformation.

Consequently, three model systems were developed in which the prophage were active and the cells were transformable (Table S1). *S. pneumoniae* RMV8 (Kwun et al. 2018) hosted a prophage encoding a C1-type regulator inserted at *attB*_OXC_, a chromosomal alteration that does not affect competence induced exogenously through addition of CSP (Kwun et al. 2022) (Fig. 1D). This phage’s lysogeny module also included a gene annotated as *thaL*, for “transmembrane HIRAN-domain protein associated with lysogeny” (Fig. 1E). *S. pneumoniae* 08B02743 (Croucher et al. 2016) hosted a prophage encoding an ImmAR-type regulator inserted at *att*_MM1_, the lysogeny module of which included a gene annotated as *szaL*, for “surface-exposed zinc metalloprotein associated with lysogeny”. Finally, *S. pneumoniae* RMV4 (Kwun et al. 2018) contained a prophage encoding an ImmAR-type regulator inserted at *att*_MM1_, the lysogeny module of which that included a gene annotated as *spaL*, for “surface-exposed zinc protein associated with lysogeny”. The ϕ08B02743 and ϕRMV4 prophage were highly similar outside of their regulatory loci, where ϕRMV8 only shared sequence similarity in the structural and lytic regions of the prophage. To improve the reproducibility of these systems (Furi et al. 2019; Kwun et al. 2022; Kwun et al. 2023), the phase-variable restriction-modification systems (Manso et al. 2014; Kwun et al. 2018) were deleted or fixed in a stable arrangement in all three (Text S2, Fig. S11-12).

To quantify the level of prophage activity enabled by these divergent regulatory systems in the absence of stimuli, the *attB*-to-*attL* ratio was used to measure the prevalence of bacteria in which the prophage had excised, relative to those in which a prophage remained integrated, following overnight growth in unsupplemented media (Fig. 2A). All three prophage exhibited similar levels of excision, suggesting the C1-type and ImmAR-type systems did not differ in their repression of the viral sequences in unstimulated hosts (Fig. 2B). Correspondingly, all the prophage were inferred to impose a fitness cost on their host cells even in unsupplemented media, as mutants in which the viral sequence was replaced with a Janus cassette grew to a higher cell density in stationary phase than the corresponding lysogens (Fig. 2C). Targeted disruption of the prophage integrase (*int*), replication (*rep*) and lytic (*lyt*) genes with a chloramphenicol acetyltransferase (*cat*) resistance gene resulted in growth phenotypes intermediate between those associated with genotypes in which prophage were intact or removed, consistent with these mutations impairing phage replication. To test whether these prophage were inducible, they were exposed to an MMC concentration that did not affect the exponential phase growth of the non-lysogenic genotype R6, or its *recA*^-^ mutant (Fig. S13). Both ϕRMV8 and ϕ08B02743 exhibited a post-MMC reduction in growth that suggested the prophage were strongly activated by MMC, whereas the response of ϕRMV4 was consistently weaker (Fig. 2D, S14-S15). Hence these three systems represent naturally transformable cells containing active viruses that differ in their response to MMC, and encode the two most common regulatory strategies employed by pneumococcal prophages.

**Figure 2.**
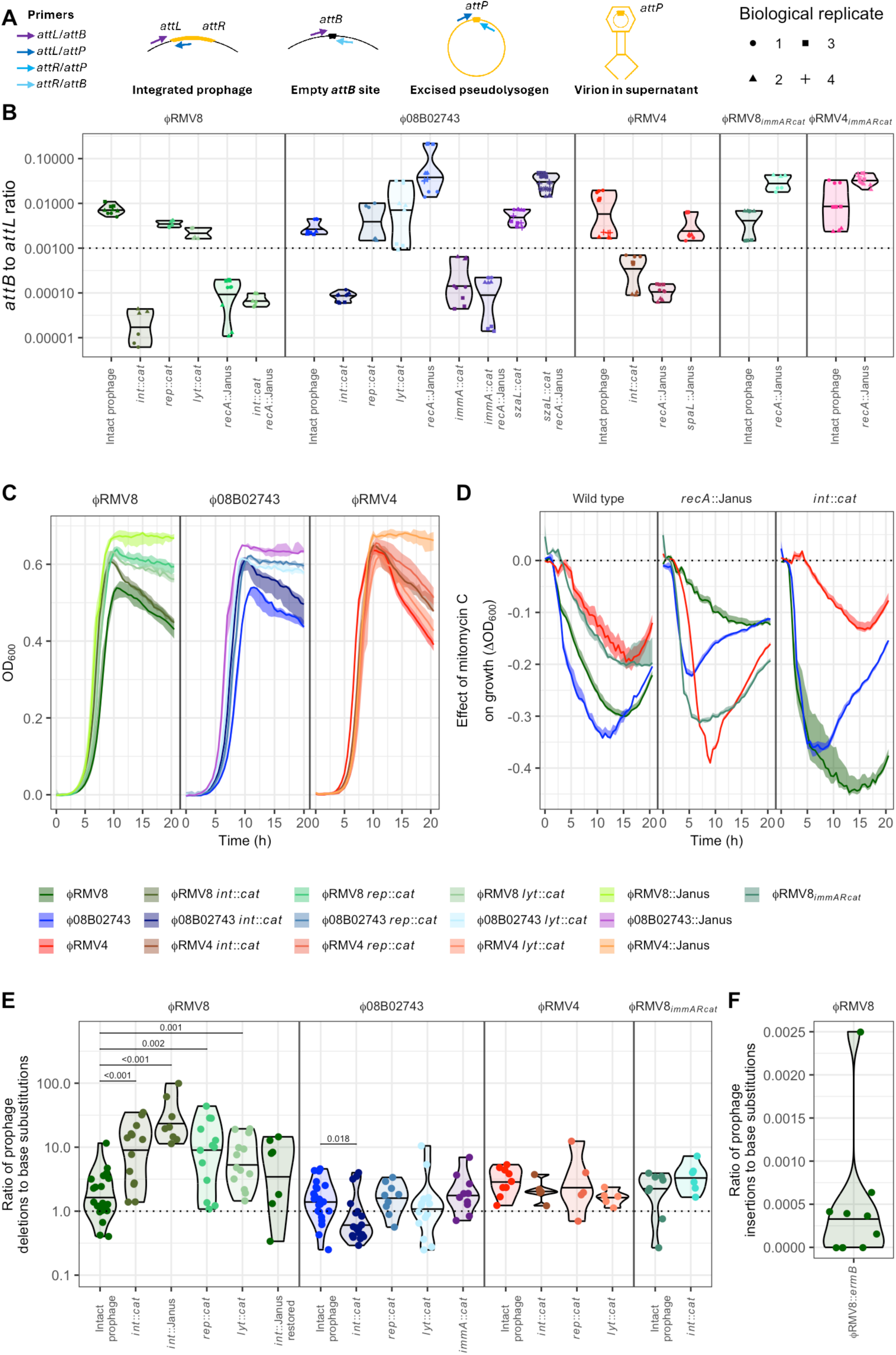
Efficient deletion of active prophage by transformation. **(A)** Diagram summarising the PCR amplicons used to analyse the topological changes associated with prophage activation. **(B)** The ratio of *attB* to *attL* sites, estimating the level of prophage excision, was measured by quantitative PCR following overnight growth in the absence of inducing stimuli. The horizontal axis separates the tested genotypes, grouped according to the prophage system, which is also indicated by the colour of the displayed data. Each point represents an individual technical replicate measurement, three of which were recorded for each biological replicate, as denoted by each point’s shape. The distribution of values for each genotype is summarised by a violin plot, with a horizontal line at the median. The horizontal dashed line at 10^-3^ distinguishes the actively-excising intact prophage from those that appear to be stably integrated, such as the *int*::*cat* mutants. **(C)** Growth curves comparing lysogens with cells cured of the prophage, and mutants with disruptions of the integrase (*int*), replication (*rep*) or lytic (*lyt*) loci, as annotated in Fig. 1D. **(D)** The effect of mitomycin C on the growth of prophage systems. The solid lines show the difference between the median OD_600_ of cells grown in mitomycin C and the median OD_600_ of the same genotype grown in parallel in unsupplemented media. The shaded area shows the range of OD_600_ values observed in cultures containing mitomycin C, similarly adjusted by subtracting the median OD_600_ of untreated cultures. **(E)** Deletion of prophage by transformation. The horizontal axis indicates the genotype of the host cells that were transformed with DNA from a donor genotype of the corresponding prophage system, in which the prophage was replaced by a Janus cassette, and the *rpoB* gene contained a base substitution conferring rifampicin resistance. The exception was the ϕRMV8 *int*::Janus mutant, which was instead removed by donor DNA in which the prophage was replaced by an *ermB* macrolide resistance marker. Each point represents an independent biological replicate in which the ratio of prophage replacement to base substitution acquisition was quantified through selection on appropriate media. The violin plots summarise the distribution and indicate the median value with a horizontal line. The vertical axis has a logarithmic scale, and the dashed line shows the value at which prophage are deleted as efficiently as base substitutions are exchanged. The efficiency with which each mutant prophage was deleted was compared to the equivalent values for the intact prophage of the same system using a two-tailed Wilcoxon rank sum test, with a Holm-Bonferroni correction for multiple testing within each panel. Significant differences are marked by horizontal bars, annotated with the calculated *p* value. **(F)** Insertion of prophage by transformation. DNA from a donor carrying a ϕRMV8*_immARcat_ int*::Janus prophage, and a base substitution conferring rifampicin resistance, was used to transform ϕRMV8::*ermB* cells. Each point represents the ratio of prophage insertions to base substitutions in an independent biological replicate. The distribution of these points is summarised using a violin plot, with a horizontal line at the median value.

### Pneumococcal prophage are efficiently deleted by transformation

To test whether transformation is an efficient mechanism for curing prophage from bacterial genomes, a base substitution causing rifampicin resistance was introduced into each mutant in which the prophage had been replaced with a Janus cassette. These double mutants were a source of high-molecular weight donor genomic DNA (Fig. S16) that could replace the prophage, resulting in aminoglycoside resistance, or transfer an unlinked single nucleotide polymorphism, resulting in rifampicin resistance (Fig. S17). The ratio of transformation with these two resistance phenotypes therefore enabled quantification of the efficiency with which prophage were deleted, relative to the transfer of a single base substitution (Apagyi et al. 2018). This ratio was measured two hours after the addition of the DNA, as this allowed for the resistance phenotypes to be expressed, but minimised the time for replication and selection to distort the ratio away from that reflecting the relative rates of mutation import (Fig. S18). For all three prophage systems, the median ratio of deletion to base substitution acquisition was above one (Fig. 2E). This contrasted with the rate at which transformation was able to insert complete prophage sequences, which was estimated as >10,000-fold lower than the rate at which base substitutions were imported (Fig. 2F). Therefore, transformation is an efficient mechanism for removing, but not importing, prophage.

These same deletion ratios were also calculated for the impaired prophage mutants, to test whether phage activation sometimes enabled the viruses to escape deletion (Fig. 2E). In the ϕRMV8 system, these mutations significantly increased the efficiency of prophage deletion ∼3-10 fold. This was primarily driven by a higher rate of prophage deletion, rather than a change in rifampicin resistance acquisition (Fig. S14). The high deletion rate of the ϕRMV8 *int*::*cat* mutant, which should remain embedded within its *attB* site (Fig. 2B), demonstrated efficient integration of the aminoglycoside resistance marker was not dependent on the excision of the prophage from the host chromosome. To confirm this higher rate was associated with the *int* mutation, the deletion rate was replicated in an *int*::Janus mutant, in which ϕRMV8 deletion was assayed by measuring replacement by an *ermB* macrolide resistance marker. The deletion rate then fell after the counter-selectable Janus cassette was replaced through restoration of the functional *int* gene (Fig. 2E). Hence the rate of deletion of the C1-regulated ϕRMV8 prophage depended on the ability of the prophage to activate.

By contrast, removal of the *int* gene decreased the efficiency with which the ϕ08B02743 prophage was deleted. Other mutations impairing ϕ08B02743 or ϕRMV4 did not change the frequency with which transformation removed these elements. To test whether this was attributable to the ImmAR-type regulator, the *szaL* gene of ϕ08B02743 was replaced with a *cat* gene, enabling the transfer of the regulatory locus of this prophage into the ϕRMV8 background to generate ϕRMV8*_immARcat_* (Fig. 1E). This hybrid prophage was inducible by MMC (Fig. 2D, S10), suggesting it was functional. This transfer did not substantially affect the rate at which the intact prophage was deleted, which remained similar to that of both the parental elements (Fig. 2E). In contrast to ϕRMV8, disruption of the *int* gene did not significantly elevate the deletion rate of ϕRMV8*_immARcat_*. This difference did not seem to be attributable to changes outside the prophage (Fig. S19), implicating the ImmAR-type regulators as the cause. Yet, the ImmA protease itself did not appear to have a direct role in suppressing the deletion of ImmAR-regulated prophage, as an ϕ08B02743 *immA*::*cat* mutant was just as efficiently removed from the genome as the intact prophage (Fig. 2E). Therefore the rate of deletion of the ImmAR-regulated prophage appeared to be independent of their ability to activate.

One hypothesis explaining these differences between the regulatory systems was that there existed a difference in their spatial sensitivity to the HR events enabled by transformation. As RecA-ssDNA nucleoprotein filaments are expected to interact with the C1-type protein, itself often tightly associated with the phage regulatory locus (Ptashne et al. 1982), it seemed likely that activation might be maximal when DNA specifically targeted the prophage element, as proposed for *Bacillus subtilis* ϕ105 (Garro 1973). This would contrast with the bipartite ImmAR-type regulators, in which ImmA could encounter stimuli throughout the cell, before interacting with prophage-bound ImmR. Hence impairment of the prophage encoding ImmAR-type regulators would reduce prophage-driven lysis of cells acquiring the Janus cassette or rifampicin resistance mutation equally. By contrast, impairment of ϕRMV8 would only increase the frequency of Janus cassette acquisition, as the prophage would not respond to HR importing rifampicin resistance at a distant locus. This was tested by transforming cells with PCR amplicons targeted at different parts of the prophage. Although PCR amplicons targeting the regulatory region did appear to activate ϕRMV8 more effectively than those annealing to other parts of the prophage, the same pattern was observed in ϕRMV8*_immARcat_* (Fig. S20). Hence the apparent spatial sensitivity of prophage activation was likely an artefact of local variation in recombination rate, as observed in other assays (Apagyi et al. 2018). Therefore the prophage deletion ratios were not explained by regulators’ differential spatial sensitivity to HR events.

### C1-regulated pneumococcal prophage activate in response to CSP

An alternative hypothesis was that C1-type regulators responded more sensitively, or rapidly, to competence induction than ImmAR-type regulators. This could explain the similarity in the deletion rates across all intact prophage as a consequence of C1-regulated prophage being activated equally in both cells deleting the prophage or acquiring rifampicin resistance, whereas ImmAR-regulated prophage were not activated by the end of the assay in either. Such elevated sensitivity of C1-type regulators would account for the differential deletion rates of impaired C1-regulated prophage. These mutated viruses may be too slow to lyse cells before being deleted by transformation when targeted by imported DNA (Fig. 2C). Nevertheless, these prophage could still lyse host cells that became competent, and acquired rifampicin resistance, if they were not targeted by HR themselves. Hence the deletion rate would rise because more cells survived when transformation removed the prophage, but the same fraction of bacteria that acquired rifampicin resistance would be lost. To test this hypothesis, the activation of the four prophage systems, including ϕRMV8*_immARcat_*, was studied through assaying gene expression (Fig. 3A) and prophage excision (Fig. 3B) after exposure to MMC or competence induction by CSP for two hours, corresponding to the period allowed for transformation-mediated deletion (Fig. 2E).

**Figure 3.**
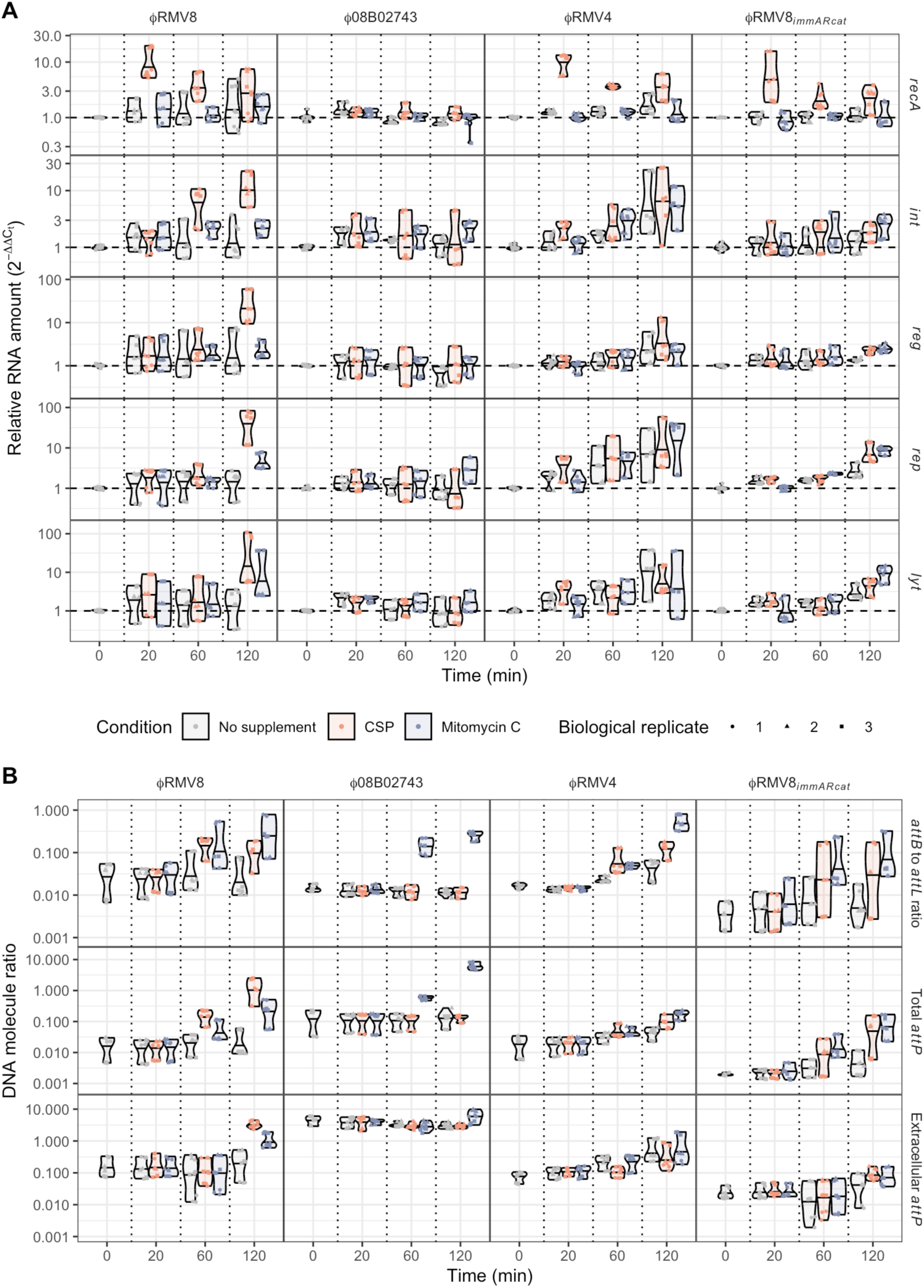
Activation of prophage in response to competence induction and exposure to mitomycin C. **(A)** Changes in RNA concentration measured by quantitative reverse transcriptase PCR. The expression of the *recA* gene and four prophage loci (*int*, *reg*, *rep* and *lyt*) were measured in each of the prophage systems during early exponential phase in rich media, and 20, 60 and 120 minutes after exposure to competence stimulating peptide (CSP), which induces competence, or mitomycin C (MMC). Expression was quantified as a change relative to the mean value at the pre-exposure timepoint, standardised to the expression of *rpoA*. Each plot corresponds to a particular combination of a prophage system, defined by the column, and a locus, defined by the row. Different colours distinguish the treatment types, and vertical dashed lines separate the different timepoints. For each combination of locus, prophage system, timepoint and condition, three technical measurements were made for each of three biological replicates, as indicated by the shapes of the points. Their distribution is summarised by a violin plot, which indicates the median value with a horizontal line. **(B)** Changes in DNA topology measured by quantitative PCR, using the primer pairs described in Fig. 2A. Data are shown as in panel (A), except that DNA concentrations were calculated as absolute ratios. The top row quantifies the excision of the prophage through comparing the ratio of the empty *attB* site, restored by prophage excision, to the *attL* site, formed by integrated prophage. The middle row shows the concentration of *attP* sites, formed by prophage excision and replication, in DNA extracted from pelleted cells. The bottom row shows the concentration of *attP* sites in DNA extracted from culture supernatant, representing virions released by host cell lysis.

In both the ϕRMV8 and ϕRMV8*_immARcat_* systems, which share the same host cell genotype, *recA* transcription was upregulated ∼5-10 fold 20 min post-CSP, but was not affected by MMC (Fig. 3A). Consistent with the C1-regulated prophage being most sensitive to competence induction, ϕRMV8 responded more strongly to CSP than MMC. Then the *int* gene was elevated 60 min post-CSP, followed by ∼10-100-fold upregulation of the replication and lysis genes by 120 min post-CSP (Fig. 3A). These kinetics are consistent with C1-regulated prophage evading deletion through activation. They contrasted with the response of ϕRMV8*_immARcat_*, in which there was a smaller increase in the expression of regulatory and replication genes 120 min post-CSP, but no rise in lytic gene expression relative to untreated cells. Yet both ϕRMV8 and ϕRMV8*_immARcat_* responded near-identically to MMC, with 2-10-fold rises in the expression of all prophage genes by 120 min post-stimulus (Fig. 3A). Hence only the C1-regulated ϕRMV8 was induced by CSP to express the genes necessary to lyse host cells, in keeping with the more efficient deletion of this prophage when these loci were disrupted by mutation (Fig. 2E).

The short-term dynamics of the ϕRMV4 system were comparable to those of ϕRMV8*_immARcat_*: although *recA* transcription rose following exposure to CSP, the prophage did not exhibit a substantial response to MMC or CSP. Instead, its activity rose over time as population density increased, suggesting the phage may be sensitive to a quorum sensing signal, which are commonly employed by pneumococci (Aggarwal et al. 2020), or nutrient availability. Either mechanism would also explain why ϕRMV4 mutations affected stationary phase cell density more strongly than the exponential growth rate (Fig. 2C). The least induction was observed in ϕ08B02743, in which host *recA* transcription did not rise following its exposure to CSP. However, when the prophage was deleted from the host cells, this response was restored (Fig. 4A). Similarly, CSP only induced upregulation of the late competence gene *comEA* when ϕ08B02743 was absent. Consequently, the prophage caused a ∼20-fold reduction in the transformation efficiency of its host (Fig. 4B). Therefore neither ϕ08B02743 or its host cell responded to CSP, as the prophage inhibited the induction of cellular competence genes.

**Figure 4.**
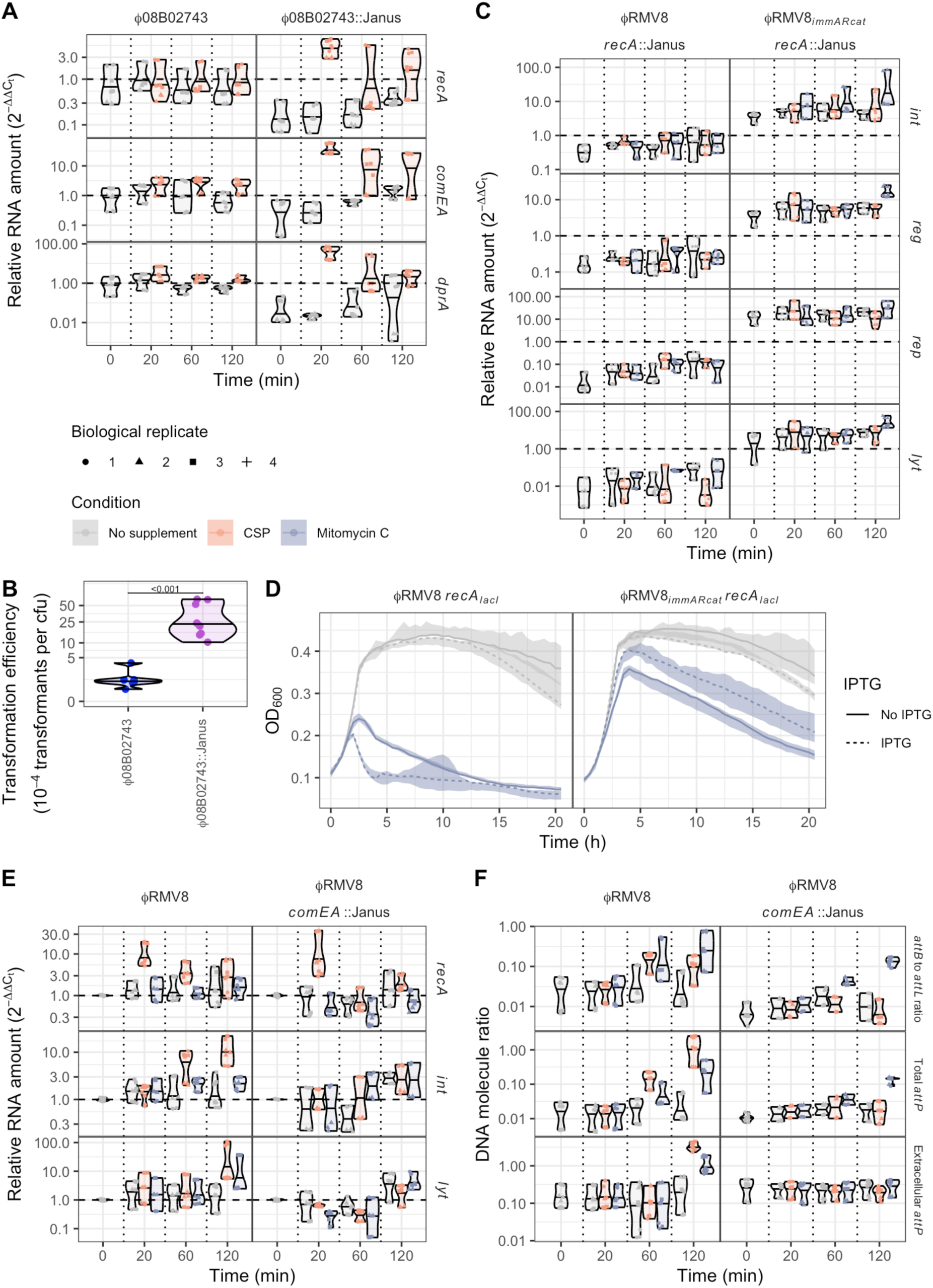
The interaction between prophage, RecA and ssDNA. **(A)** The effect of ϕ08B02743 deletion on competence gene expression. The effect of CSP on the expression of *recA*, *comEA* and *dprA* was assayed in the ϕ08B02743 system, and a derivative in which the prophage has been replaced by a Janus cassette. Data are shown as in Fig. 3A, except that all measurements are standardised to the initial timepoint obtained from the ϕ08B02743 system. **(B)** The effect of ϕ08B02743 deletion on the transformation efficiency of host cells. The ϕ08B02743 system, and a derivative in which the prophage has been replaced by a Janus cassette, were transformed with genomic DNA encoding an *rpoB* gene containing a base substitution conferring rifampicin resistance. The frequency of transformation was compared between the two genotypes using a two-tailed Wilcoxon rank sum test. The significance of the difference between them is indicated by the displayed *p* value. **(C)** Effect of RecA on prophage gene expression. The expression of the four prophage loci assayed in Fig. 3A were measured under equivalent conditions in *recA* ::Janus derivatives of the ϕRMV8 or ϕRMV8*_immARcat_* systems. The RNA levels were standardised relative to those in the equivalent prophage systems expressing *recA* under control of its native promoter at the initial timepoint. Therefore an expression level of one, indicated by the horizontal dashed line, indicates the pre-exposure level of expression in the wild type systems. **(D)** The effect of elevated RecA expression on prophage activation. The growth of bacteria with *recA* under the control of an IPTG-inducible *lacI* repressor binding site (*recA_lacI_*), and carrying either ϕRMV8 or ϕRMV8*_immARcat_*, is shown as in Fig. 2C. The colours distinguish cells grown in normal culture, and those exposed to mitomycin C. The dashed lines show the median OD_600_ for cells grown in the presence of the *recA* inducer IPTG, whereas the solid lines show the equivalent measurement from those grown without this supplement. **(E)** The effect of single-stranded DNA import by the competence pore ComEA on the expression of ϕRMV8 genes. The data for *comEA*^+^ wild type cells, shown in Fig. 3A, are compared to equivalent measurements in *comEA*::Janus mutants. **(F)** The effect of single-stranded DNA import by the competence pore ComEA on the excision of ϕRMV8. The data for *comEA*^+^ wild type cells, shown in Fig. 3B, are compared to equivalent measurements in *comEA*::Janus mutants.

Three sets of qPCR assays were used to understand how these gene expression differences related to prophage excision and activation (Fig. 3B). In addition to measuring excision using the *attB*-to-*attL* ratio, the extent of excised prophage replication was measured as the concentration of *attP* relative to chromosomal DNA, estimated using *rpoA* copy number. The same ratio was also measured in the culture supernatant, to quantify the release of virions relative to cell lysis, as it is not possible to use plaques to enumerate phage that infect encapsulated pneumococci (López and García 2004). Consistent with the gene expression data, ϕRMV8 exhibited the strongest response to CSP (Fig. 3B). This prophage excised from the chromosome in the first 60 min after the addition of CSP, before replicating, and lysing host cells by 120 min. MMC triggered a similar, albeit weaker, response. Hence competence is a natural stimulus for the activation of C1-regulated pneumococcal prophage.

The ImmAR-regulated prophage exhibited a different response to both stimuli. MMC induced all three to excise from the chromosome after 60 min, with the concentration of circular ϕ08B02743 rising ∼100-fold, despite none of them showing a strong transcriptional response to this stimulus at this timepoint. However, the absence of a substantial rise in the concentration of extracellular phage DNA implied there was limited cell lysis and virion release by 120 min. This would be consistent with increasing proportions of the ImmAR-regulated prophage populations becoming pseudolysogens in response to MMC. The response of both ϕRMV4 and ϕRMV8*_immARcat_* to CSP was similar to their behaviour after exposure to MMC, whereas ϕ08B02743 suppressed all responses to CSP (Fig. 3A). The divergence of the ϕRMV8*_immARcat_* response from that of ϕRMV8 supported the attribution of this behavioural change to the C1-type and ImmAR-type systems themselves. Therefore ImmAR-regulated prophage excised in response to MMC or CSP, but frequently stalled in a pseudolysogenic state.

### RecA can repress or activate prophage excision

The stronger response of ϕRMV8 to CSP, which increases *recA* expression, than MMC, which generates ssDNA lesions without stimulating *recA* transcription (Fig 3A), is likely attributable to its dependence on RecA. That the ImmAR-type regulators responded similarly to both stimuli suggested they were less dependent on RecA for activation. To test this difference, the induction of lysogens by MMC was assayed in *recA*^-^ cells (Fig. 2D, S21). The C1-regulated ϕRMV8 caused substantially less cell lysis in response to MMC in a *recA*^-^ mutant. However, the ImmAR-regulated prophage all lysed a high proportion of host cells in the absence of RecA. Although MMC appeared to have a diminished effect on the activation of ϕ08B02743, this was mainly a consequence of the *recA*^-^ mutant exhibiting a substantial growth defect even in unsupplemented media (Fig. S21). This suggested ImmAR-type regulators could function independently of, or potentially be repressed by, RecA.

To test for evidence of RecA-mediated repression, the *attB*-to-*attL* ratio was measured in wild type cells and *recA*::Janus mutants of all four prophage systems in the absence of any inducing stimuli, as excision from the genome was the most reproducible signal of ϕ08B02743 activity (Fig. 2B). Normal excision of the C1-type prophage ϕRMV8 required RecA, as did excision of the ImmAR-type prophage ϕRMV4, which was independent of SpaL. Yet the excision of both ϕ08B02743 and ϕRMV8*_immARcat_* increased when *recA* was disrupted. This repression by RecA was still evident in an *szaL*^-^ background, but excision of ϕ08B02743 was abrogated when *immA* was disrupted, implicating ImmA itself in this signalling pathway. Correspondingly, the hybrid prophage ϕRMV4*_immARcat_*, generated by transfer of the ϕ08B02743 lysogeny regulation locus into ϕRMV4, also increased its level of circularisation in a *recA*^-^ host. Therefore the activation of excision by ImmAR-type proteins exhibited differential sensitivity to RecA.

In all systems, the *attB*-to-*attL* ratio fell below 10^-3^ when the integrase was disrupted. However, all the *int*::*cat* mutants were able to lyse host cells in response to MMC at least as strongly as the intact prophage (Fig. 2D, S21), suggesting excision is not a prerequisite for efficient lytic gene expression. Quantitative PCR assays demonstrated this was not associated with *in situ* replication (Fig. S22). Significantly, the phage genes for lysis, rather than for lysogeny, were orientated with the coding bias of the pneumococcal genome at all 17 *attB* sites, facilitating their expression even while the prophage were still chromosomally integrated (Fig. S4-6). The separate regulation of excision and lysis added further complexity to the role of RecA in MMC induction of prophage, with ϕRMV8 being RecA-dependent, ϕ08B02743 activating through a RecA-independent mechanism, while RecA was necessary only for the excision of ϕRMV4 from the chromosome.

### RecA can repress or activate prophage gene expression

To understand the differential effects of RecA on C1-type and ImmAR-type regulators (Fig. 3A), the protein’s effects on ϕRMV8 and ϕRMV8*_immARcat_* were compared. Quantifying gene expression in *recA*::Janus mutants (Fig. S23), and standardising these to pre-exposure expression levels in *recA*^+^ cells (Fig. 4C), demonstrated that ϕRMV8 genes were repressed by 10-100-fold in *recA*^-^ cells, whereas ϕRMV8*_immARcat_* genes were ∼5-fold upregulated. Both CSP and MMC failed to activate ϕRMV8 *recA*::Janus, whereas ϕRMV8*_immARcat_ recA*::Janus still responded to MMC, suggesting activation was driven by ssDNA independently of RecA. Hence RecA’s repressive effect could be an indirect consequence of the protein competing for ssDNA-based structures that activated the prophage. Therefore, to test whether the effect of RecA changed with its concentration, the *recA* genes of ϕRMV8 and ϕRMV8*_immARcat_* were engineered to be under control of a *lacI* repressor sequence that was inducible by isopropyl β-D-1-thiogalactoside (IPTG; Fig. S24). While IPTG caused a small decrease in cell growth in unsupplemented media for both prophage systems, it enhanced the effects of MMC on ϕRMV8, while limiting the induction of ϕRMV8*_immARcat_* (Fig. 4D). This effect of IPTG was dose-dependent and not observed when *recA* was expressed from its native promoter (Fig. S25). Hence replacing a C1-type regulator with an ImmAR-type regulator reversed the sensitivity of a prophage to RecA concentration.

The synergistic activation of ϕRMV8 by RecA overexpression and MMC were consistent with RecA-ssDNA nucleoprotein filaments being the trigger for prophage activation. This suggested the activation of C1-type regulator by CSP may by driven by the import of ssDNA. Therefore ϕRMV8 gene expression was quantified in *comEA*::Janus mutants, which lack a functional competence pore (Fig. 4E). Despite *recA* still being upregulated after CSP addition, the ϕRMV8 genes failed to respond to the induction of competence in the *comEA*::Janus mutant. Assaying the excision and replication of the prophage confirmed induction by CSP depended on the import of DNA, whereas the response to MMC-induced ssDNA lesions did not (Fig. 4F, S26).

### ImmAR-regulated prophage adopt divergent HR-sensitive activation strategies

The contrasting repression of ϕ08B02743’s ImmAR-type regulators by RecA (Fig. 2B, 4C) suggested these proteins may be activated by ssDNA alone, as for the ImmA orthologue PprI (Lu et al. 2024), or by the association of a different protein with ssDNA. Hence the effects of deleting the genes for other HR-associated DNA-binding proteins on the ability of exogenous stimuli to induce ϕ08B02743 were assayed (Fig. S27-S28). The most substantial reduction in cellular growth was observed after adding MMC or CSP to a *dprA*::Janus mutant culture (Fig. 5A). This effect depended upon the presence of ϕ08B02743 (Fig. S27-S28), indicating it was a consequence of phage-driven cell lysis. This result was counterintuitive, as DprA represses induction of the competence regulon (Mirouze et al. 2013), which includes *recA*. In keeping with this regulatory activity, disruption of *dprA* facilitated the CSP-induced upregulation of both *comEA* and *recA* in 08B02743 to a similar extent as deletion of the ϕ08B02743 prophage (Fig. 5B). Correspondingly, prior to exposure to CSP, *dprA* transcription was reduced >5-fold when ϕ08B02743 was deleted from cells (Fig. 4A). Hence the prophage-mediated suppression of competence induction appeared to be achieved through constitutively elevating DprA levels. In the absence of DprA, the induction of competence activated ϕ08B02743 to lyse host cells in a similar manner to ϕRMV8, despite the absence of a C1-type regulator (Fig. 5A).

**Figure 5.**
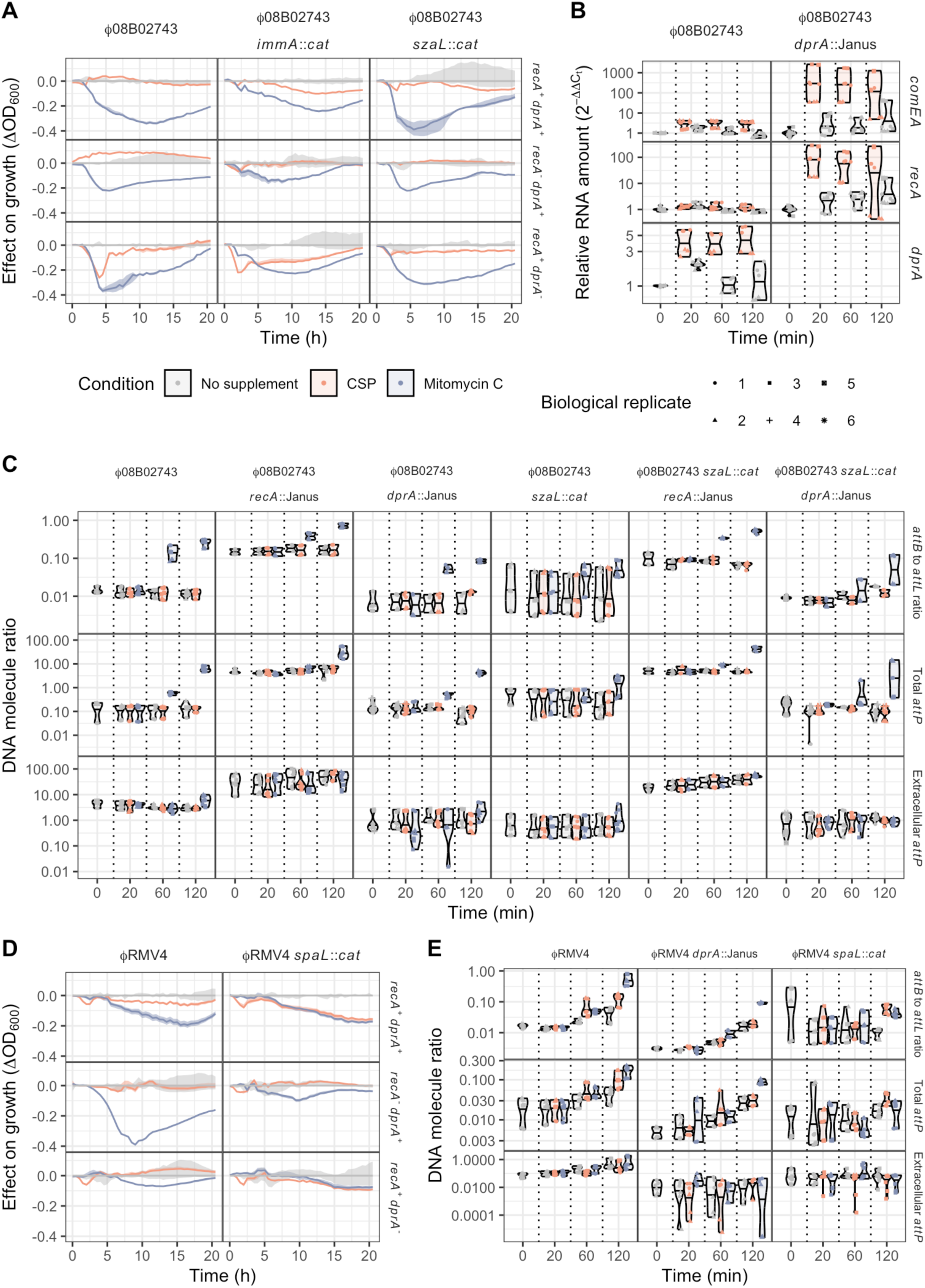
Effect of DprA and population density on prophage activation. **(A)** Effect of MMC and CSP on the growth of ϕ08B02743, and derivatives in which the *immA* or *szaL* lysogeny module genes were disrupted. Each line shows the deviation from the median growth of the genotype in unsupplemented media, as in Fig. 2D. The ribbons show the range in measurements across replicates. The graphs are separated into columns, corresponding to the prophage genotype, and rows, according to whether the *recA* or *dprA* host cell ssDNA-binding proteins were intact. **(B)** The effect of MMC and CSP on the expression of competence-associated genes in 08B02743, and a mutant in which *dprA* was disrupted. Data are shown as in Fig. 3A. **(C)** The effect of prophage regulators and host cell ssDNA-binding proteins on prophage excision and activation. The three rows quantify the excision, replication and release of the prophage, as in Fig. 3B. Each column corresponds to a different combination of mutations in the prophage-encoded regulators and host-encoded ssDNA-binding proteins. **(D)** Effect of MMC and CSP on the growth of ϕRMV4, and a derivative in which the *spaL* lysogeny module gene was disrupted. Data are shown as in Fig. 2D. **(E)** The effect of prophage regulators and DprA on prophage activation. The three rows quantify the excision, replication and release of the prophage, as in Fig. 3B.

To understand which part of the prophage regulatory machinery drove this response, both the *immA* and *szaL* lysogeny cluster genes (Fig. 1E) were disrupted in backgrounds lacking the *recA* and *dprA* genes (Fig. 5A). While the disruption of *immA* made ϕ08B02743 less responsive to MMC, in addition to reducing prophage activation in unsupplemented media (Fig. S29), the *immA*^-^ *dprA*^-^ double mutant retained the increased sensitivity to CSP associated with the *dprA*^-^ single mutant. These observations supported the existence of an ImmAR-independent activation mechanism (Fig. 5A), implicating the ϕ08B02743-specific SzaL protein. Such a pathway would be consistent with neither ϕRMV8 nor ϕRMV4 being induced at an elevated rate following disruption of *dprA* (Fig. S27-S28). An *szaL*^-^ single mutant replicated the MMC hypersensitivity of the *dprA*^-^ mutant, but was less susceptible to induction by CSP (Fig. 5A, S29). Disrupting *dprA* in the *szaL*^-^ background did not restore the CSP sensitivity observed in the *dprA^-^* single mutant (Fig. 5A). This was consistent with SzaL triggering prophage activation in response to the competence induction enabled by the absence of DprA. Despite SzaL mediating this DprA-dependent activation, disruption of *szaL* did not restore the induction of *recA* transcription by CSP (Fig. S30), implying the upregulation of *dprA* was driven by a distinct activity of the prophage. Hence SzaL functions separately from ImmAR to sensitise ϕ08B02743 to the induction of competence.

The activation of ϕ08B02743 by CSP peaked after ∼4 h (Fig. 5A), which likely reflects the competent state being atypically prolonged in the absence of shutdown by DprA, as indicated by the slow decline in late competence gene expression observed in the *dprA*::Janus mutant (Fig. 5B). Yet over the shorter period relevant to deletion by transformation, SzaL did not have a substantial effect on phage gene expression (Fig. S30). The responses of ϕ08B02743 to external stimuli were more rapidly detected through assays of the prophage’s topological state (Fig. 3B). Therefore analogous tests were used to measure how *recA*^-^ and *dprA*^-^ mutants affected the response to MMC and CSP (Fig. 5C). Neither DprA nor SzaL had a substantial effect on ϕ08B02743 excision or replication, although both processes were detectably higher in the presence of MMC and the absence of RecA. However, the *dprA*^-^ and *szaL*^-^ mutations were associated with a decreased ratio of prophage to chromosomal DNA in the culture supernatant. The double mutant exhibited the same phenotype as the single mutants, again consistent with their effects being mediated through a common pathway. Given the altered patterns of cell lysis in the *szaL*^-^ and *dprA*^-^ mutants (Fig. 5A), this suggested SzaL moderated the size and timing of the burst of virions generated by intracellular replication in response to DprA-dependent signals.

The equivalent approach was used to test whether DprA and SpaL formed an analogous signalling pathway in ϕRMV4 (Fig. 5D, S29). The *spaL*^-^ mutation increased the activation of the prophage by CSP. However, rather than the transient induction observed in ϕ08B02743, this activation persisted as cell density rose. Correspondingly, although *spaL* disruption had little effect on prophage gene expression (Fig. S30), loss of this gene eliminated the apparent cell density-dependence of ϕRMV4 excision, replication and lysis (Fig. 5E). While the *spaL*^-^ mutation had a substantial effect on ϕRMV4 activation in a *recA*^-^ background, there was little evidence of an interaction with DprA, the loss of which almost abrogated cell lysis by ϕRMV4 (Fig. 5D). Correspondingly, the *dprA*^-^ mutant still underwent population density-dependent excision and replication, but the near lack of lysis meant viral DNA was difficult to detect in the supernatant (Fig. 5E). Therefore these similar ImmAR-regulated prophage employed divergent activation strategies that integrated information from co-regulators and the HR protein DprA, in addition to RecA, to alter the rates of pseudolysogeny and virion release.

### ImmAR-type regulators require longer signal duration to cause cell lysis

The combination of signals required to release virions of the ImmAR-regulated prophage suggested these viruses would be slower to commit to cell lysis than the C1-regulated ϕRMV8. Nevertheless, both types of prophage lysed cells with similar kinetics when continuously exposed to MMC (Fig. 2D). To test whether the time taken to decide to lyse the host cell differed between the four prophage systems, each was exposed to MMC pulses of 20 min (equivalent to the period for which the competence machinery is induced), 60 min (the timepoint at which prophage were observed to have excised from the chromosome), and 120 min (the timepoint at which cells were lysed by ϕRMV8). The washed cells were then transferred into in unsupplemented media (Fig. 6A, Fig. S31).

**Figure 6.**
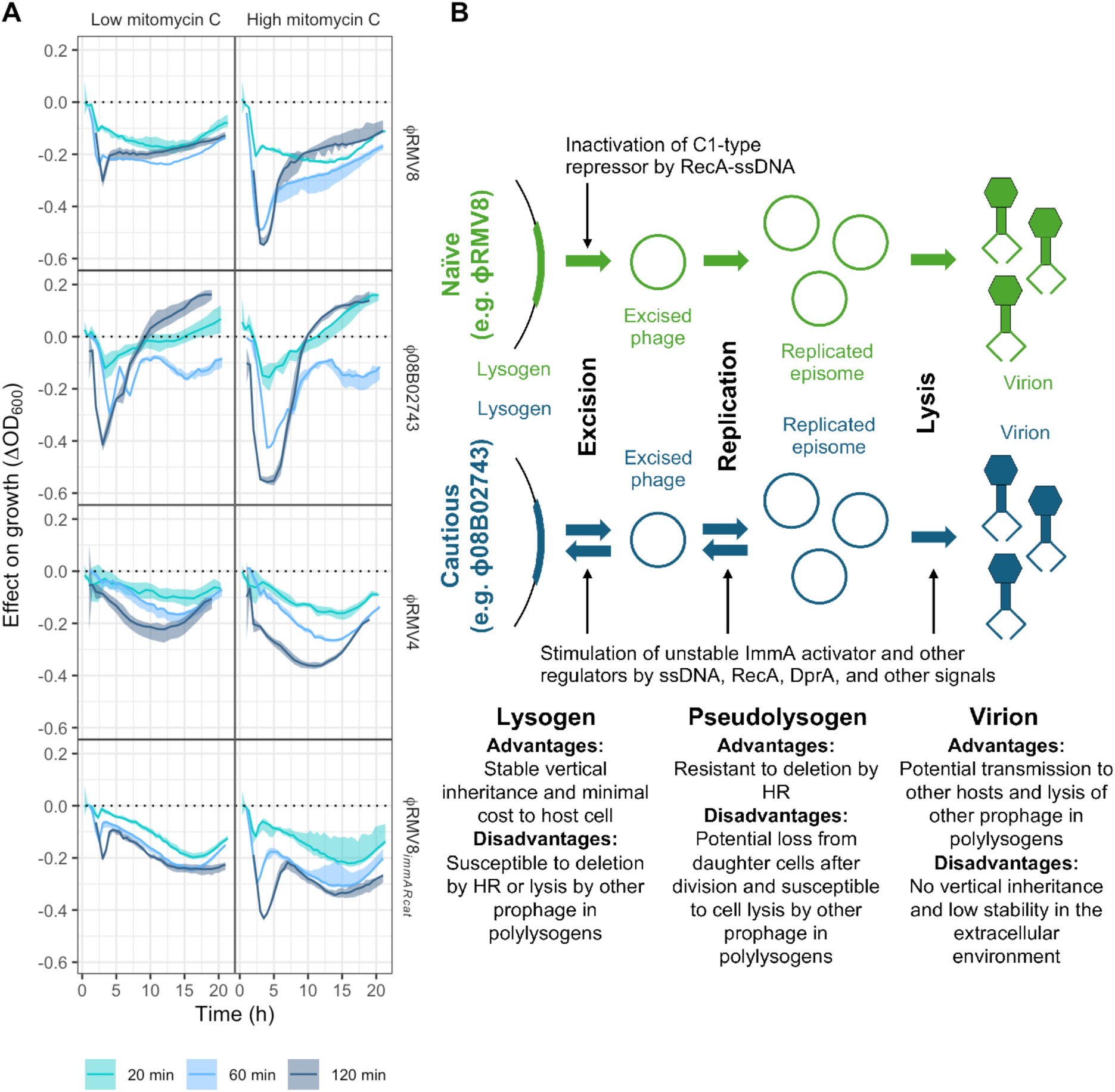
Contrasting prophage strategies in the response to inducing stimuli. **(A)** The effect of mitomycin C pulses of different durations on prophage activation. Each prophage system was exposed to a 20, 60 or 120 minute pulse of mitomycin C, either at a low (5 ng μl^-1^) or high (10 ng μl^-1^) concentration, prior to being transferred to fresh unsupplemented media. The line plots show the effect of the pulse on the subsequent growth of cells, with the colour corresponding to different pulse durations. The solid line shows the difference between the median post-pulse OD_600_ and the median OD_600_ of untreated cultures, grown in parallel. The shaded area shows the range of OD_600_ values observed in post-pulse cultures, similarly adjusted by subtracting the median OD_600_ of untreated cultures. (B) A summary of the differences between the “naïve” C1-regulated ϕRMV8 prophage, and the “cautious” ImmAR-regulated ϕ08B02743 and ϕRMV4 prophage.

These experiments demonstrated that ϕRMV8 committed to lysing host cells between 20 and 60 min, with the additional 60 min MMC exposure causing little additional activation. However, the three ImmAR-regulated prophage increasingly committed to host cell lysis throughout the 120 min. This effect was observed at two different MMC concentrations, and was not an artefact of ϕRMV8 being more sensitive to continuous exposure to MMC (Fig. S32). Therefore the C1-type regulators and ImmAR-type regulators prophage are associated with two different strategies for controlling the activation of phage (Fig. 6B).

## Discussion

This study demonstrates that pneumococcal transformation is a highly efficient mechanism for the deletion of prophage (Fig. 2E). The was despite the three characterised natural prophage each exhibiting distinct strategies for evading deletion. The C1-regulated ϕRMV8 activated its lytic cycle in response to a short pulse of MMC (Fig. 6A), or the transient induction of competence by CSP (Fig. 3). As only a small proportion of pneumococci both become competent after CSP administration (Veening et al. 2008; Kwun et al. 2023) and import DNA, the detected magnitude of phage induction suggests a high sensitivity to RecA-ssDNA filaments. Phage were excised from the genome after 60 min, and had activated transcription, replicated and lysed their host cells by 120 min. Such lysogens that activated in competent cells were described as “naïve” in *Bacillus subtilis* (Fig. 6B), as their induction was characterised as a maladaptive response to the competence-associated triggering of the SOS response (McVeigh and Yasbin 1996). However, the high sensitivity of the C1-type regulator to RecA-ssDNA filaments can instead be understood as an adaptation that enables prophage to escape HR-mediated elimination, which affects vertical transmission through inheritance, by rapidly transitioning to a lifecycle that enables horizontal transmission (Kwun et al. 2019). The speed of this response appears crucial for the prophage to evade deletion by HR, suggesting a finely-balanced race between MGEs and the cellular recombination machinery. Yet the RecA-ssDNA filaments appear late in the transformation process (Johnston et al. 2014). Hence it is likely the C1-type repressors recognise these structures because they are highly-conserved as the point of convergence (Rocha et al. 2005) between the otherwise variable competence machinery, found across many bacteria (Johnston et al. 2014) and archaea (Fonseca et al. 2020), as well as other HR mechanisms (Lang et al. 2012; Chen et al. 2018) that could drive prophage deletion.

By contrast, the ImmAR-regulated prophage exhibited a “cautious” strategy (Fig. 6B). After 60 min, both ϕRMV4 and ϕRMV8*_immARcat_* excised from the genome in response to CSP or MMC (Fig. 3B). The magnitudes of these responses were similar at a population level, which suggests frequent, rapid excision of prophage in the subset of cells that induce competence. Although ϕ08B02743 only responded to MMC, this is likely a consequence of its own strategy for evading deletion by HR, which involves inhibiting transformation by elevating expression of the competence repressor DprA (Fig. 5B). However, none of the ImmAR-regulated prophage substantially upregulated any viral genes by 120 min (Fig. 3A). This is consistent with the longer MMC pulse required to induce these viruses (Fig. 6A), suggesting the excised pseudolysogens may reintegrate once the activating stimulus is removed. Temporarily becoming a pseudolysogen, to evade a time-limited risk of removal by HR, before reinserting into the chromosome to ensure stable inheritance after cell division is likely to be an effective means of maximising vertical transmission in a transiently competent host (Fig. 6B). This is consistent with similarly reversible MMC-induced excision of phage-related chromosomal islands, which are persistently associated with pneumococcal genotypes in population genomic datasets (Lim et al. 2025). Such behaviour is difficult to rationalise in the absence of the risk of deletion by HR. Furthermore, the advantage is private to the excised MGE itself, whereas inhibiting cellular competence induction or functioning also benefits competitor MGEs within the same host cell.

Therefore both the naïve and cautious strategies likely evolved under selection to evade chromosomal curing. However, the co-existence of the two strategies suggests each is optimal in different situations. One possible explanation is that cautious prophage are likely to cause less of a fitness cost to their host, but risk not transmitting from host cells that also contain naïve prophage, which are faster to cause cell lysis. Hence the cautious prophage would be more successful in monolysogens (∼35% of pneumococci) than polylysogens (∼40% of pneumococci; Fig. 1B). This is consistent with the observation that there is a negative relationship between the number of phage regulators found in an isolate, and the proportion of those regulators of the ImmAR-type (Fig. S2). Furthermore, within the cautious phage, there were differences in the effect of both RecA on excision (Fig. 2B) and DprA on lysis (Fig. 5A,D), consistent with differing sensitivity to ssDNA-based structures. Such divergence provides additional opportunities for competition between prophage in polylysogenic hosts. Correspondingly, the suppression of competence induction by ϕ08B02743 not only reduces its deletion by HR, but also interferes with the regulation of other prophage within the same host, without affecting the RecA-independent activation of ϕ08B02743 itself (Fig. 3B, Fig. 5A). Therefore the multiple pressures shaping phage evolution can explain the co-existence of these distinct strategies between competing viruses (Fig. 6B).

The detailed characterisation of representative prophage in this analysis precludes the sample size necessary to determine whether the naïve and cautious strategies always correlate with the C1-type and ImmAR-type regulators, respectively. Nevertheless, the cautious ϕRMV8*_immARcat_* was generated from the naïve ϕRMV8 simply by switching these systems (Fig. 2E, 3, 4B). This is consistent with the change depending on a single locus. These differences are also explicable through the biochemistry of the regulators: while *E. coli* phage λ C1 is considered to be an irreversible switch (Ptashne et al. 1982), the ImmA-type protease activator of at least one streptococcal phage is less stable than its paired ImmR repressor (Koberg et al. 2015), suggesting that after an inducing stimulus is removed, an increased ratio of ImmR to ImmA may reverse the induction of the phage. Furthermore, the phage regulatory proteins’ contrasting response to inducing signals can also explain their differential distribution between *attB* sites (Fig. 1B). C1-type regulators dominate at *attB* sites where prophage inhibit transformation through causing a loss-of-function disruption to a protein-coding gene (Denapaite et al. 2010; Croucher et al. 2011) (e.g. *attB_comYC_*, *attB_ssbB_*). This may be because even transient restoration of a protein-coding gene by reversible excision would likely enable sufficient production of competence machinery components to enable transformation at the time at which the prophage reinserted.

The ImmAR-regulated prophage are associated with strategies that cause a smaller reduction in host cell competence: the ∼10-fold reduction in host cell transformation efficiency associated with csRNA3 modification (Kwun et al. 2022), or the ∼20-fold reduction caused by ϕ08B02743 elevating DprA levels. By contrast, HR is effectively shut down in pneumococcal lineages in which prophage disrupt *comYC* (N. Croucher, Chewapreecha, et al. 2014) or transposons disrupt *comEC* (D’Aeth et al. 2021). Similarly, the import of the MGE-encoded RocRp RNA in *Legionella* species, representing a modified form of the competence-regulating cell-encoded RocR RNA, causes a >100-fold reduction in transformation efficiency (Durieux et al. 2019). Nevertheless, the relatively small effects are still advantageous to the MGEs: while a 10-fold reduction in host transformation will prevent ∼90% of MGE deletions, a 100-fold reduction will only prevent an additional ∼9% of removals through HR. This advantage of partial inhibition of host transformation has been experimentally verified using genomic islands in *Acinetobacter baumannii* (Tuffet et al. 2024). Such benefits to MGEs are likely to underlie the extensive variation observed in transformation efficiencies across species, even between closely-related isolates (Mazzamurro et al. 2024).

These divergent strategies across prophage are further complicated by the complex regulation of pneumococcal competence (Kwun et al. 2023), which has likely emerged through the co-evolutionary arms race with MGEs. The highly-conserved DprA protein (Claverys et al. 2009; Johnston et al. 2014) appears to be a key interface in this interaction, consistent with related proteins having anti-phage functions (Burroughs et al. 2015). By coating ssDNA imported by the competence machinery (Claverys et al. 2009), DprA delays the formation of RecA-ssDNA filaments that activate C1-regulated prophages, while also shortening the duration of competence (Mirouze et al. 2013), potentially limiting the induction of slowly-activating ImmAR-regulated prophages. Additionally, it is likely that both the rapid shutdown of competence (Mirouze et al. 2013), and the “bet hedging” induction of this state in only a subset of pneumococcal populations (Veening et al. 2008; Kwun et al. 2023; Prudhomme et al. 2024), reflects the trade-offs between the benefits of chromosomal curing of MGEs by HR, and potential for activating prophage.

Hence MGEs adopt a combination of strategies to evade deletion by HR. They limit the induction of competence through their targeting of *attB* sites and encoding functions that interfere with host cell expression patterns. If this fails, they can excise from the genome in response to the import of single-stranded DNA into the cell. Finally, they can specifically lyse transformable cells, selecting for genotypes that hedge their bets against the risks of inducing competence. This has substantial implications for pneumococcal epidemiology, as transformation is required for the acquisition of resistance (Dowson et al. 1993; D’Aeth et al. 2021) and evasion of vaccine-induced immunity (D’Aeth et al. 2021). Such impacts of the co-evolutionary arms race between phage and RecA-mediated HR will be ubiquitous across prokaryotes, given the broad distribution of MMC-sensitive prophage across microbiomes (Williamson et al. 2007; Paul 2008; Engelhardt et al. 2011; Maurice et al. 2011; Minot et al. 2011; Kim and Bae 2018; Roy et al. 2020), and the plethora of cellular DNA exchange mechanisms that depend on HR, which include the mobilisation of DNA through gene transfer agents (Lang et al. 2012), extracellular vesicles (Berleman and Auer 2013), and virions facilitating generalised, specialised and lateral transduction (Humphrey et al. 2021). Its impact may be even more extreme in other species, as the competence machinery is broadly conserved (Johnston et al. 2014), yet often difficult to induce *in vitro*, suggesting a strong pressure to suppress its activation. Hence as both MMC-responsive prophage (Schleper et al. 1992; Prangishvili et al. 2006; Götz et al. 2007; Fusco et al. 2020) and competence, let alone cellular HR (Lang et al. 2012; Johnston et al. 2014; Chen et al. 2018; Fonseca et al. 2020), are more taxonomically broadly distributed than the SOS response (Erill et al. 2007), it is likely that deletion of integrated genomic parasites was the original selection pressure for the evolution of RecA- and ssDNA-sensitive regulators such as C1 and ImmAR. Hence this ancient global conflict between MGEs and their hosts has determined, and will continue to shape, the evolution and epidemiology of many bacterial pathogens.

## Methods

### Bacterial isolates and sequence data

All pneumococcal genotypes used in this study were derived from the previously-characterised *S. pneumoniae* RMV8 (Kwun et al. 2018; Kwun et al. 2022); *S. pneumoniae* 08B02743 (Croucher et al. 2016); *S. pneumoniae* RMV4 (Kwun et al. 2018); *S. pneumoniae* 11930 (Croucher et al. 2011), or *S. pneumoniae* R6 (Lanie et al. 2007). The parental genotypes, and accession codes for associated sequence data, are described in Table S1. All mutant genotypes used in this work are listed in Table S3. All experimentally-characterised pneumococci lacked the phase-variable *ivr* locus (Manso et al. 2014; Kwun et al. 2018; Croucher et al. 2024), encoding a restriction-modification system that affects prophage deletion (Text S2, Fig. S9). Similarly, the *tvr* locus was either deleted, or its arrangement was locked through disruption of *tvrR*. These modifications used the primers detailed in Table S4.

### Bioinformatic and statistical analysis of prophage sequences

The bioinformatic analyses of prophage sequences are detailed in Text S1. In summary, prophage were identified through a BLASTN (Camacho et al. 2009) comparison of 20,047 Global Pneumococcal Sequencing (GPS) project isolates (Gladstone et al. 2019) with a manually-curated database of 63 known prophage sequences. This enabled the identification of 16 *attB* sites (Table S2). A further BLASTN search identified genomes in which these *attB* sites were disrupted.

An independent analysis of prophage sequences identified in a smaller dataset (Croucher, Coupland, et al. 2014) enabled the identification of C1-type and ImmA-type regulatory proteins. These were used to generate hidden Markov models using MAFFT (Katoh and Standley 2013) and HMMer3 (Eddy 2011), available from FigShare, which were then used to classify proteins encoded by the GPS isolates (see Text S1).

Plots and statistical analyses were undertaken in R (R Core Team 2019) using the tidyverse (Wickham et al. 2019), cowplot (Wilke 2020), ggpubr (Kassambara 2020), ggtree (Yu et al. 2017), genoPlotR (Guy et al. 2010), circlize (Gu et al. 2014) and fitdistrplus (Delignette-Muller and Dutang 2015) libraries.

### Culturing and growth assays

*S. pneumoniae* were grown at 35°C in 5% CO_2_. Culturing on solid media used Todd-Hewitt broth supplemented with 0.5% yeast extract (Millipore) and 1.5% agar (Sigma-Aldrich). Culturing in liquid media used a 2:3 ratio mixture of Todd-Hewitt broth (Sigma-Aldrich) with 0.5% yeast extract (Sigma-Aldrich) and Brain-Heart Infusion media (Sigma-Aldrich), as described previously (Kwun et al. 2022). For growth assays, 2x10^4^ cells from titrated frozen stocks stored in 10% glycerol (Sigma-Aldrich) were used to inoculate 200 μl liquid media in 96-well microtiter plates at 35°C in a 5% CO_2_ atmosphere. The optical density at a wavelength of 600 nm (OD_600_) was measured at 30 min intervals using a FLUOstar Omega microplate reader (BMG LABTECH). At least three independent biological replicates of each genotype were grown in separate wells for each analysis.

For measuring prophage induction by different stimuli, pneumococcal cells were first grown in 10 ml of mixed media at 35°C in 5% CO_2_ overnight. A 750 μl sample was used to inoculate a fresh 10 ml liquid culture, which was maintained under the same conditions until it reached an OD_600_ of 0.2, measured using an Ultrospec 10 photospectrometer (Amersham Biosciences). Two hundred microlitre samples of this early exponential phase culture were then transferred into individual wells of 96-well microtitre plates. Individual wells were then supplemented with stimuli with the potential to induce prophage: 0.75 μl 0.1 mg ml^-1^ CSP1, or 0.5 mg ml^-1^ CSP2 (AnaSpec; Table S1); 1 μl 500 mM calcium chloride (Sigma-Aldrich); 2 μl 0.5 μg ml^-1^ mitomycin C (Cayman Chemical Company); 2 μl 9% aqueous hydrogen peroxide solution (LCM Group); 2 μl 0.02 M ZnCl_2_ (Sigma-Aldrich); 5 μl 0.02 M sodium citrate tribasic dihydrate (Sigma-Aldrich), or 5 μl 0.2 M manganese (II) chloride tetrahydrate (Sigma-Aldrich). The optical density of each well was then measured as described above.

### Generation and sequencing of DNA constructs

To extract intracellular DNA, cells were pelleted by centrifugation for 10 min at 4000 *g*, followed by resuspension in 480 μL lysis buffer (Promega) and 120 μL 30 mg mL^−1^ lysozyme (Promega). The mixture was incubated at 35 °C for 30 min. Genomic DNA was then extracted from cells using the Wizard Genomic DNA purification kit (Promega). Whole genome sequences were generated using a PromethION system (Oxford Nanopore Technologies; Table S1).

To quantify the concentration of phage DNA released into the medium from lysed cells, a 2 ml culture sample was spun down at 13,000 *g*, at 10 °C, for 10 min. The supernatants were then filtered through 0.22 μm cellulose filters (Sartorius) and stored at -80 °C. The frozen supernatants were thawed and diluted in a 1:1 ratio with water, then incubated at 95 °C for 10 min. This freezing, thawing and heating cycle was repeated once more, to ensure release of phage DNA from virions. This processed supernatant was then used directly as the template for subsequent experiments.

PCR amplification used either DreamTaq (Sigma-Aldrich), for establishing the size of amplicons, or Platinum PCR SuperMix High Fidelity (Invitrogen), for generating constructs for mutagenesis or sequencing. The oligonucleotides (Invitrogen) used to prime PCR amplifications are listed in Table S4. For the construction of mutants, the upstream (or left) flanking region was amplified with oligonucleotides that added an *Apa*I restriction site at the 5’ end of the right primer. The downstream (or right) flanking region was amplified with oligonucleotides that added a *Bam*HI restriction site at the 5’ end of the left primer. The appropriate resistance marker (a Janus cassette, or an *aph3’*, *cat* or *ermB* gene) was amplified using primers that carried an *Apa*I or *Bam*HI site at the 5’ end of the left and right primers, respectively.

Individual DNA constructs were extracted from agarose gels after electrophoresis using the QIAquick PCR & Gel Cleanup kit (Qiagen). DNA concentrations were quantified using dsDNA BR Assay Kit for the Qubit (Qiagen). PCR amplicons were digested using the appropriate restriction enzymes (Promega), according to the manufacturer’s instructions. The digested DNA molecules were ligated using T4 DNA ligase (Sigma-Aldrich) at room temperature overnight, and the full construct amplified by PCR. Each DNA amplicon was checked through agarose gel electrophoresis or dideoxy terminator sequencing (Genewiz), depending on its final use.

### Transformation and mutagenesis

Competence was induced by addition of the appropriate CSP pherotype (Table S1) at early exponential phase, defined as an OD_600_ of 0.2-0.25. A 1 mL sample of the culture was mixed with 5 μl 500 mM calcium chloride and the transforming DNA (1 μg for genomic DNA, or 100 ng for a PCR construct). After a 2 h incubation, cells were plated on the appropriate selective media. To select transformed cells, solid media were supplemented with the appropriate antibiotic. For the Janus cassette or an *aph3*’ marker, kanamycin (Sigma-Aldrich) at 400μg ml^-1^ was used. For the *ermB* marker, erythromycin (Sigma-Aldrich) at 1μg ml^-1^ was used. For the *cat* marker, chloramphenicol (Sigma-Aldrich) at 4 μg ml^-1^ was used. For the alteration of *rpoB*, rifampicin (Fisher Scientific) at 4 μg ml^-1^ was used.

For the assays of prophage deletion, donor genotypes were constructed by selecting for the replacement of a prophage by an appropriate resistance marker, then adding a rifampicin resistance base substitution. The recipient cells were transformed with 1 μg of the genomic DNA. The transformants that had acquired resistance markers were selected on agar plates supplemented with either rifampicin or kanamycin. For the assay of prophage insertion, dual selection with both kanamycin and chloramphenicol was used to ensure the full-length ϕRMV8*_immARcat_ int*::Janus prophage had been acquired by the recipient.

### Extraction of RNA for gene expression quantification

For each biological replicate, cells were grown in 10 ml of media overnight, and a 750 μl sample was used to inoculate six fresh 10 ml cultures grown in parallel. At an OD_600_ of 0.2, a total of 4 ml was collected across all cultures to represent the pre-treatment timepoint. Two cultures were maintained as untreated controls; two were supplemented with 25 μl CSP (1 mg ml^-1^ CSP1, or 0.5 mg ml^-1^ CSP2; Table S1), and two were supplemented with 100 μl mitomycin C. At 20, 60 and 120 min post-treatment, 4 ml of cell culture was collected to represent each condition. All samples were immediately treated with RNA protect in a 1:1 ratio by volume. RNA was then extracted using the SV Total RNA Isolation System (Promega), according to the manufacturer’s recommendations.

### Nucleic acid analysis with quantitative real-time PCR

The concentration and quality of all extracted RNA samples was assessed using a Nanodrop ND-1000 spectrophotometer (Thermo Fisher Scientific). A 0.2 μg sample of RNA was then treated with amplification-grade DNase I (Invitrogen) and used to template a reverse transcription reaction using the First-Strand III cDNA synthesis kit (Invitrogen). Each reaction used 100 units of the SuperScript III reverse transcriptase, 1 μL of 100 μM random hexamer primers (Thermo Fisher Scientific) and 1 μL of 10 mM dNTP mix (Bioline). This mixture was incubated at 25 °C for 5 min to enable annealing; then 55 °C for a further 30 min for DNA synthesis, prior to reverse transcriptase inactivation through heating at 70 °C for 15 min.

Quantitative real-time PCR amplifications were run using MicroAmp Optical 96-well reaction plates (Thermo Fisher Scientific) and the QuantStudio 7 Flex Real-Time PCR System (Applied Biosystems). Samples were denatured at 50 °C for 2 min, prior to 40 amplification cycles of denaturation at 95 °C for 15 s, followed by annealing and elongation at 60 °C for 1 min. Validation of the results used melt curve analysis of each amplicon on each plate. RNA quantification measured transcript concentration relative to the detected level of *rpoA*. DNA quantification converted levels detected by qPCR to absolute concentrations using standard curves generated for each amplicon of interest. For each analysis, three technical replicate measures were made of three or more biological replicates, unless otherwise specified. All RNA expression levels were standardised relative to the levels of *rpoA*.

## Data Availability

The raw data are available from a Github repository (https://github.com/nickjcroucher/prophage_transformation_analysis). The hidden Markov models used for identifying C1-type and ImmAR-type regulator proteins are available from FigShare (https://doi.org/10.6084/m9.figshare.26425075). The sequence data used in this study are available from the European Nucleotide Archive with the accession codes listed in Table S1, with additional information available from FigShare (https://doi.org/10.6084/m9.figshare.26425042 and https://doi.org/10.6084/m9.figshare.26425066).

## Code Availability

The code used in this analysis is available from https://github.com/nickjcroucher/prophage_transformation_analysis.

## Acknowledgements

This work was supported by a Sir Henry Dale fellowship jointly funded by Wellcome and the Royal Society (grant 104169/Z/14/A), and the UK Medical Research Council and Department for International Development (grant MR/R015600/1).

