## Supplementary materials for "Chromosomal curing drives an arms race between bacterial transformation and prophage"

Text S1-S2

Figures S1-S32

Tables S1-S4

### Text S1: Bioinformatic analysis of prophage sequences

#### Identification of prophage sequences

The draft assemblies of the 20,047 isolates analysed by the Global Pneumococcal Sequencing (GPS) project (Gladstone et al. 2019) were searched with a manually-curated set of 63 prophage sequences using BLASTN (Camacho et al. 2009) v2.6.0 with default settings. The distribution of sequences of prophage origin was assessed by identifying the maximum alignment length between the full set of representative prophages and each individual pneumococcal isolate. Using the population-wide distribution, an empirically-determined criterion of a maximum alignment length greater than 5 kb was used to identify pneumococci likely to contain a prophage (Fig. S2). This suggested 12,194 pneumococci (60.8%) contained a prophage. However, this is likely to be an underestimate, as it omits those prophage that do not exhibit detectable similarity to those used to search the isolate assemblies.

#### Identification of *att* sites

The positions at which representative prophage inserted into the pneumococcal genome were determined through pairwise comparisons of lysogenic GPS isolates with *S. pneumoniae* R6 (Hoskins et al. 2001) (accession code AE007317) using the Artemis Comparison Tool (Carver et al. 2008) (Fig. S5-6). For each unique *attB* site, flanking sequences of equal length were extracted to enable its status to be determined in other genomes. For most sites, a ~400 bp sequence from *S. pneumoniae* R6 was used to represent the intact *attB* site. For the $attB_{\mathrm{OXC}}$ and $attB_{ccnB}$ sites, the duplication of csRNA sequences upon prophage integration necessitated an ~800 bp sequence be used to represent the intact *attB* site. For the *attB*$_{cbpG}$ and *attB*$_{relA}$ sites, variation within the pneumococcal species required that alternative *attB* alleles were identified from GPS isolates, in addition to the *S. pneumoniae* R6 locus, to enable the identification of the intact, but divergent, loci. For the *attB*$_{ccnB}$ site, the extent of species-wide variation required that two alternative alleles were included from the GPS population.

This set of *attB* sequences was used to query the database of GPS isolate draft assemblies using BLASTN with default settings. An *attB* site was inferred to be intact in an isolate if the corresponding draft assembly contained a single continuous alignment to at least two-thirds of the *attB* sequence. This represented the situation in which the genome contained no prophage at this site, resulting in an uninterrupted alignment. The *attB* site was inferred to be absent if the draft assembly did not generate a single continuous alignment to at least one-third of the *attB* sequence. This represented the situation in which the locus was either absent from the genome, or not fully assembled. The *attB* site was inferred to be infected with a prophage in an isolate if the longest single continuous alignment to the corresponding draft assembly was between one-third and two-thirds of the *attB* sequence. This represented the situation when the integration of a prophage generates *attL* and *attR* sites, each of which is expected to match to half of the sequence representing the *attB* site.

These thresholds had to be adjusted for two *attB* sites. At *attB*$_{relA}$, the variable length of a BOX interspersed repeat sequence (Croucher et al. 2011) resulted in false positive inferences of prophage insertions, unless the threshold upper alignment length for inferring infections at this site was decreased to 230 bp (of a 401 bp query). At $attB_{hlpA}$, the duplication of sequence on prophage insertion resulted in insertions at the site being missed unless the threshold upper alignment length for inferring an uninfected *attB* site was increased to 280 bp (of a 400 bp query).

This characterisation of *attB* sites in individual genomes (Table S2), followed by a search of the entire collection for evidence of insertions at these locations, was undertaken iteratively until no more *attB* sites could be found. This resulted in the identification of 17 *attB* sites (Fig. S4; Table S2). At least one site was inferred to be disrupted by an insertion in 14,917 pneumococci (74.4% of the collection). These included 12,073 isolates identified as lysogenic through comparisons with known prophage sequences (80.9%). The *attB* sites that were most commonly disrupted in isolates in which no prophage could be detected were $attB_{scr}$ and $attB_{eno}$ (Fig. S1). The $attB_{scr}$ site is associated with the CIPhR element (Fig. S7). A BLASTN search was used to identify matches >3.75 kb in length to the *S. pneumoniae* ATCC 700669 CIPhR sequence at identities above 85% across the GPS isolates. The distribution of this element was able to explain the insertions at $attB_{scr}$ that were not caused by prophage (Fig. S1). For $attB_{eno}$, the discrepancy instead reflected a high frequency of PRCI insertions at this site. Therefore, while the prophage insertion at $attB_{eno}$ was analysed (Fig. S5), the site was explained from broader analyses of the distribution of prophage (Fig. 1B).

**Phylogenetics of integrase proteins**

For each *attB* site, a representative integrase protein sequence was extracted. These proteins were aligned with Muscle (Edgar 2004) v5.1, using the default settings. The phylogeny was generated using FastTree (Price et al. 2010) v2.1.11, using the default settings. The clades were manually identified and annotated with ggtree (Yu et al. 2017).

**Identification of regulatory systems**

Of the 672 prophage-like sequences (Croucher et al. 2014) identified by an analysis of 616 genomes from Massachusetts (Croucher et al. 2013), 134 corresponded to the CIPhR element (Croucher et al. 2009; Croucher et al. 2014) that inserts into *attB*$_{scr}$. The 538 sequences corresponding to active prophage were observed to contain proteins associated with the domains Peptidase_M78 (Pfam domain PF06114; now renamed IrrE N-terminal-like domain, Interpro domain IPR010359), corresponding to ImmA-type regulators; or Peptidase_S24 (Pfam domain PF00717; now renamed Peptidase S24/S26A/S26B/S26C, Interpro domain IPR015927), corresponding to C1-type regulators (Punta et al. 2012; Blum et al. 2021). ImmA-type proteins were encoded by 185 prophage, and C1-type regulators were encoded by 116 prophage. A systematic analysis of their distribution found all but three prophage longer than 30 genes were associated with one of these regulatory proteins, but none contained both systems (Fig. S10). The 237 prophage sequences not associated with either regulatory system were generally short, incomplete fragments of larger elements. Therefore, pneumococcal prophage appear to be predominantly regulated by one of two mutually-exclusive regulatory mechanisms: ImmAR-type or C1-type regulators.

These sequences were used to identify prophage regulatory proteins in the 20,047 GPS genomes (Gladstone et al. 2019). First, separate multiple sequence alignments of the 185 ImmA proteins, and the 116 CI proteins, were generated with MAFFT v7.305b (Katoh and Standley 2013). Hidden Markov models (HMMs) were generated from these alignments with HMMer v3.2.1 (Eddy 2011). These were then used to search the proteome of each isolate for proteins similar to the Massachusetts ImmA-type and C1-type representatives. The distributions of E values for alignments of both HMMs to proteins across all isolates were used to define an empirically-determined threshold of $10^{-30}$, for both protein types, to identify true positive phage regulators (Fig. S10).

Applying this threshold across isolates found 9,841 pneumococci (49.1%) contained an ImmA-type phage regulator, and 8,612 pneumococci (43.0%) contained a C1-type phage regulator. To test whether these systems were mutually exclusive, their distribution was analysed in the subset of 8,076 monolysogenic isolates, defined as those in which only a single *attB* site was disrupted. This found 577 instances of isolates encoding both systems (7.14% of monolysogenic isolates). The annotations of representative pneumococci suggested C1-type regulators were also found in some PRCIs. Therefore an HMM specific to PRCI C1-type regulator proteins was constructed using sequences extracted from genomes belonging to GPSC1, GPSC7, GPSC21, GPSC41 and GPSC64 (Fig. S10). A strict E value threshold of $10^{-125}$ was used to distinguish these from prophage C1-type regulators. Excluding the 2,913 proteins identified by this HMM from the prophage analysis reduced the number of monolysogenic isolates encoding both regulators to 235 (2.91% of monolyosgenic isolates). Given the diversity of PRCI and prophage sequences, and the imperfect nature of the assemblies, this seems consistent with the two systems being mutually exclusive of one another.

Using this refined approach, the number isolates encoding C1-type prophage regulators was reduced to 7,402 (36.9%). Overall, 13,394 pneumococci (66.8%) contained at least one representative of either regulator. Hence of the 12,194 isolates inferred to contain a prophage, based on sequence similarity to known representatives, only 596 (4.89%) did not contain either a C1-type or ImmAR-type prophage regulator.

**Synthesis of prophage analyses**

Combining the comparison with known prophage sequences, disruption of *attB* sites, and search for phage regulators (Fig. S2) found the three approaches reached a consensus on 11,463 pneumococci being lysogenic (57.2%) and 5,498 being non-lysogenic (27.4%). Among the 3,086 isolates on which there was no consensus across the three methods, the modal category (1,388 pneumococci) corresponded to isolates in which an *attB* site was disrupted, and a phage regulator was identified, but the draft assembly contained no detectable similarity with the collection of 63 prophage sequences. This is likely a consequence of the high levels of diversity observed across prophage. Consistent with this inference, the next most common categories of discordant results across the three analyses comprised instances where either no regulator could be identified with the HMMs trained on a limited dataset (589 isolates), or cases where there was no substantial match with the 63 prophage: pneumococci in which the only evidence of prophage was the detection of a regulator protein (393 isolates) or a disrupted *attB* site (574 isolates). These discrepancies may represent prophage with rare sequences, fragmented assemblies, or the acquisition of prophage proteins by other mobile genetic elements. Additionally, there were 135 isolates in which a prophage was detected, but no *attB* site appeared to be disrupted. Finally, there were seven isolates in which a prophage was detected, but no regulator or disrupted *attB* site could be identified.

Therefore these three analyses were consistent with one another in determining whether a pneumococcus was infected with a prophage. Although the search for similarity with known prophage sequences only provided a binary inference of whether an isolate was lysogenic or not, both the identification of disrupted *attB* sites, and phage regulators, quantified the number of distinct prophage per isolate. The estimates across all isolates were strongly correlated (Pearson's $R$ = 0.87; Fig. S2). Therefore the results of analyses should be consistent across either method. As the categorisation by regulatory protein type only divided prophage into two groups, the *attB* site analysis results were used by default, as they enabled a higher-resolution classification of the prophage.

**Combined analysis of integrases and *attB* sites**

The representative integrase sequences included near-identical proteins modifying *ccnC* at the most frequently infected *attB* site, $att_{\mathrm{OXC}}$, and a similar site within the paralogous csRNA gene, *ccnB* at $attB_{ccnB}$ (Furi et al. 2019; Garriss and Henriques-Normark 2020). Very similar integrases also recognised sites near *relA* and *ply*, neither of which shared sequence similarity with the csRNA genes (Fig. S7). Yet both were very close to BOX repeat elements, which can be transcribed and fold into stem-loop structures, similar to the csRNAs. Hence the recognition of the *attB* sites may depend on attributes other than just sequence. To test whether proximity to BOX elements was a factor, the distance of each *attB* site to the nearest BOX element was plotted (Fig. S7). This did not suggest a strong association between

BOX elements and *attB* sites across the genome. Instead, the low similarity between the *attB* sequences recognised by near-identical integrases appeared indicative of pneumococcal prophage being under diversifying selection to target *attB* sites not frequently occupied by other elements.

### Text S2: Effect of phase-variable restriction-modification systems on prophage deletion

*S. pneumoniae* isolates typically harbour two phase-variable restriction-modification systems (RMS): the inverting variable restriction (*ivr*) locus encodes the *Spn*III RMS (Croucher et al. 2014; Manso et al. 2014), and the translocating variable restriction (*tvr*) locus encodes the *Spn*IV RMS (Croucher et al. 2014; Kwun et al. 2018). Rearrangements at these loci are sufficiently rapid to occur within overnight culturing of pneumococci. This changes the sequence motifs targeted by the systems, consequently affecting genome-wide patterns of chromosomal methylation.

#### The effect of the *Spn*III and *Spn*IV systems on deletion of ϕ08B02743

Initial tests of the efficiency of prophage deletion by transformation used donor DNA extracted from derivatives of the *S. pneumoniae* 08B02743 isolate in which the prophage had been replaced with a Janus cassette (ϕ08B02743::Janus), and a base substitution conferring rifampicin resistance had been added. The recipient genotypes differed in the mutations introduced within ϕ08B02743. The ratios of prophage deletion to base substitution acquisition were generally below one, and varied considerably between recipient genotypes (Fig. S11). However, this variation in transformation efficiency was also associated with differences in growth, as previously observed for epigenetic variants of clinical isolates in which such phenotypic divergence was associated with alterations in the phase-variable *ivr* and *tvr* loci (Kwun et al. 2022; Kwun et al. 2023). These loci encode RMSs that can change the expression of the competence system (Kwun et al. 2023), inhibit the uptake of resistance genes (Kwun et al. 2018), and also function as an abortive infection system upon recognising prophage sequences as non-self (Furi et al. 2019).

Consistent with phase variation causing the observed variation in transformation efficiency, PCR amplification assays (Kwun et al. 2018) demonstrated that the recipient genotypes encoded different arrangements at the *tvr* locus. Therefore both the *ivr* and *tvr* loci were deleted from 08B02743 using the Janus cassette, and the prophage mutations reintroduced into this background.

The prophage deletion experiments were then repeated in the Δ*ivr* Δ*tvr* mutants (Table S3). The ratios measured in these experiments were more consistent across genotypes, and substantially higher. This is consistent with variation in the *Spn*III and *Spn*IV systems either inhibiting the acquisition of the Janus cassette, or targeting activated prophage through an abortive infection mechanism.

**Effects in other prophage systems**

The ratio of prophage deletion to rifampicin resistance acquisition through transformation was also measured for mutants of the ϕRMV8 and ϕRMV4 phage hosted by cells with intact *ivr* loci, and *tvr* loci that were fixed in a specific orientation through deletion of the *tvrR* recombinase (Kwun et al. 2018) (Fig. S11). These ratios were consistently lower than one across multiple prophage mutants. Therefore the *ivr* locus was deleted from each genotype. However, deletion of the entire *tvr* locus proved challenging in each of these backgrounds. Therefore experiments were undertaken in Δ*ivr* Δ*tvrR* cells (Table S3). Each time these genotypes were modified through the introduction of additional mutations, the arrangement of the *tvr* locus was established through PCR amplification assays (similar to those exemplified by Fig. S11) to check there was no variation between genotypes being assayed.

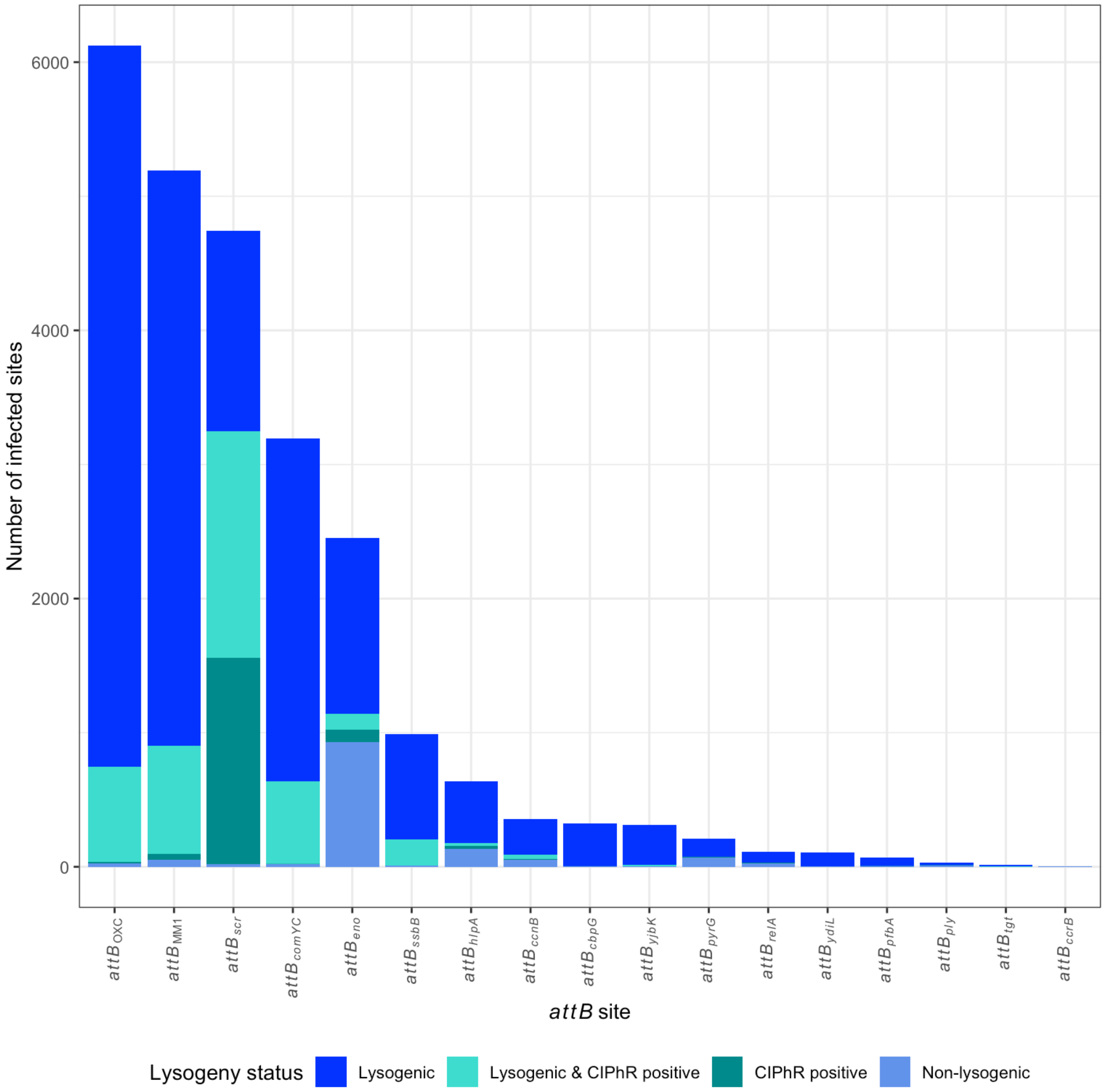


**Figure S1** Congruence between the analysis of disrupted *attB* sites and the identification of prophage and CIPhR sequences. BLASTN searches were used to identify pneumococci carrying prophage and CIPhR sequences (Text S1). Independent analyses of *attB* sites were used to identify those which appeared to contain an insertion (Text S1). Across all but two *attB* sites, isolates with evidence of insertions were predominantly associated with prophage sequences. For $att_{scr}$, the discrepancy was explained by the integration of CIPhR elements at this site. For $attB_{eno}$, the discrepancy was caused by phage-related chromosomal islands (PRCIs) frequently inserting at the same site. Therefore $attB_{eno}$ was excluded from further analyses of prophage distributions.

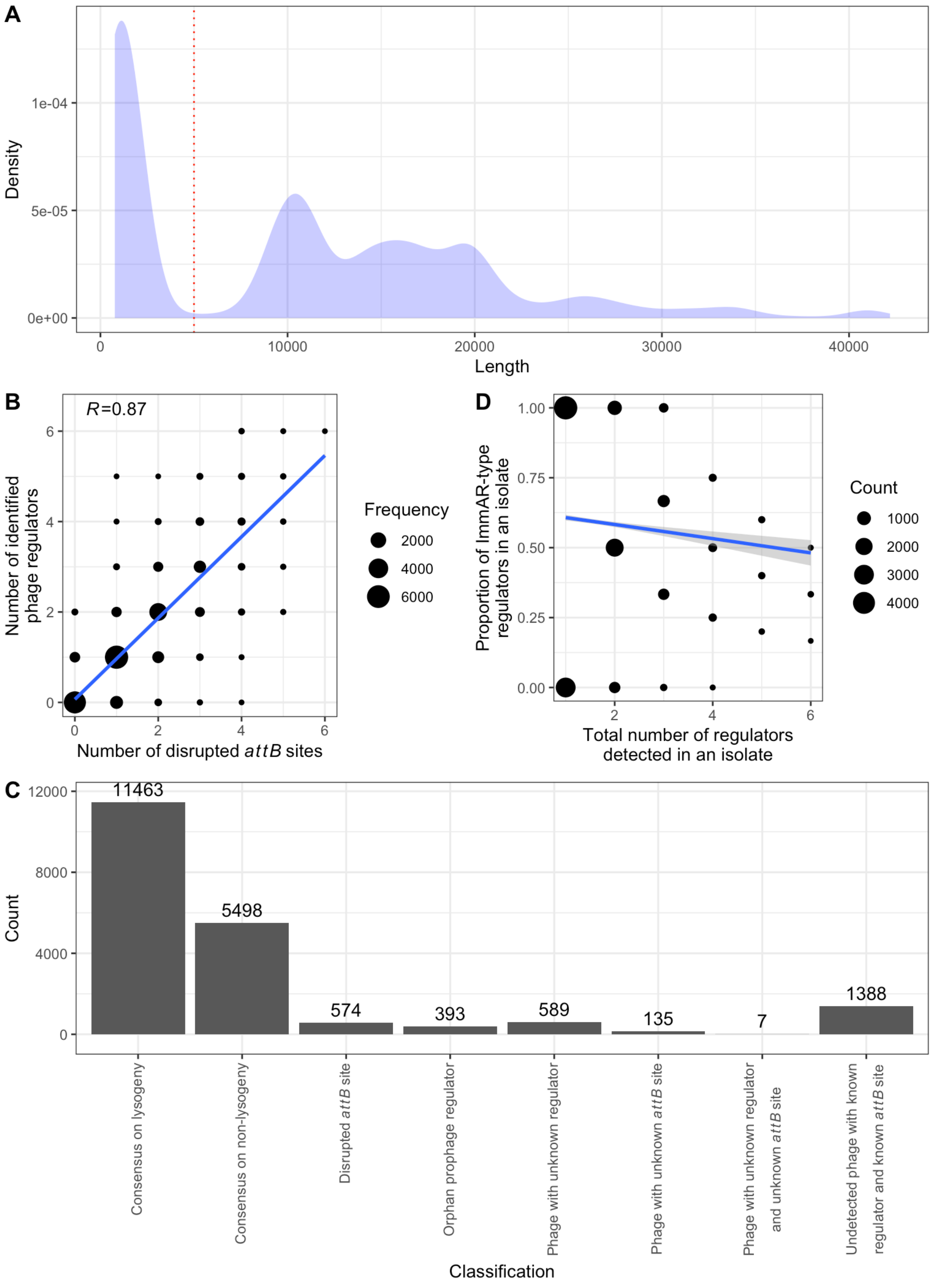
A
Density
1e-04
5e-05
0e+00
0
10000
20000
30000
40000
Length
B
R=0.87
Number of identified phage regulators
Number of disrupted attB sites
Frequency
2000
4000
6000
D
Proportion of ImmAR-type regulators in an isolate
1.00
0.75
0.50
0.25
0.00
Total number of regulators detected in an isolate
Count
1000
2000
3000
4000
C
12000
8000
4000
0
Count
11463
5498
574
393
589
135
7
1388
Consensus on lysogeny
Consensus on non-lysogeny
Disrupted attB site
Orphan prophage regulator
Phage with unknown regulator
Phage with unknown attB site
Phage with unknown regulator and unknown attB site
Undetected phage with known regulator and known attB site
Classification

**Figure S2** Estimating the number of pneumococci infected with prophage. (A) Density plot showing the distribution of the longest BLASTN alignment length between each of the 20,047 Global Pneumococcal Sequencing Project pneumococcal genomes and the 63 representative prophage. The vertical dashed line shows the threshold length of 5 kb used to distinguish false positive alignments from those indicating the presence of a prophage in the corresponding pneumococcal genome. (B) Scatterplot comparing the number of disrupted *attB* sites and the number of identified prophage regulatory proteins across each of the 20,047 Global Pneumococcal Sequencing Project pneumococcal genomes. The size of the point corresponds to the number of isolates found with the corresponding combination of values. The blue line represents the best-fitting linear model of the relationship. The Pearson's correlation coefficient, *R*, is annotated on the plot. (C) Barplot summarising the congruence of the three methods used to identify prophage in the 20,047 Global Pneumococcal Sequencing Project pneumococcal genomes. (D) Scatterplot showing the relationship between the number of prophage in an isolate from the Global Pneumococcal Sequencing Project, calculated as the sum of C1-type and ImmAR-type regulators, and the proportion of those regulators that were of the ImmAR-type. The size of the point represents the frequency of each specific combination of regulator counts in the collection. The blue line represents the maximum likelihood fit of a generalised linear model of the binomial family, using a logit link function, to these data. The grey shaded region shows the 95% confidence intervals of this model output.

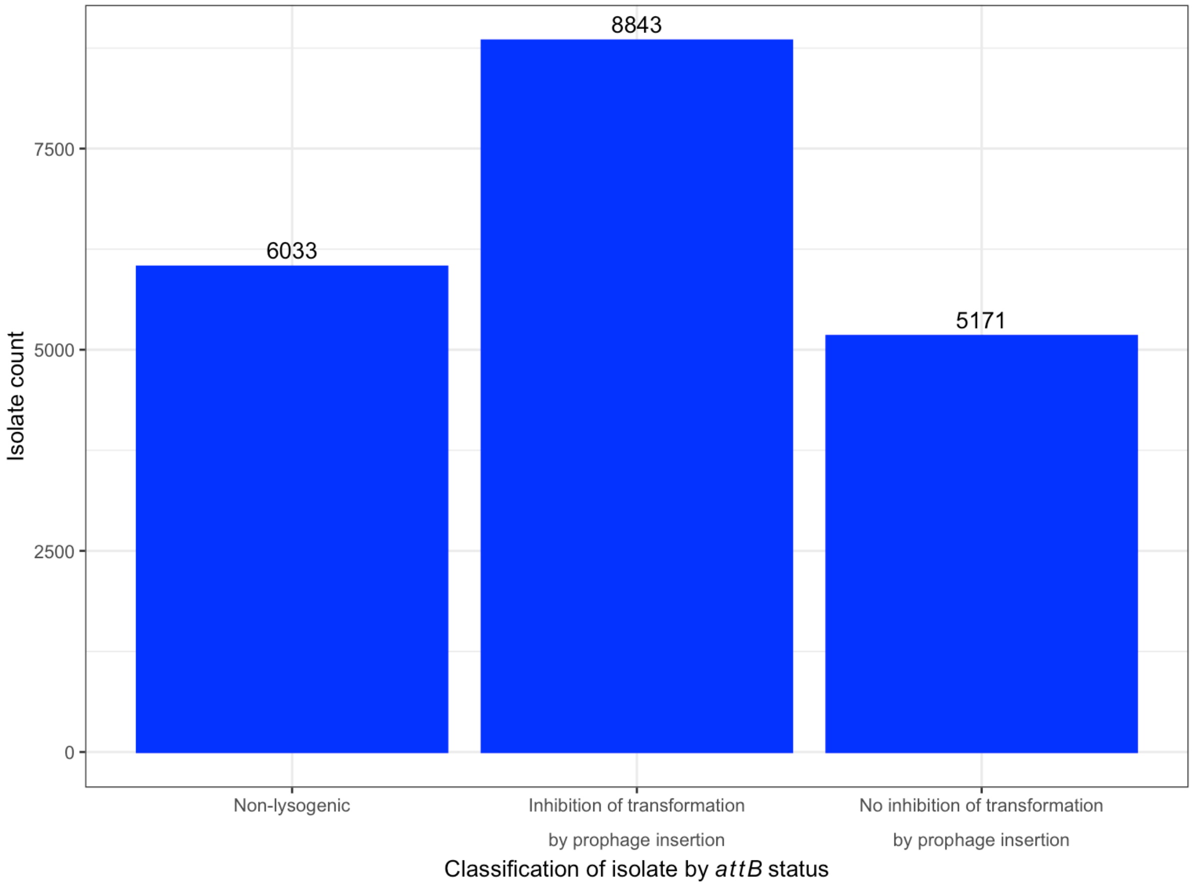


**Figure S3** Barplot showing the effect of prophage integration on transformation efficiency across the 20,047 Global Pneumococcal Sequencing Project pneumococcal genomes. Isolates were inferred to be lysogenic if at least one *attB* site was disrupted in the genome. Transformation was classified as being inhibited if a prophage was inserted in one of the *attB* sites associated with reducing the activity of the competence system: $attB_{comYC}$, $attB_{ssbB}$, $attB_{\mathrm{OXC}}$, or $attB_{yjbK}$.

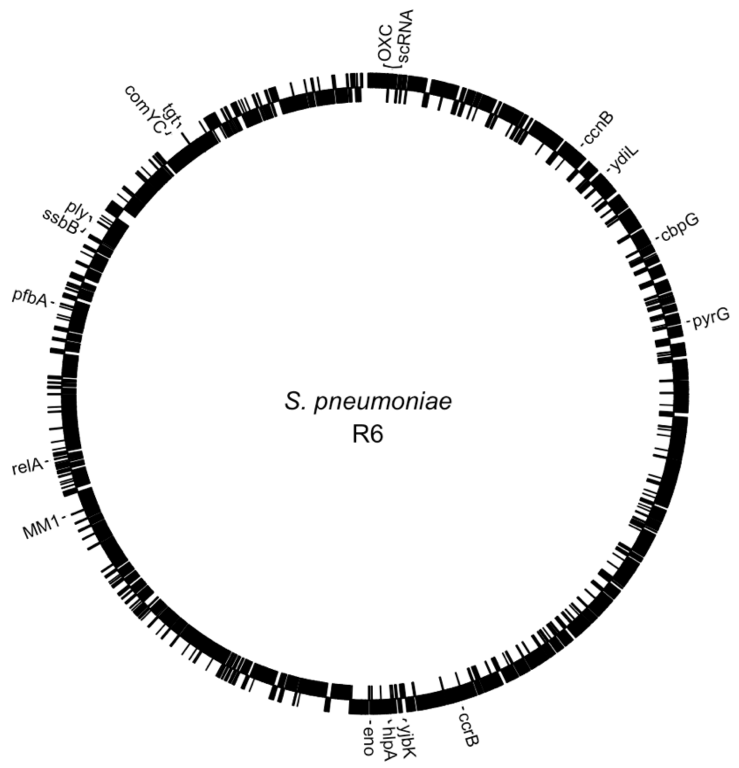


**Figure S4** Distribution of the identified *attB* sites around the *S. pneumoniae* R6 genome (Table S2). Black boxes in the outer ring show protein coding sequences encoded by the top strand of the genome, while those in the inner ring show protein coding sequences encoded by the bottom strand of the genome. The chromosome is divided into two replichores by the origin of replication at the top, and terminus of replication near the bottom. These correspond with the strong coding bias of the genome, with the protein coding sequences aligned with replication of the leading strand.

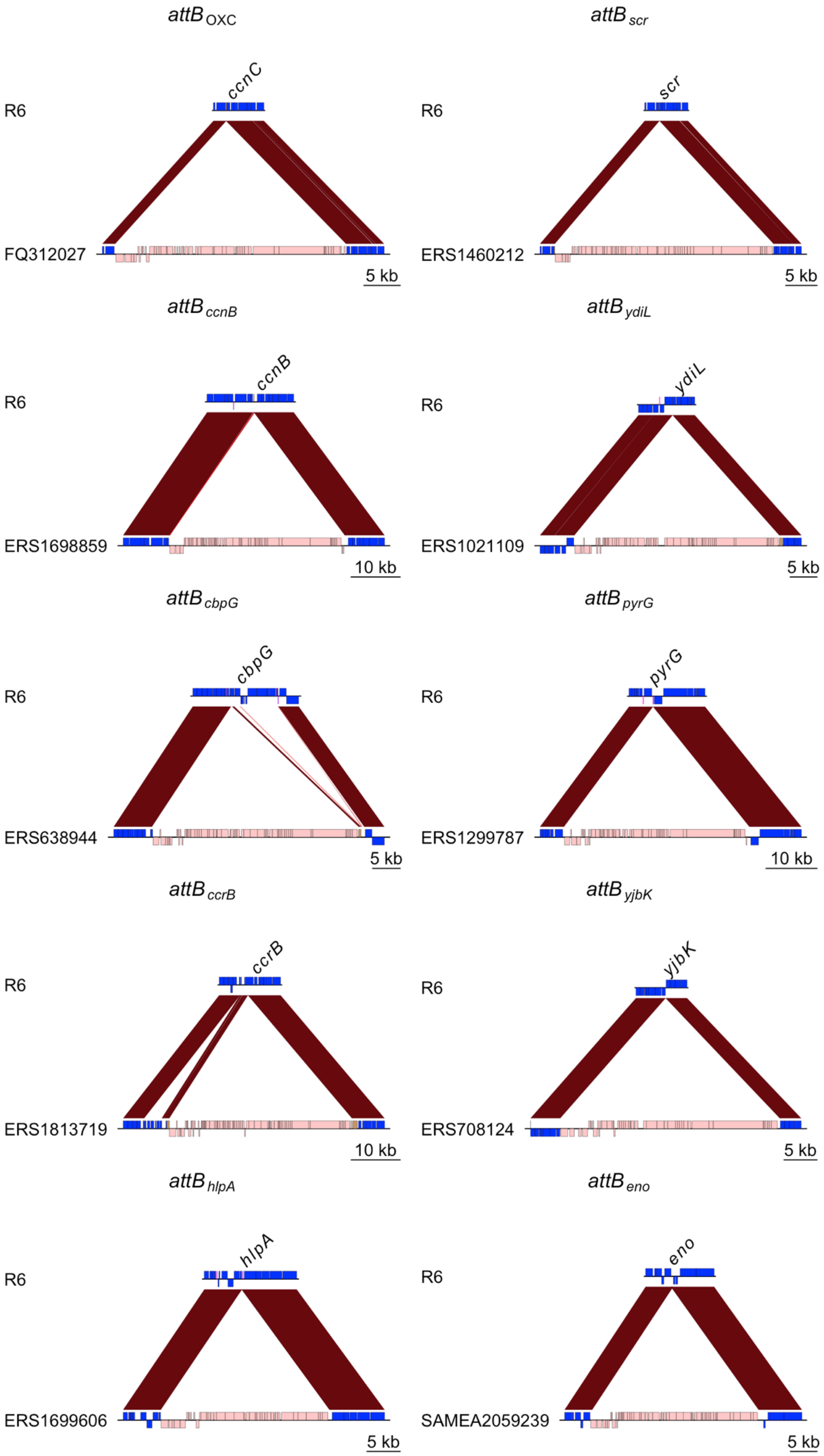
attB OXC
attB scr
R6
ccnC
FQ312027
5 kb
R6
scr
ERS1460212
5 kb
attB ccnB
attB ydiL
R6
ccnB
ERS1698859
10 kb
R6
ydiL
ERS1021109
5 kb
attB cbpG
attB pyrG
R6
cbpG
ERS638944
5 kb
R6
pyrG
ERS1299787
10 kb
attB ccrB
attB yjbK
R6
ccrB
ERS1813719
10 kb
R6
yjbK
ERS708124
5 kb
attB hlpA
attB eno
R6
hlpA
ERS1699606
5 kb
R6
eno
SAMEA2059239
5 kb

**Figure S5** Prophage insertions at the ten *attB* sites in the right-hand replichore in Fig. S4. Each panel shows the alignment of the uninfected *attB* sites in *S. pneumoniae* R6 with the infected site in another isolate, annotated with the accession code of the sample in the European Nucleotide Archive. The red bands link regions of similar sequences, as identified by BLASTN, coloured as described in Fig. 1D. Coding sequences are represented by boxes above the line, when encoded by the top strand, or below the line, when encoded on the bottom strand. Coding sequences within the prophage are coloured pink, while others are coloured blue. As the uninfected *attB* sites are shown in the same orientation in each panel, this demonstrates the lysogeny genes of the prophage are consistently arranged counter to the coding bias of the genome, whereas the replication, structure and lysis genes are arranged in alignment with the coding bias of the genome.

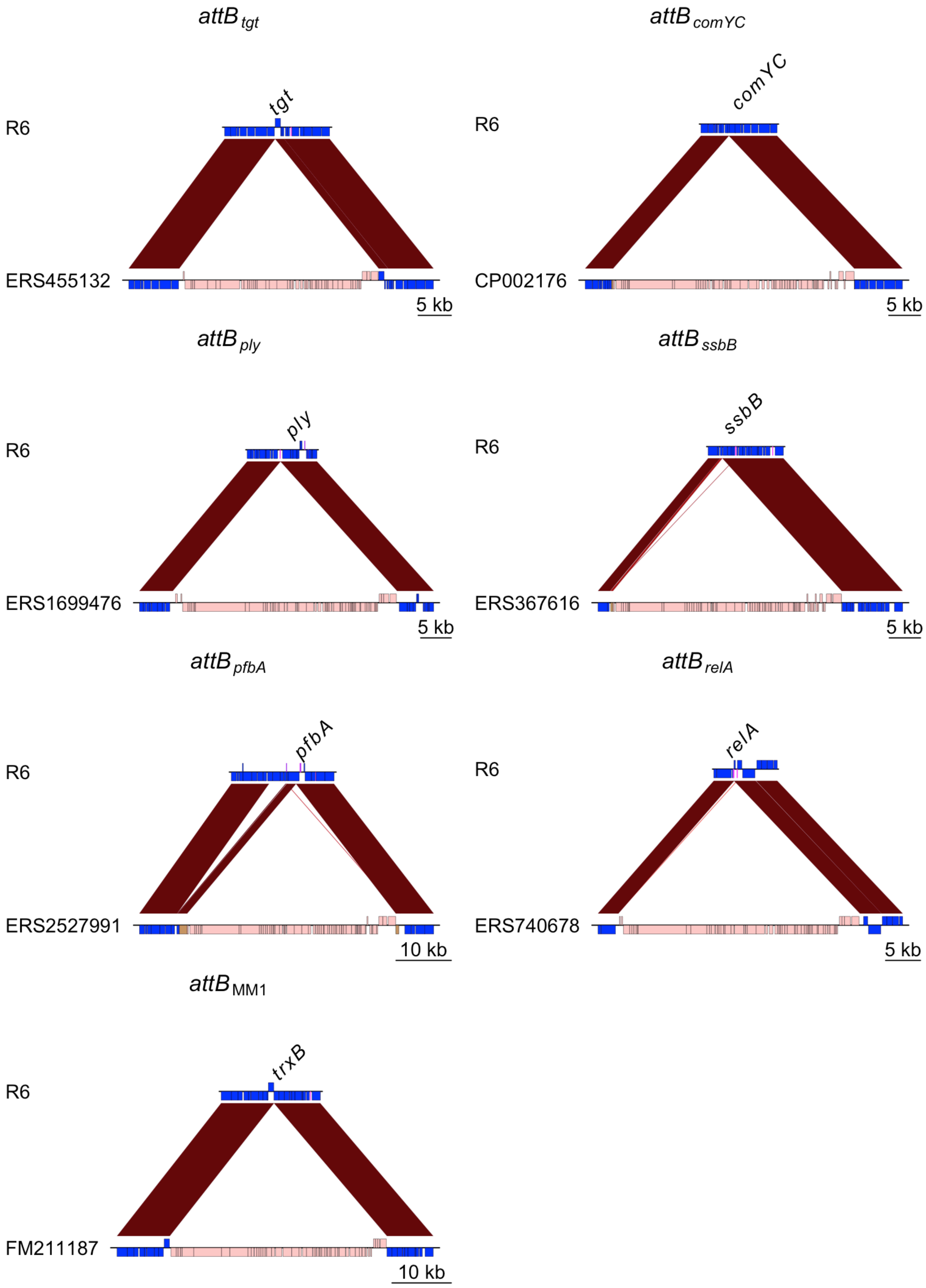


**Figure S6** Prophage insertions at the seven *attB* sites in the left-hand replichore in Fig. S4. Data are shown as in Fig. S5. As the uninfected *attB* sites are shown in the same orientation in each panel, this again demonstrates the replication, structure and lysis genes are consistently aligned with the coding bias of the genome.

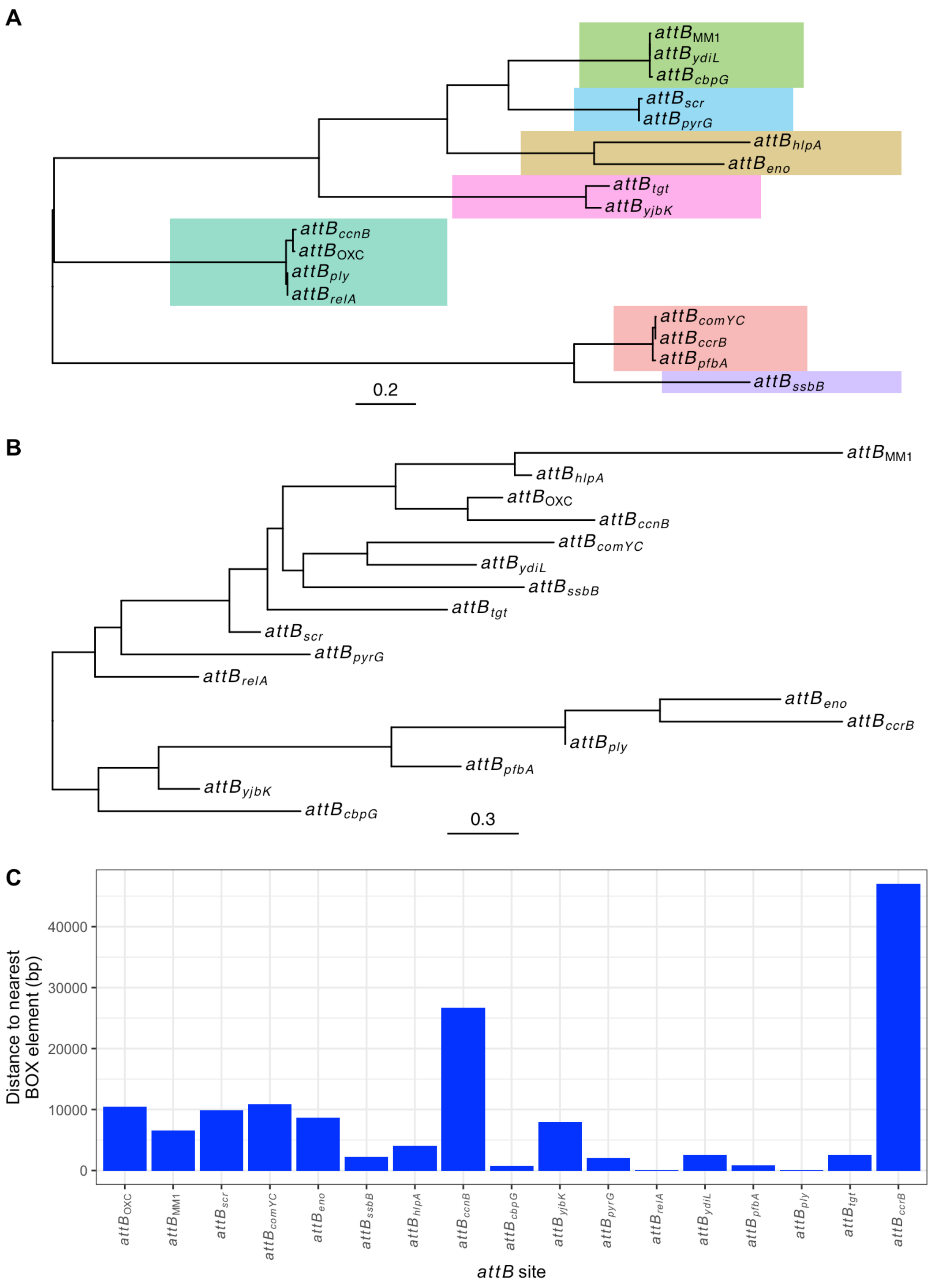


**Figure S7** Comparing the properties of *attB* sites. (A) Diversity of prophage integrase proteins. A single integrase sequence was selected to represent each *attB* site, and used to generate this maximum-likelihood phylogeny, shown rooted at its midpoint. Clades of closely-related integrases are annotated by coloured blocks. (B) Diversity of prophage *attB* sites. This midpoint-rooted maximum likelihood phylogeny was generated from 25 nt sequences representing each *attB* site in the *S. pneumoniae* R6 genome. (C) Barchart showing the distance between each *attB* site and the nearest BOX element annotated in the *S. pneumoniae* R6 genome.

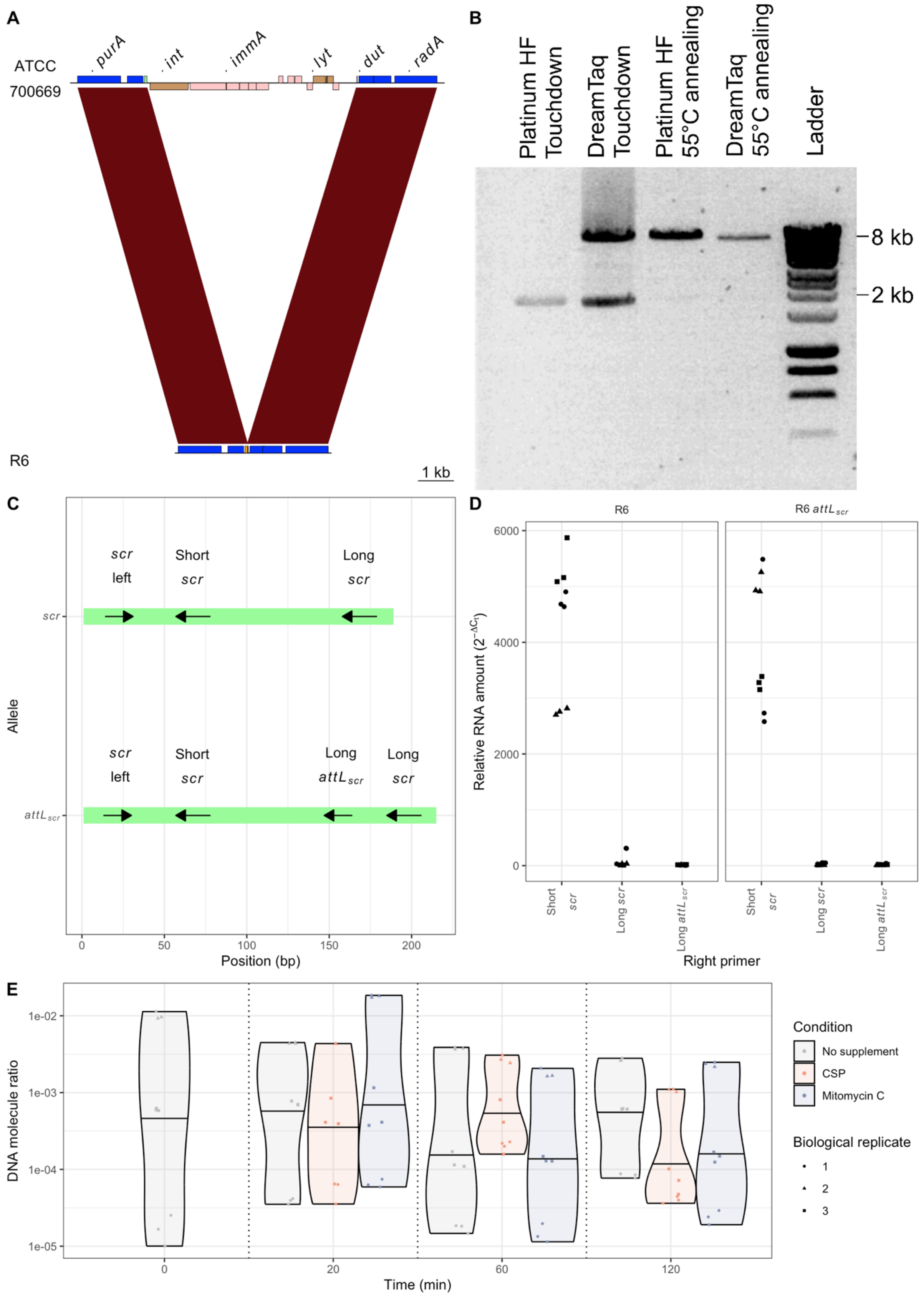


**Figure S8** Characterisation of the CIPhR element. (A) Comparison of *S. pneumoniae* ATCC 700669, which has a CIPhR insertion (Table S1), with the unmodified locus in *S. pneumoniae* R6. The red bands link regions of similar sequence identified by BLASTN. The boxes indicate genes, which are above the middle line if they are encoded on the top strand of the genome, and below the line if they are encoded on the bottom strand of the genome. Blue boxes represent cellular genes; pink boxes represent intact ClPhR genes; brown boxes represent ClPhR pseudogenes, and the green box represents the *scr* RNA gene, encoding the scRNA. (B) Assaying excision of the ClPhR element in *S. pneumoniae* 11930, a transformable representative of the PMEN1 strain. PCR amplification of the ClPhR locus was undertaken using two different polymerases under both standard (using a 55°C annealing temperature) and touchdown thermocycling programmes. Agarose gel electrophoresis revealed two bands, corresponding to amplicons with sizes of ~2 kb and ~8 kb. The longer amplicon was of the size expected when the ClPhR element was integrated into the chromosome. The shorter amplicon was of the size expected when the *attB* locus was restored when the element was excised. Dideoxy terminator sequencing of the shorter amplicon confirmed this corresponded to the empty *attB* locus. (C) Positions of primer binding sites used to amplify the *scr* gene. As only a subsection of the *scr* gene annotated in *S. pneumoniae* R6 matches the corresponding Rfam family (accession code RF00169), multiple combinations of primers were used to determine the length of the transcribed gene. These primers targeted the short version of the *scr* gene, which corresponded with the Rfam domain, and the long version of the *scr* gene, which matched the R6 annotation. The long version of the *scr* gene would be modified by the integration of the ClPhR element, and therefore an additional primer was used to detect whether such an altered sequence was correctly generated in the R6 mutant $attL_{scr}$ (Fig. 1C). (D) Results of quantitative PCR on cDNA generated from R6 and a derivative containing the $attL_{scr}$ sequence generated by the integration of the ClPhR element. Each point represents a technical replicate measurement of the amount of detected RNA, relative to the abundance of *rpoA* transcripts. The shapes of the points show the biological replicate from which the measurement was taken. These results demonstrated the short version of the *scr* gene was abundant in the tested cells, but the long forms were present at only negligible levels. Hence pneumococci produce the short form of the RNA. (E) Quantification of the excision of the ClPhR element. The ratio of the *attB* sites, generated by ClPhR excision, to *attL* sites, generated by ClPhR integration, was estimated using qPCR in early exponential phase and 20, 60 and 120 minutes after exposure to CSP or MMC. Control samples were taken concurrently from samples growing in unsupplemented media. The ratios were close to $10^{-3}$, consistent with very low levels of excision, and were unchanged in response to either CSP or MMC.

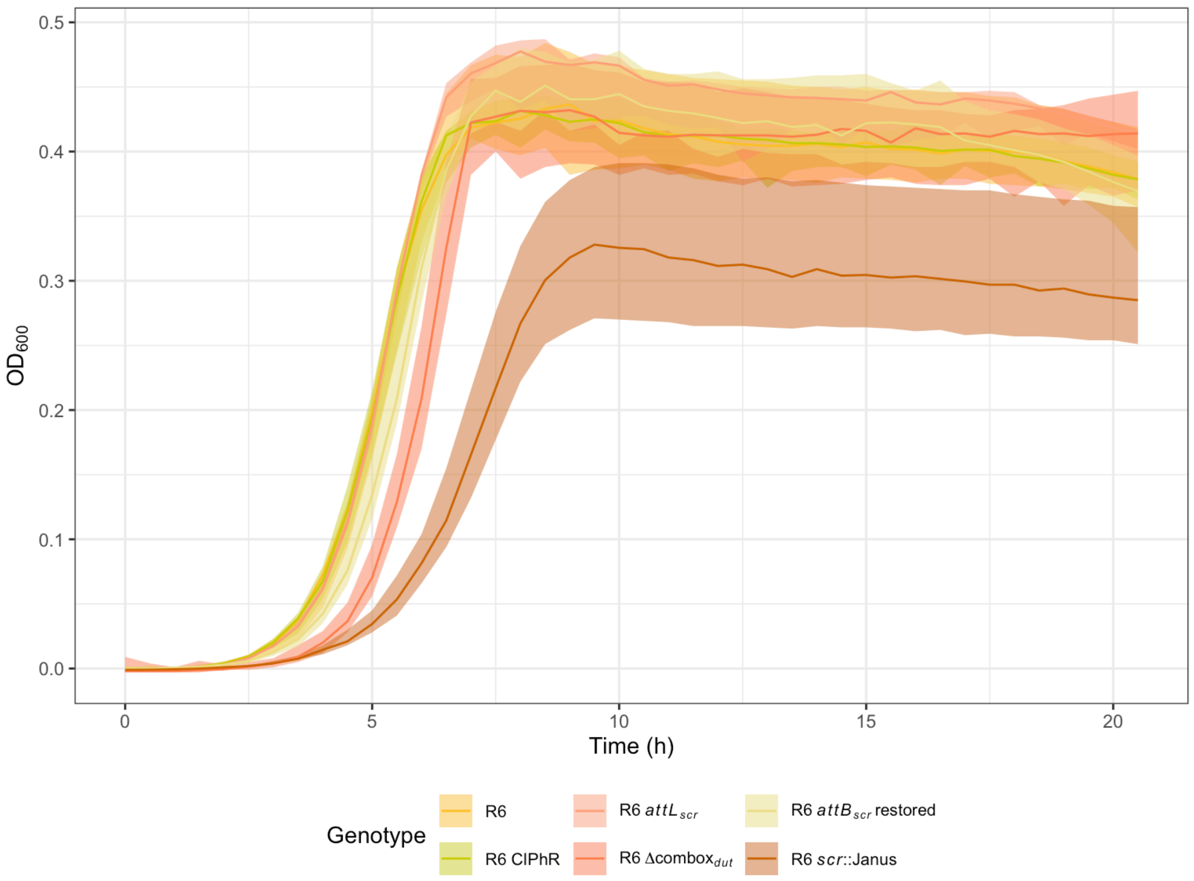


**Figure S9** Effect of mutations at the *attB*$_{scr}$ locus on the growth of *S. pneumoniae* R6 in unsupplemented media. Three biological replicates were measured for each genotype. The solid lines represent the median optical density at 600 nm ($OD_{600}$), and the shaded ribbon shows the range of $OD_{600}$ values.

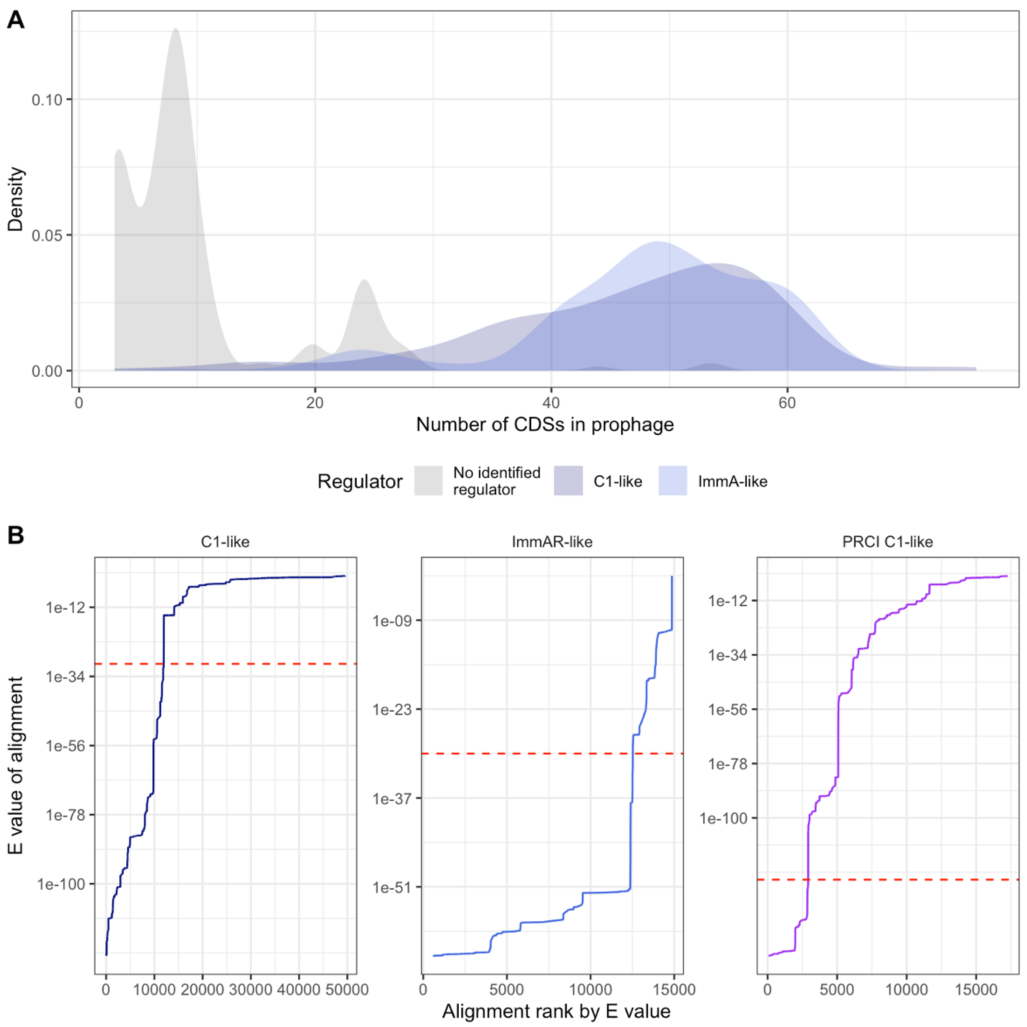


**Figure S10** Identification of prophage regulatory proteins. (A) Analysis of the regulatory proteins in the 538 prophage-like sequences identified in a set of 616 isolates from vaccine surveillance in Massachusetts. Elements were categorised by whether they contained the Peptidase S24/S26A/S26B/S26C domain (Interpro domain IPR015927), the IrrE N-terminal-like domain (Interpro domain IPR010359), or neither. The density plot shows the distribution of the number of genes in each prophage-like sequence, grouped by the regulatory protein types. This demonstrates that almost all full-length prophage (containing at least ~30 genes) contained either a Peptidase S24/S26A/S26B/S26C domain or a IrrE N-terminal-like domain. (B) Calibrating the hidden Markov models (HMMs) used to identify prophage regulatory proteins. HMMs were generated for C1-like proteins, corresponding to the Peptidase S24/S26A/S26B/S26C domain (Interpro domain IPR015927), and ImmA-like proteins, associated with the IrrE N-terminal-like domain (Interpro domain IPR010359), using proteins from the Massachusetts dataset. Scanning the proteome of the 20,047 Global Pneumococcal Sequencing Project pneumococcal genomes with these HMMs highlighted the need to exclude false positive matches to phage-related chromosomal islands (PRCIs), requiring a further HMM to be generated to PRCI C1-like regulator proteins (Text S1). The line graph shows the E values of the most significant alignments of the HMMs to the proteome, ordered by their rank. The horizontal dashed lines in each panel represent the threshold that separates the top-ranked, highly significant alignments from the majority of less significant, false positive alignments.

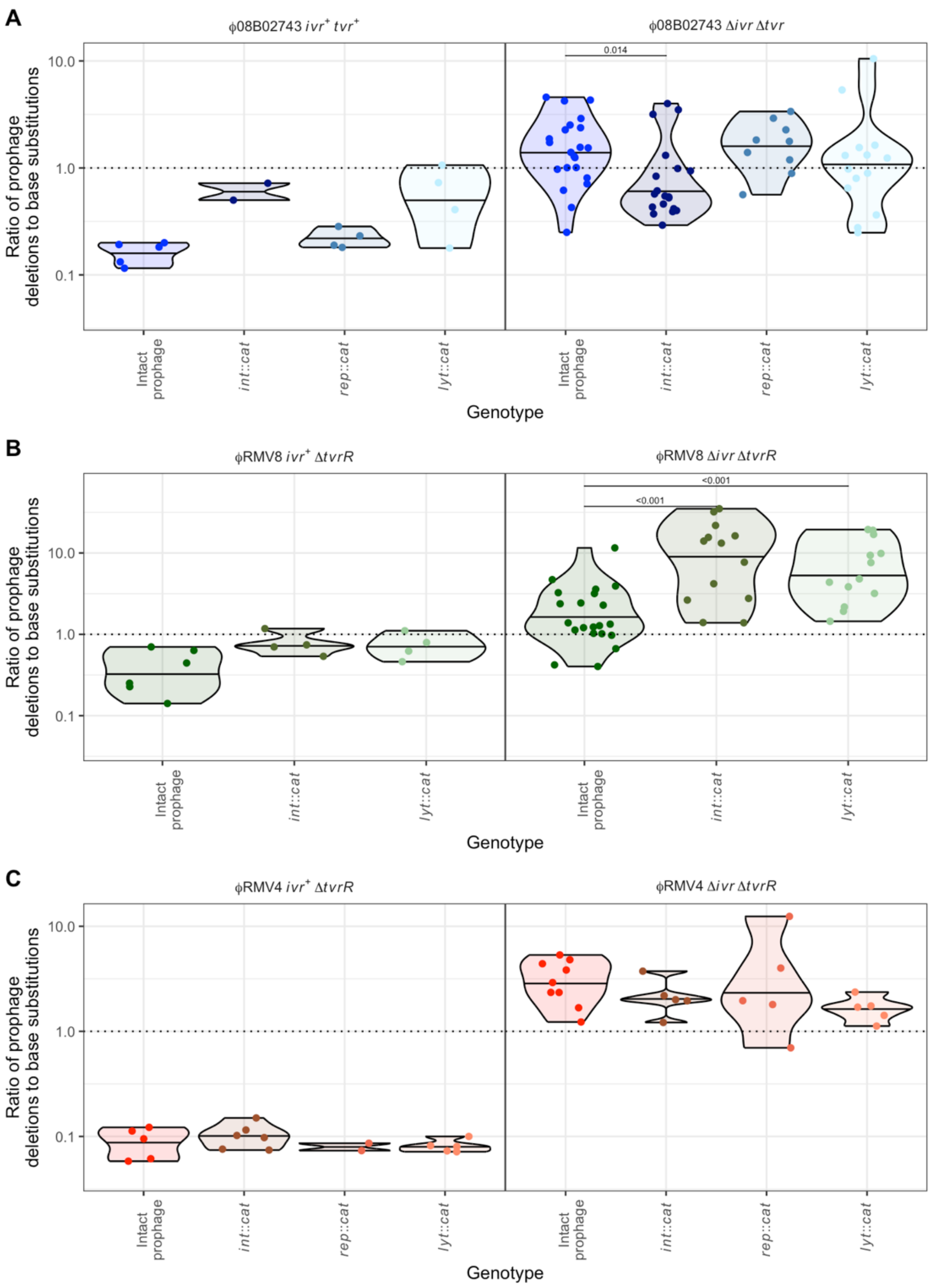


**Figure S11** Effect of phase-variable restriction-modification systems on the deletion of prophage by transformation. Each plot shows the ratio of prophage replacements to acquisitions of a base substitution, as in Fig. 2E. Each point represents an independent biological replicate, separated along the horizontal axis by the genotype of the recipient in the transformation experiment. (A) Prophage deletion ratios in the ϕ08B02743 system. The left panel shows the ratios measured when the *ivr* and *tvr* loci were intact. The right panel shows the equivalent measurements when both loci were deleted. (B) Prophage deletion ratios in the ϕRMV8 system. The left panel shows the ratios measured when the *ivr* was intact, and the arrangement of the *tvr* locus was fixed by deletion of the *tvrR* recombinase gene. The right panel shows the equivalent measurements when the *ivr* locus was removed, while the *tvr* locus remained locked. (C) Prophage deletion ratios in the ϕRMV4 system, shown as described for panel (B).

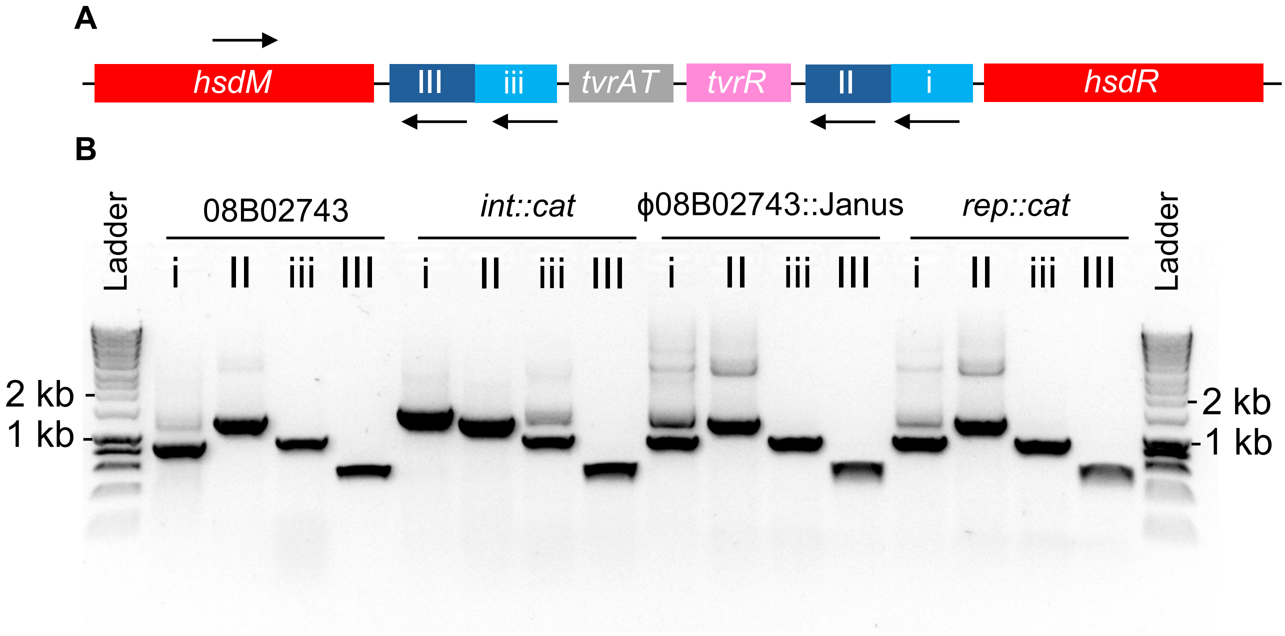


**Figure S12** Variation in the *tvr* locus associated with phenotypic differences. (A) Diagram showing an example of a typical *tvr* locus structure. The boxes represent protein coding genes. The arrow above the genes represents the binding site of the left primer used for PCR amplification, at a fixed position within the *hsdM* methylase gene. The arrows below the genes represent right primers, each of which is specific to an *hsdS* gene fragment, represented in blue, and annotated with Roman numerals. These fragments are rearranged through excision and reintegration by the *tvrR* recombinase gene, which changes the specificity of the Type I restriction-modification system encoded by the locus. These alterations are detectable through changes in the sizes of amplicons produced with different primer combinations. (B) Agarose gel electrophoresis result showing variation of the *tvr* locus in the 08B02743 *ivr*$^+$ *tvr*$^+$ genotype. The bands represent amplicons generated by PCRs primed by oligonucleotides binding to the locus, as represented in panel (A). The patterns demonstrated the wild type and *rep*::*cat* mutant share a common *tvr* arrangement that is distinct from that of the *int*::*cat* mutant. These differences were consistent with the less efficient deletion of the prophage in the wild type and *rep*::*cat* cells, relative to the *int*::*cat* genotype (Fig. S11), being associated with this change in the arrangement of the *tvr* locus.

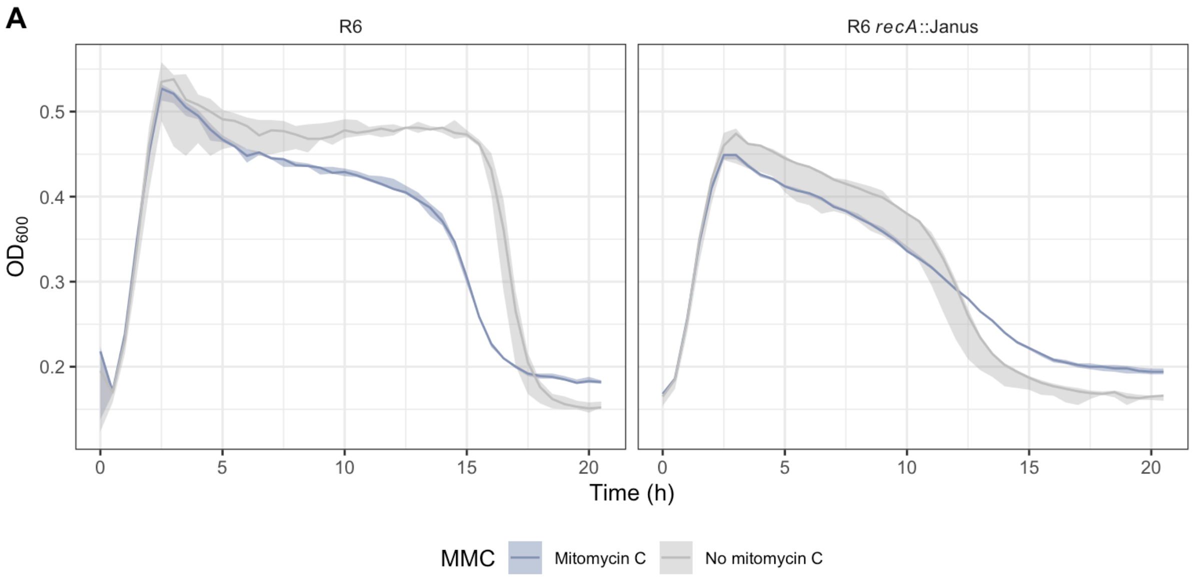


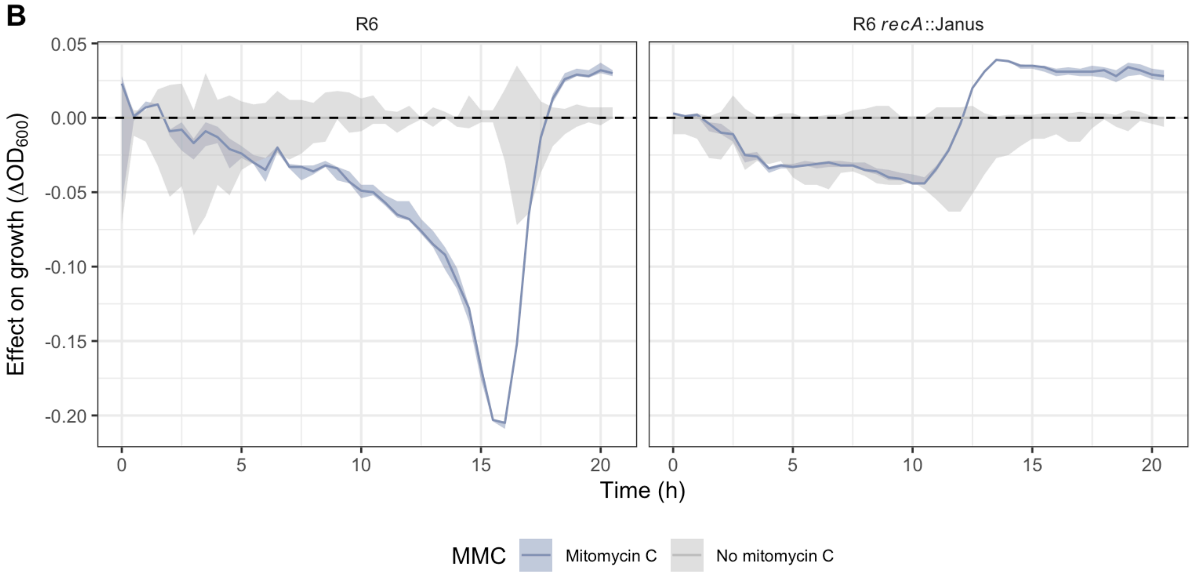


**Figure S13** Line plots showing the effect of the concentration of mitomycin C used to induce the prophage systems on the growth of the non-lysogenic isolate *S. pneumoniae* R6, and its *recA*::Janus mutant derivative. (A) Growth curves inferred using the optical density of cultures measured at 600 nm ($OD_{600}$). The solid line shows the median $OD_{600}$ observed in each condition, and the shaded ribbon shows the range of values observed across three biological replicates. The drop in $OD_{600}$ at the end of the experiment was driven by the lysis of these unencapsulated cells, demonstrating that neither cell replication, nor death, are strongly affected by this MMC concentration. (B) The effect of mitomycin C on growth, quantified through subtracting the median $OD_{600}$ measurements in the untreated control cultures from the equivalent measurements from the corresponding treated culture, using the data shown in panel (A).

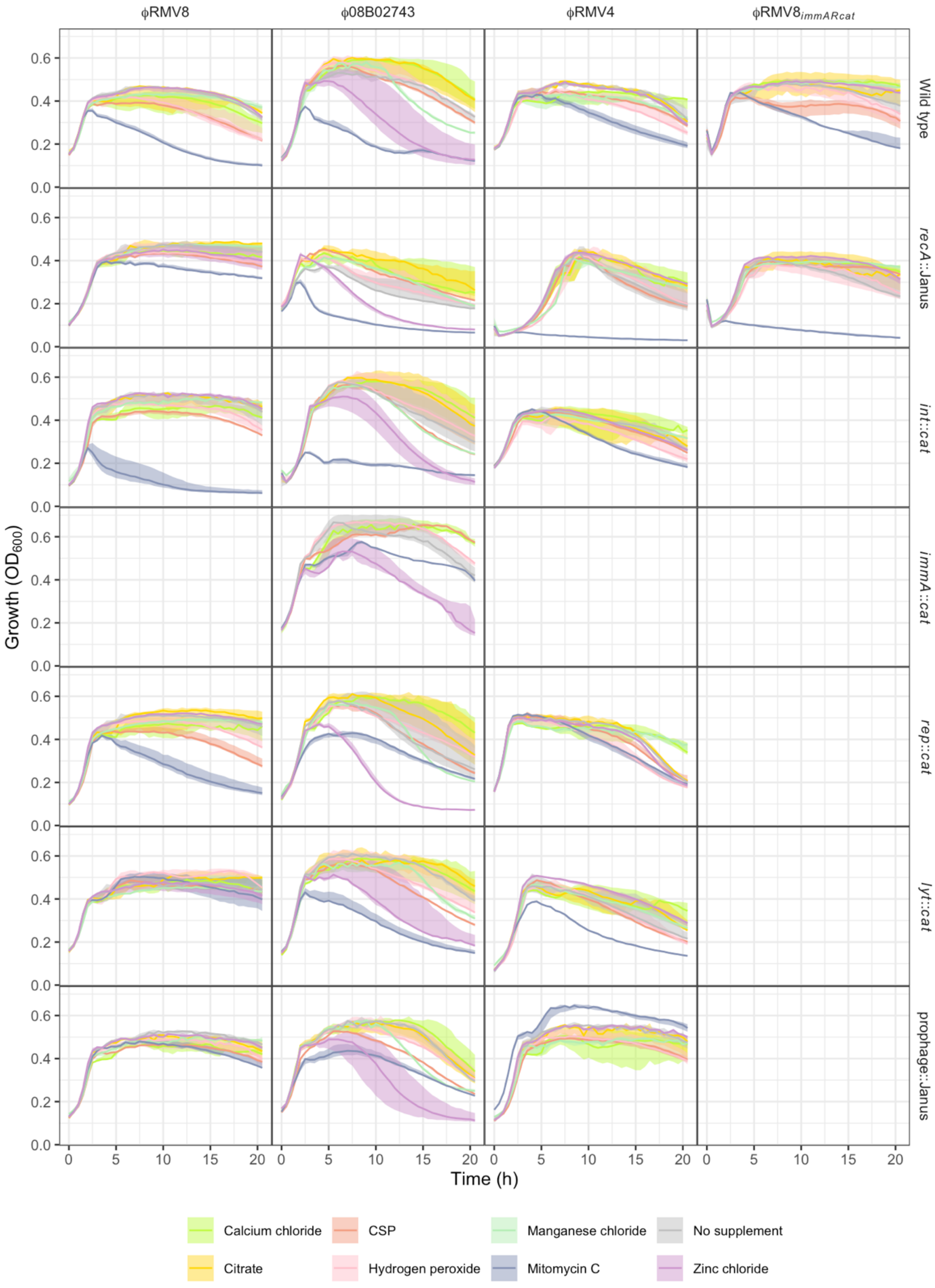


**Figure S14** Effect of exogenous chemical stimuli on the growth of prophage systems. Each panel shows the growth of a different genotype in unsupplemented media, and in the presence of different chemicals that may trigger prophage activation. At least three biological replicates were measured for all combinations of genotype and chemical stimulus. The solid lines represent the median optical density at 600 nm ($OD_{600}$), and the shaded ribbon shows the range of $OD_{600}$ values. Each column corresponds to a different prophage system, and each row corresponds to an equivalent set of mutants.

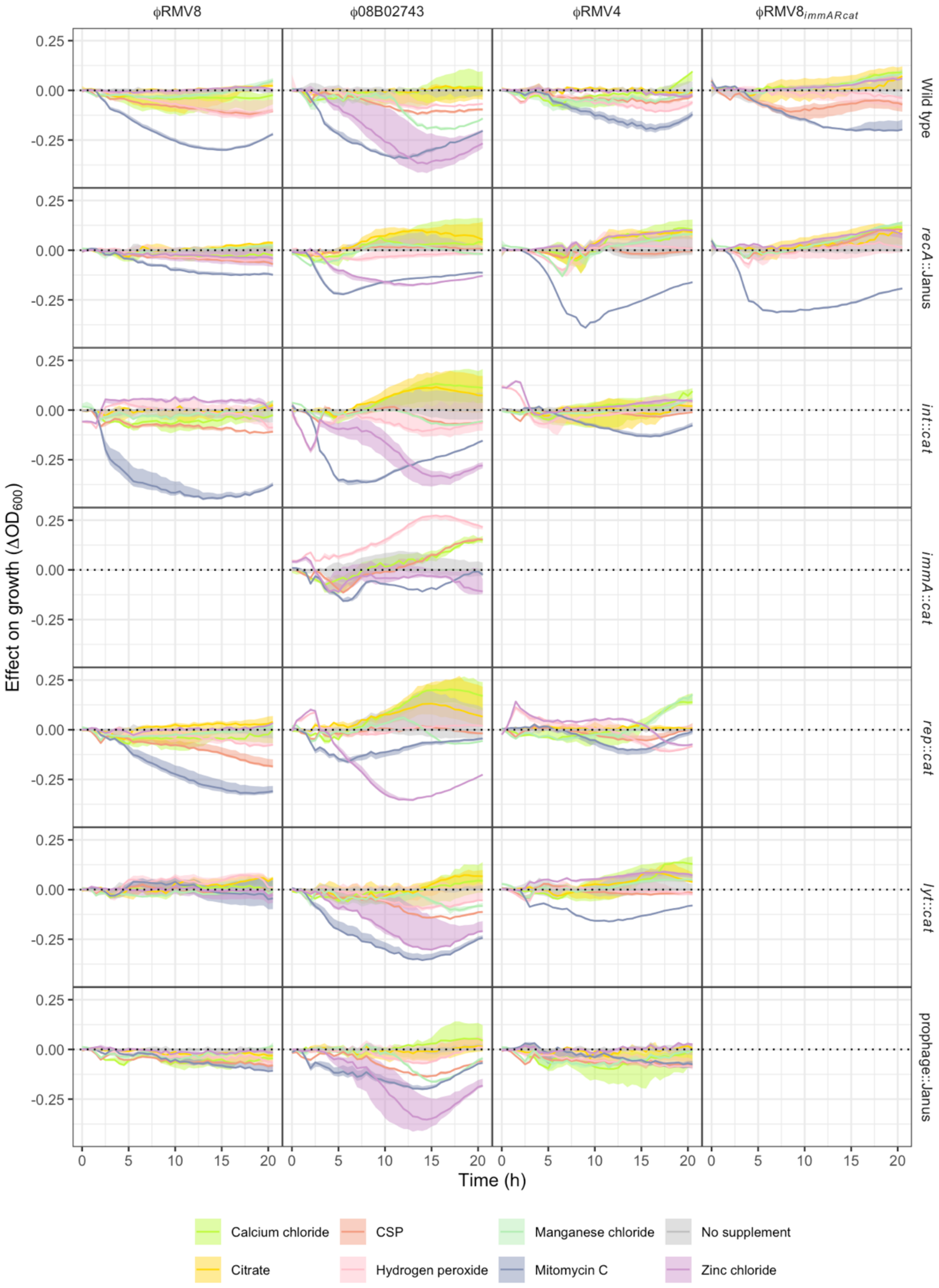


**Figure S15** Effect of exogenous chemical stimuli on the growth of prophage systems, relative to their growth in unsupplemented media. Each panel shows the difference between the median $OD_{600}$ of the culture in supplemented media, and that in unsupplemented media, as a solid line. The shaded area shows the range of $OD_{600}$ values observed in supplemented media, similarly adjusted by subtracting the median $OD_{600}$ observed in unsupplemented media. The grey ribbon shows the variation in the growth observed in unsupplemented media. At least three biological replicates were measured for all combinations of genotype and chemical stimulus. Each column corresponds to a different prophage system, and each row corresponds to an equivalent set of mutants.

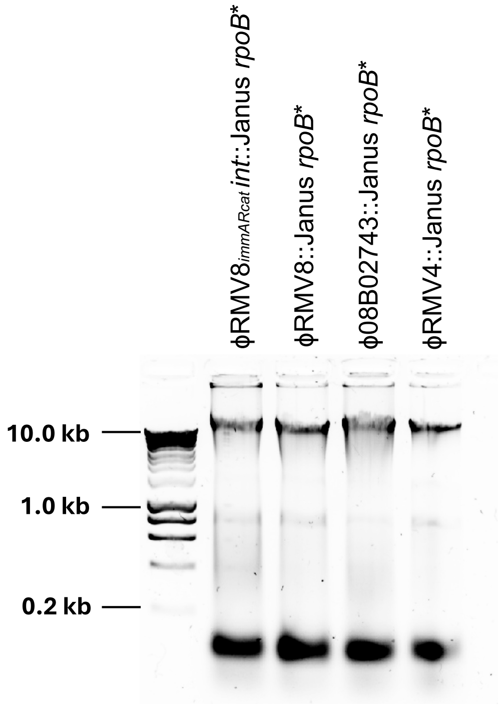


**Figure S16** Agarose gel electrophoretic separation of molecules by size in a 2.5 µg sample of four donor DNA samples, relative to the Bioline HyperLadder 1 kb. Three of these samples were used to assay the rates of prophage deletion (ϕRMV8::Janus *rpoB**, ϕ08B02743::Janus *rpoB** and ϕRMV4::Janus *rpoB**), while the fourth (ϕRMV8$_{immARcat}$ *int*::Janus *rpoB**) was used to assay the rate of prophage insertion. The similarity of the DNA molecule size distributions demonstrates that the difference in the insertion and deletion rates is not an artefact of using highly fragmented donor DNA in the insertion experiments.

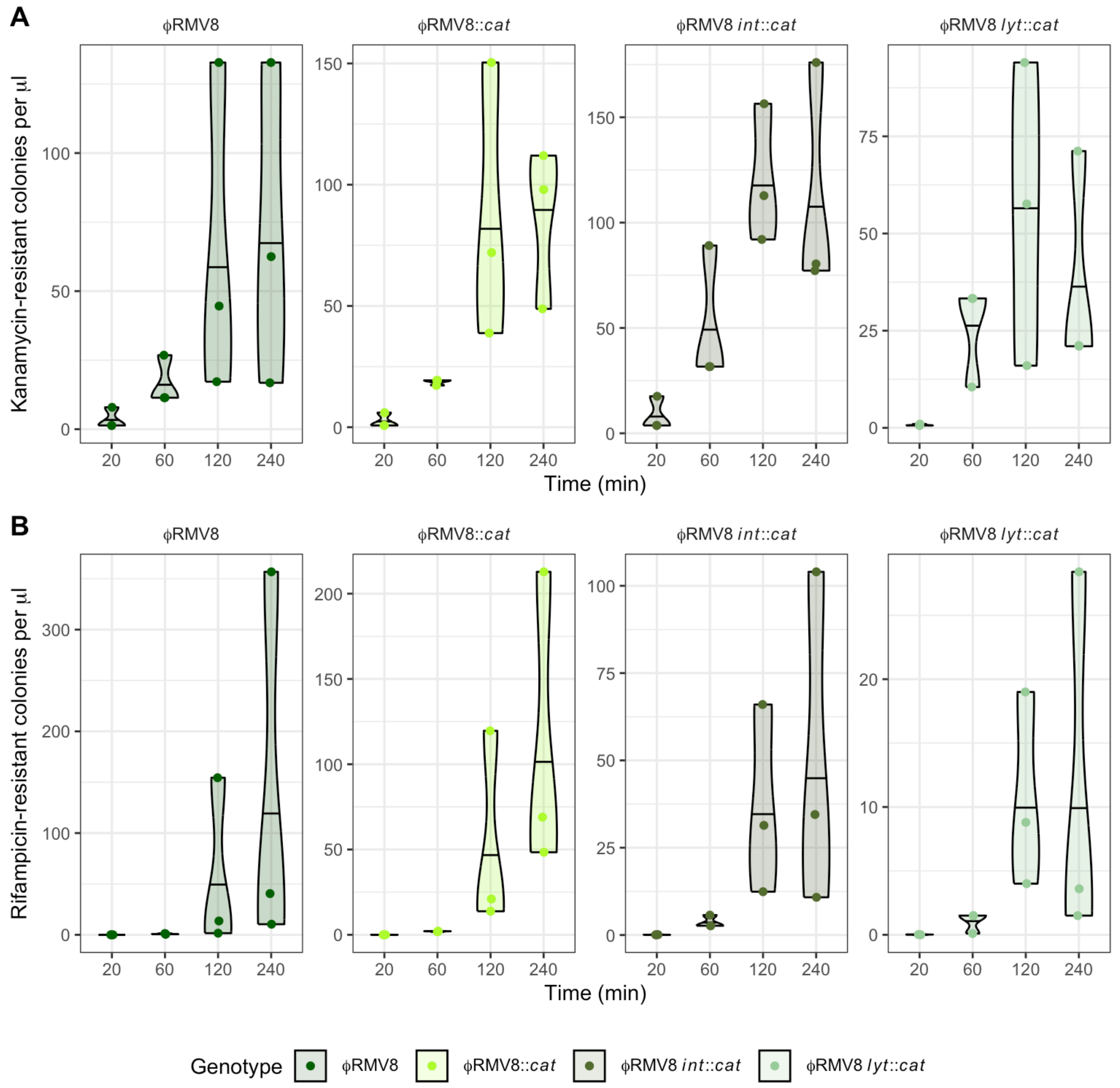


**Figure S17** Timing of antibiotic resistance phenotype expression following transformation. Four different recipient genotypes, shown in separate panels, were each transformed with DNA from an ϕRMV8 donor in which the prophage had been replaced with a Janus cassette, and the *rpoB* gene contained a base substitution conferring rifampicin resistance. The frequency of kanamycin- and rifampicin-resistant colonies, generated by expression of acquired Janus and *rpoB* sequences respectively, was measured at 20, 60, 120 and 240 min after transformation. Each point represents an independent biological replicate. Their distribution is summarised by violin plots, which include horizontal lines to indicate the median. The 120 min timepoint was sufficient to enable the detection of both kanamycin and rifampicin resistance, while minimising the time for selection to alter the ratio between these phenotypes.

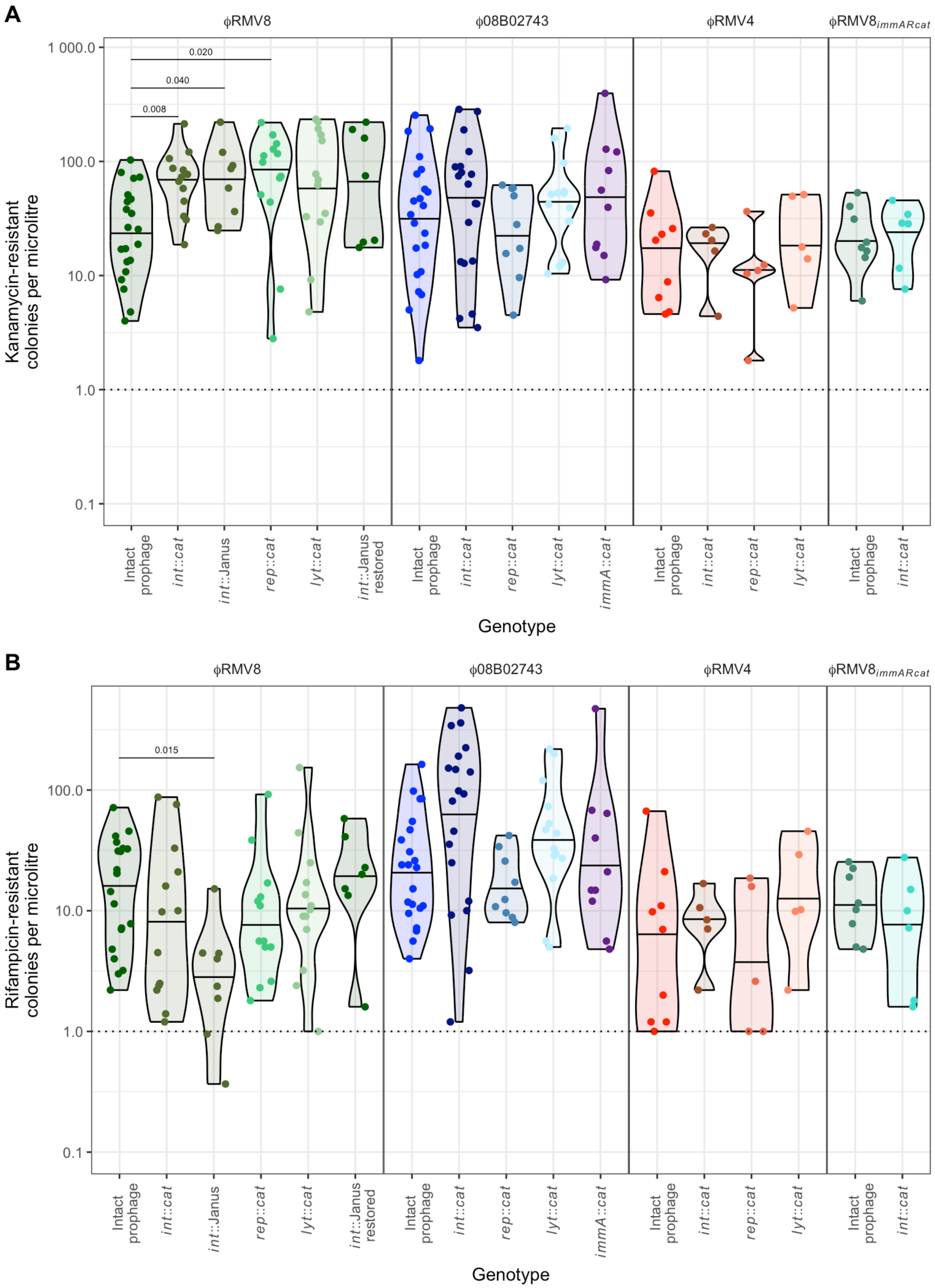


**Figure S18** Graphs showing the frequency of (A) kanamycin-resistant and (B) rifampicin-resistant colonies used to calculate the ratios shown in Fig. 2E.

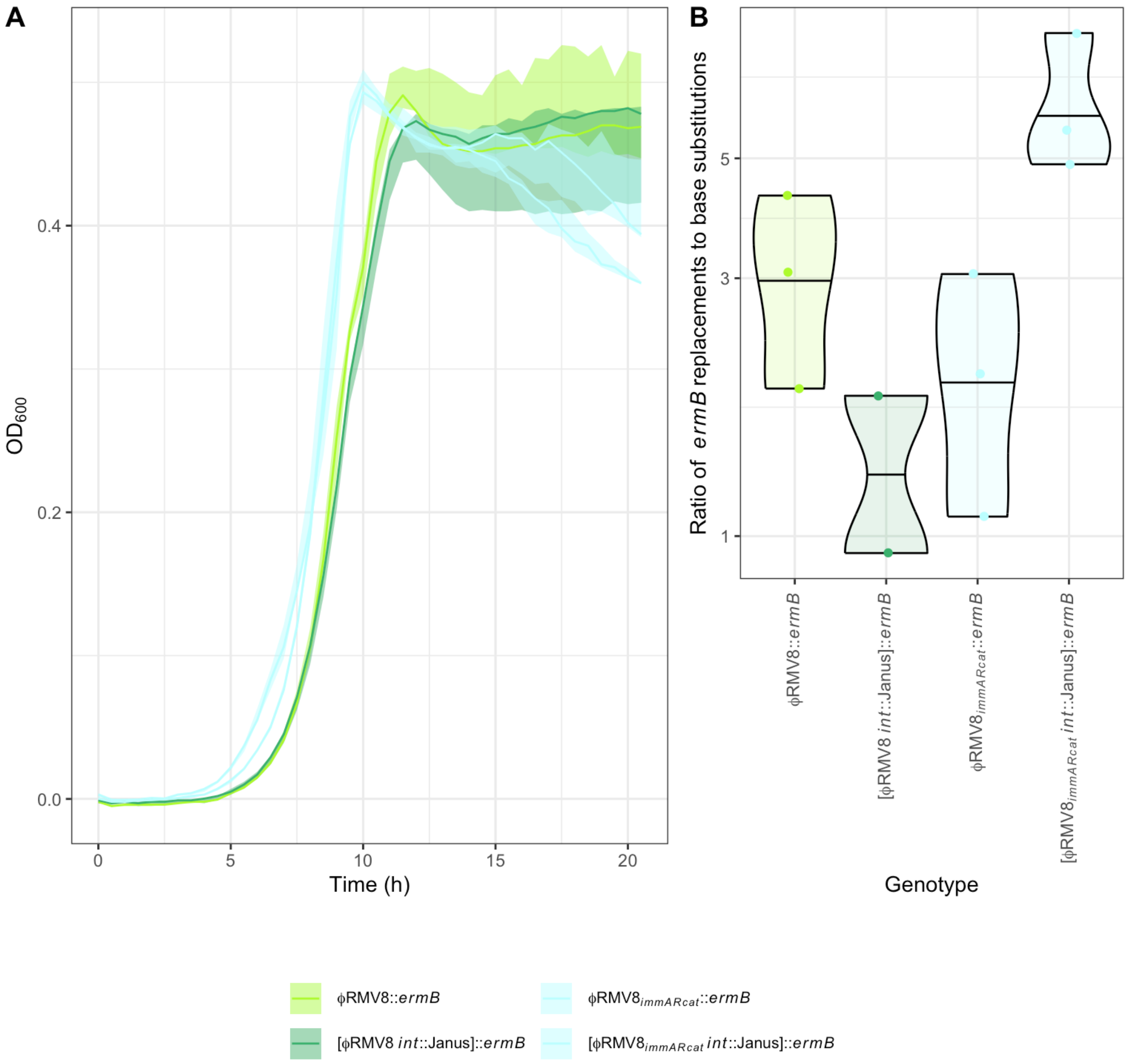


**Figure S19** Confirmation of the differential role of the integrase protein in evading deletion by transformation in ϕRMV8 and ϕRMV8$_{immARcat}$. While the ϕRMV8 *int*::Janus phage could be more efficiently removed by transformation than the intact prophage, no such difference was observed with the ϕRMV8$_{immARcat}$ *int*::Janus mutant (Fig. 2E). It is possible these could represent changes in the local homologous recombination frequency of the genome in the ϕRMV8 *int*::Janus mutant that were independent of the prophage. Therefore ϕRMV8 and ϕRMV8$_{immARcat}$, and the *int*::Janus derivatives of each, were replaced with an *ermB* erythromycin resistance marker. (A) These four genotypes exhibited similar growth profiles, suggesting there were no changes outside of the prophage locus that were affecting the growth rate. (B) Ratio of replacement of the *ermB* gene at the *attB* site by a Janus cassette, relative to the acquisition of a base substitution causing rifampicin resistance. Data are shown as in Fig. 2E. There was little difference between the genotypes, with no evidence of the ratio being highest in (ϕRMV8 *int*::Janus)::*ermB*. Therefore the higher deletion ratio of ϕRMV8 *int*::Janus in Fig. 2E can be attributed to the mutation in the prophage, not to changes in local recombination rates, or unexpected alterations occurring elsewhere in the genome.

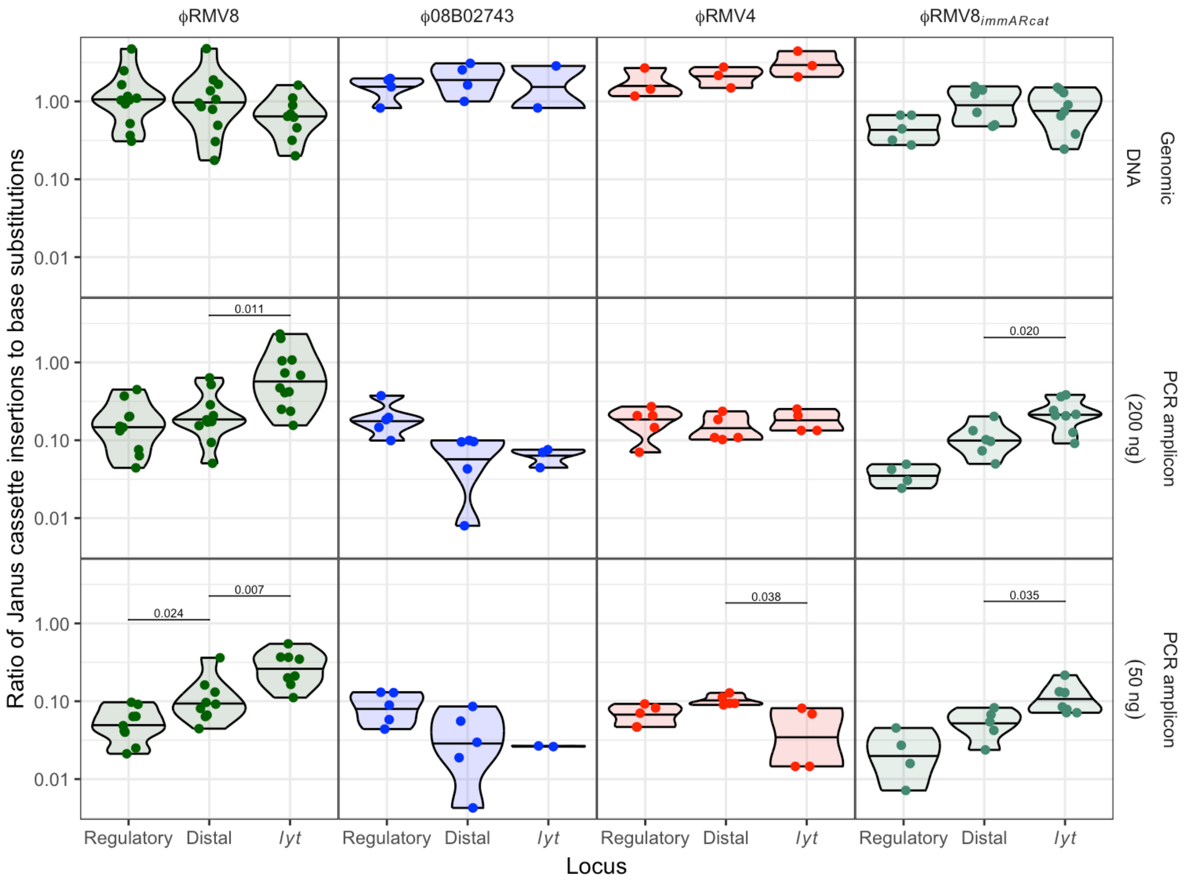


**Figure S20** Testing for the activation of prophage by transformation with targeted DNA fragments. Each point represents an independent biological replicate in which the ratio between the acquisition of a Janus cassette targeting part of a prophage, and a base substitution conferring resistance to rifampicin, was calculated. For each prophage system, a donor was constructed with a Janus cassette inserted within ~1 kb of the regulatory genes (the "regulatory" locus, encoding either a C1-type or ImmAR-type system); another had a Janus cassette inserted around 5 kb from the regulatory genes (the "distal" locus), and a final donor targeted the lytic amidase gene, the locus within the prophage furthest from the regulatory genes (Fig. 1D). Recipient cells of the same prophage system type were co-transformed with this DNA, and a consistent concentration of a PCR product generated from the *rpoB* gene that contained a base substitution conferring rifampicin resistance. The Janus cassette markers were added as genomic DNA (top row), a 3 kb PCR amplicon at high concentration (middle row), or the same PCR amplicon at low concentration (bottom row). The hypothesis was that ϕRMV8 would be more efficiently activated by the low concentration PCR amplicons targeting the regulatory locus, where they would interact with C1-type proteins associated with their binding sites within the prophage, than by those amplicons corresponding to sequences further away from the C1-type binding sites. The increasing activation of the prophage would be detectable as a decrease in the acquisition of the prophage-disrupting marker, relative to the acquisition of the rifampicin-resistance conferring base substitution. This effect was not expected in ImmAR-regulated prophage, as ImmA is not directly bound to prophage DNA, enabling it to interact with DNA in the cytosol, or annealed to other sites in the genome. Similarly, a lower, or no, difference was expected between Janus cassette insertions targeting different loci when transformation used genomic DNA, as long DNA fragments would be imported by the cell, allowing the acquisition of even distal markers to be associated with the import of DNA that would bind the regulatory locus. In agreement with the hypothesis, the relative efficiency with which the prophage-disrupting marker was acquired when PCR amplicons were used as donor DNA decreased as the marker targeted DNA closer to the regulatory locus. This was not observed when ϕRMV8 was transformed with the same markers carried on genomic DNA, or in experiments in which the ImmAR-regulated ϕ08B02743 or ϕRMV4 were transformed with equivalent markers on PCR amplicons. However, the ratio with which the resistance markers were acquired from PCR amplicons changed in the same way with ϕRMV8$_{immARcat}$ as with ϕRMV8. This suggests that the pattern was driven by properties of this genotype, rather than the prophage regulatory system. Therefore no evidence was found of transforming DNA targeting the prophage regulatory locus being more efficient at activating cell lysis as a consequence of increased induction of C1-type regulators.

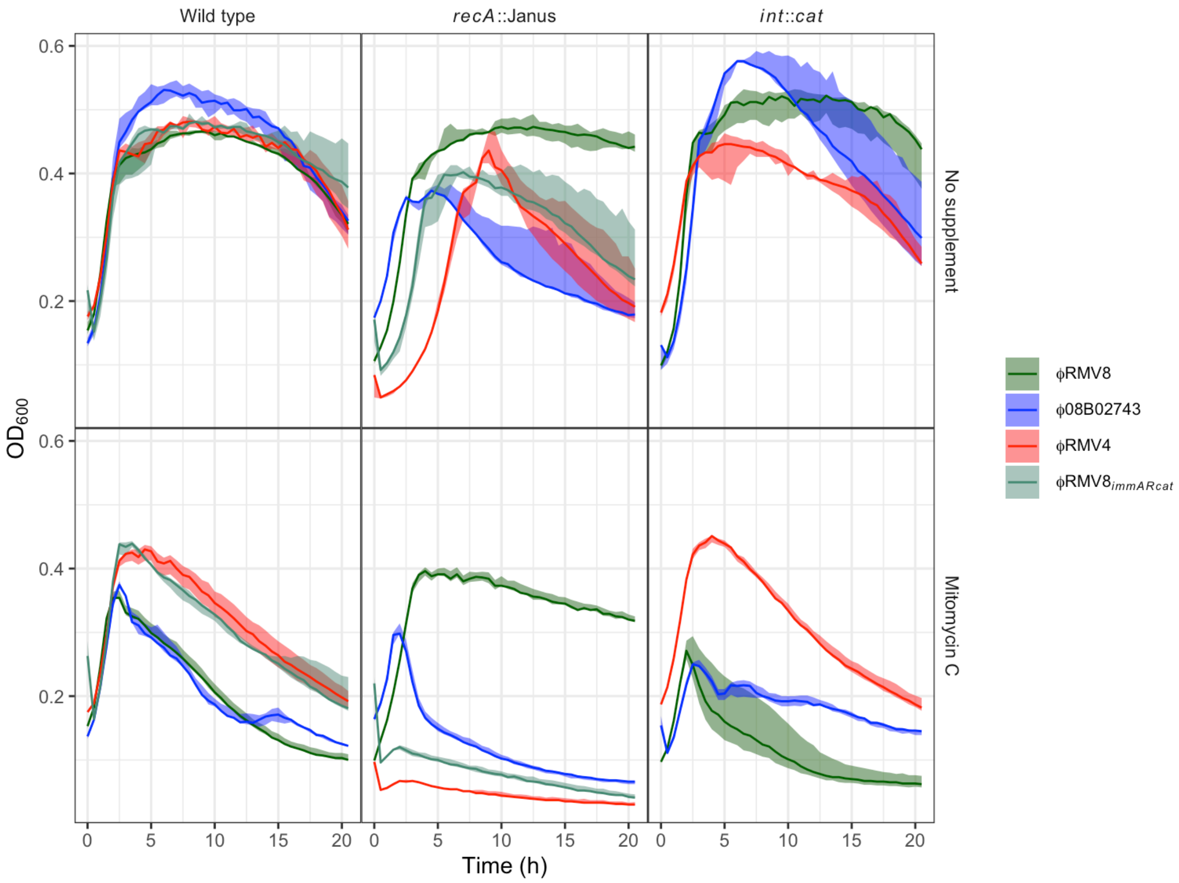


**Figure S21** Growth of prophage systems in unsupplemented media, and in the presence of mitomycin C. Data are shown as in Fig. 2C. These data were combined to generate Fig. 2D.

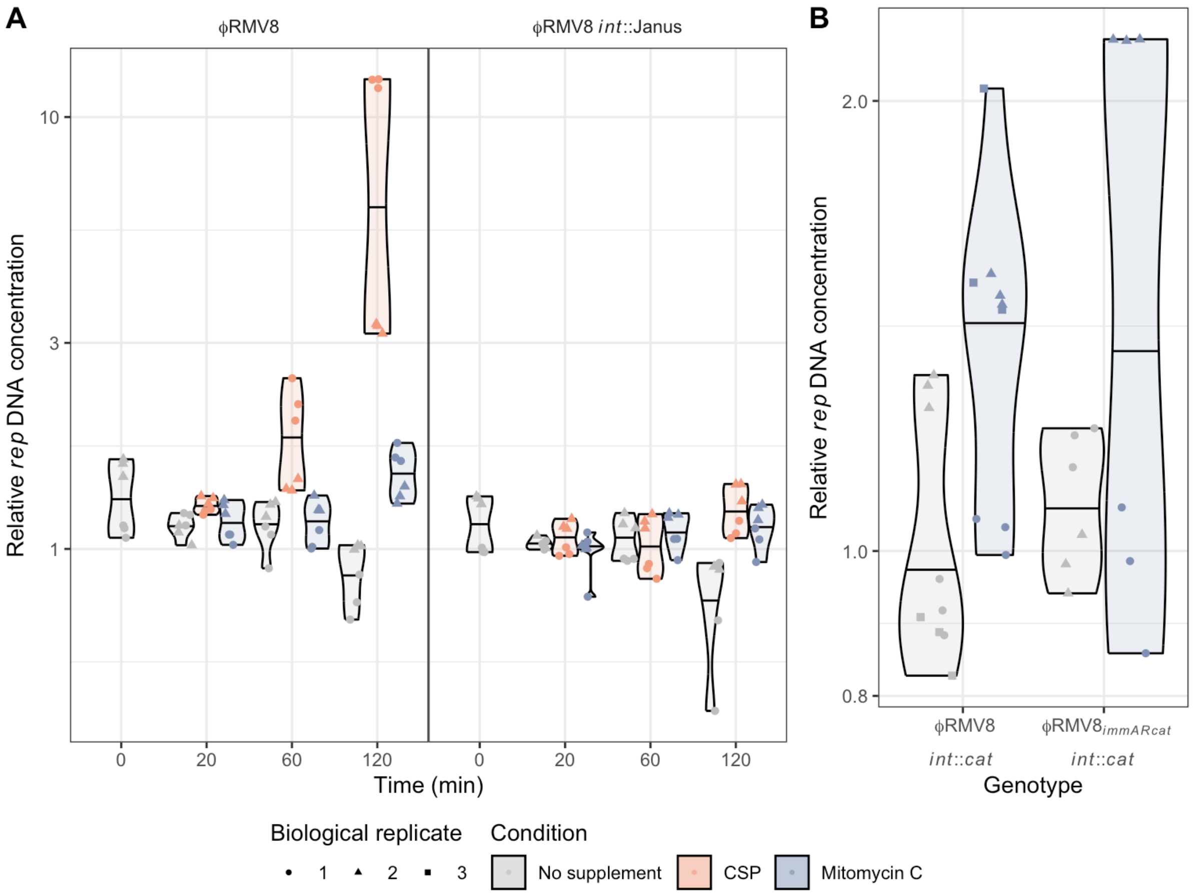


**Figure S22** Testing for *in situ* replication of integrated prophage. (A) Quantitative PCR was used to assay the copy number of the *rep* locus of ϕRMV8 following activation of the wild type prophage, and an *int*::Janus mutant, using two independent biological replicates. Data are shown as in Fig. 3B. The results show that increased copy number of phage DNA was dependent on the element excising from the chromosome, with no evidence of replication of prophage while it remained integrated into the host genome. (B) Quantitative PCR was used to assay the copy number of the *rep* locus of *int*::*cat* mutants of both ϕRMV8 and ϕRMV8$_{immARcat}$. To maximise the sensitivity for detecting replication, samples were extracted after four hours of exposure to mitomycin C, and compared to those grown for an equivalent period in unsupplemented media. No evidence was found of a substantial increase in the DNA copy number of either of the *int*::Janus mutants after exposure to mitomycin C, contrasting with the effect of this stimulus on the wild type prophage observed in Fig. 3B. Therefore ϕRMV8$_{immARcat}$ *int*::Janus shows as little evidence for *in situ* replication as ϕRMV8 *int*::Janus.

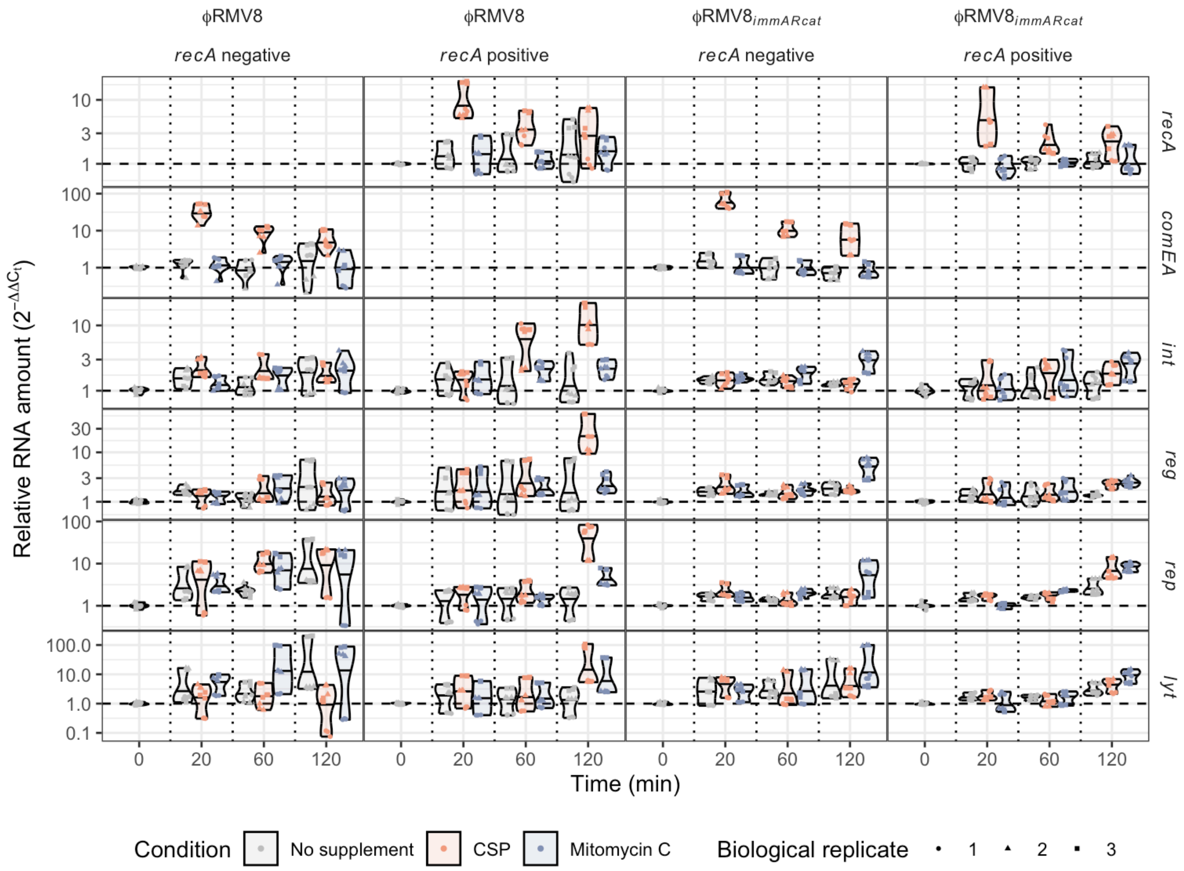


**Figure S23** Effect of RecA on the activation of the ϕRMV8 and ϕRMV8$_{immARcat}$ prophage systems. Quantitative reverse transcriptase PCR was used to assay expression of four prophage loci (*int*, *reg*, *rep* and *lyt*) and *recA*, when this gene was intact*,* or *comEA*, in the *recA*::Janus mutants. Data are shown as in Fig. 3A. These data show RecA was not needed for induction of competence, but it was necessary for the activation of ϕRMV8 by CSP. By contrast, ϕRMV8$_{immARcat}$ exhibited only a weak transcriptional response to both MMC and CSP, which was similar in both the *recA*$^{+}$ and *recA*$^{-}$ genotypes.

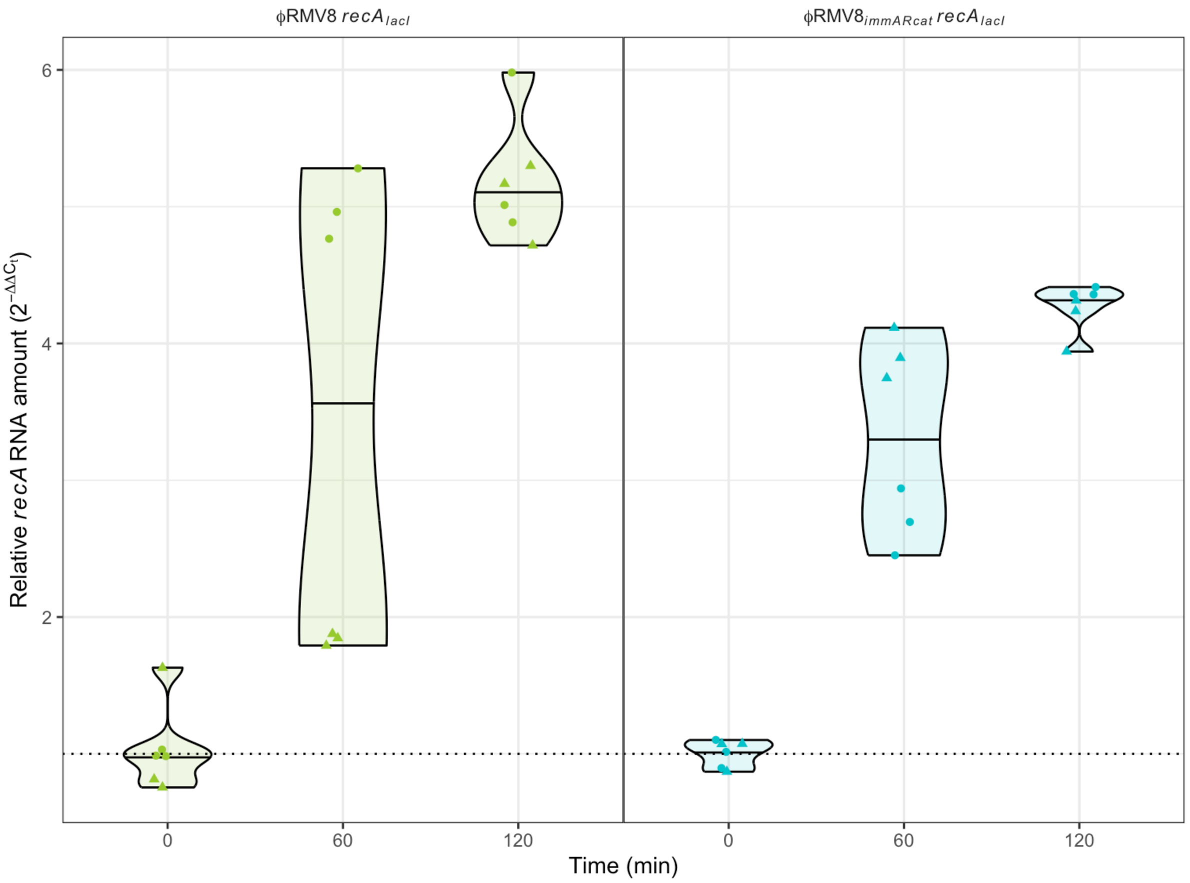


**Figure S24** Validating the IPTG-inducible *recA*$_{lacI}$ genotypes. The expression of *recA* was quantified before exposure to IPTG, and 60 and 120 min after exposure to IPTG, using two independent biological replicates. Data are shown as in Fig. 3A. The dashed line indicates the mean baseline level of expression prior to induction with 1 mM IPTG. These results confirm IPTG caused a consistent, approximately four-fold increase in the expression of *recA*, relative to *rpoA*, in both *recA*$_{lacI}$ genotypes.

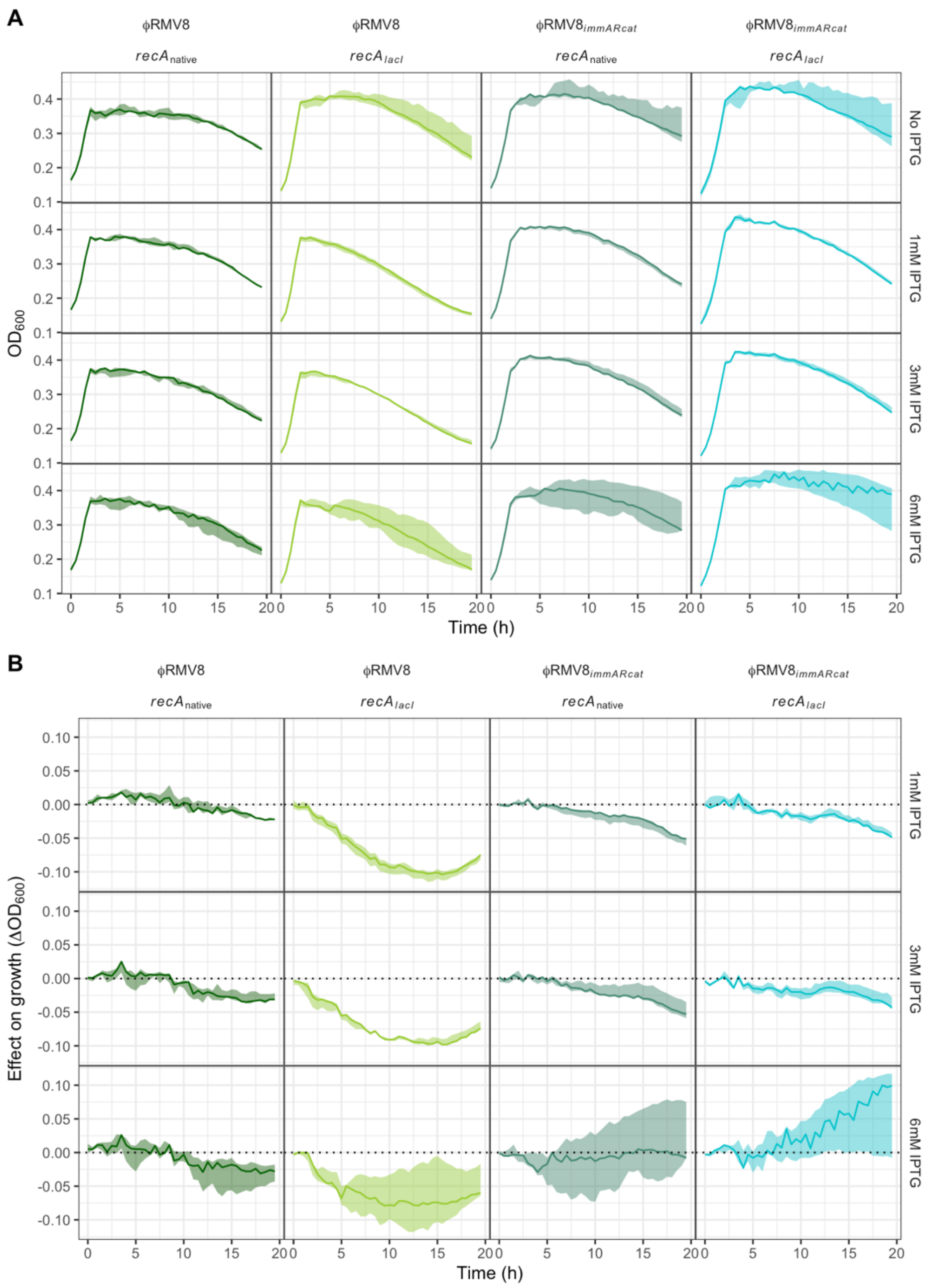


**Figure S25** Effect of IPTG induction on the growth of the ϕRMV8 and ϕRMV8$_{immARcat}$ prophage systems. Cells expressing *recA* under its native promoter, and an IPTG-inducible promoter, were grown in the presence of 5 ng ml$^{-1}$ mitomycin C and different concentrations of IPTG. (A) Each panel shows the growth curve generated by a defined combination of genotype and IPTG concentration. (B) Each plot shows the difference between cells growing in the specified IPTG concentration, and control cultures exposed to the same concentration of mitomycin C without IPTG, using the data shown in panel A, and the approach described in Fig. S15. The ϕRMV8 system was highly sensitive to induction of *recA* by any supplementation with IPTG, resulting in the increased activation of the prophage. No such effect was observed in cells expressing *recA* under its native promoter. The ϕRMV8$_{immARcat}$ system was less sensitive to IPTG. Nevertheless, at the highest concentration of IPTG, prophage induction was sufficiently repressed to increase the growth of cells. This effect was not observed when *recA* was regulated by its native promoter.

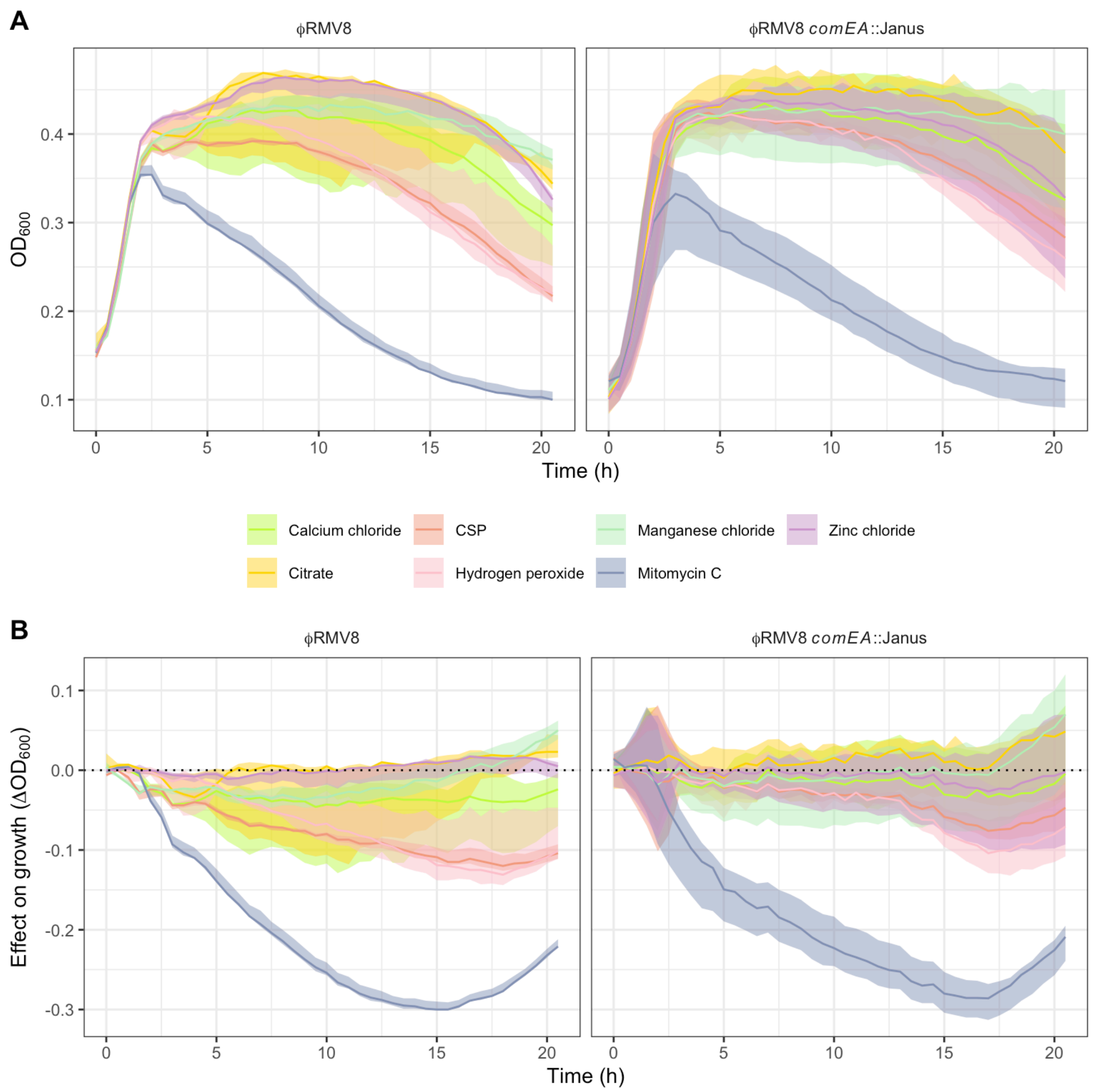


**Figure S26** Role of RecA-ssDNA import on the activation of ϕRMV8. (A) Growth curves for ϕRMV8 and ϕRMV8 *comEA*::Janus, in the presence of different supplements, are shown in separate panels. (B) Using the data shown in panel A, the effect of each supplement is shown as the difference between the optical density of a genotype growing in the presence of the exogenous chemical stimulus, and the median equivalent measurement from the same genotype growing in unsupplemented media, as in Fig. S15. These data demonstrate that although the *comEA*::Janus mutation affects the response of ϕRMV8 to CSP, which depends on import of ssDNA, it does not change the activation to MMC, which generates endogenous ssDNA.

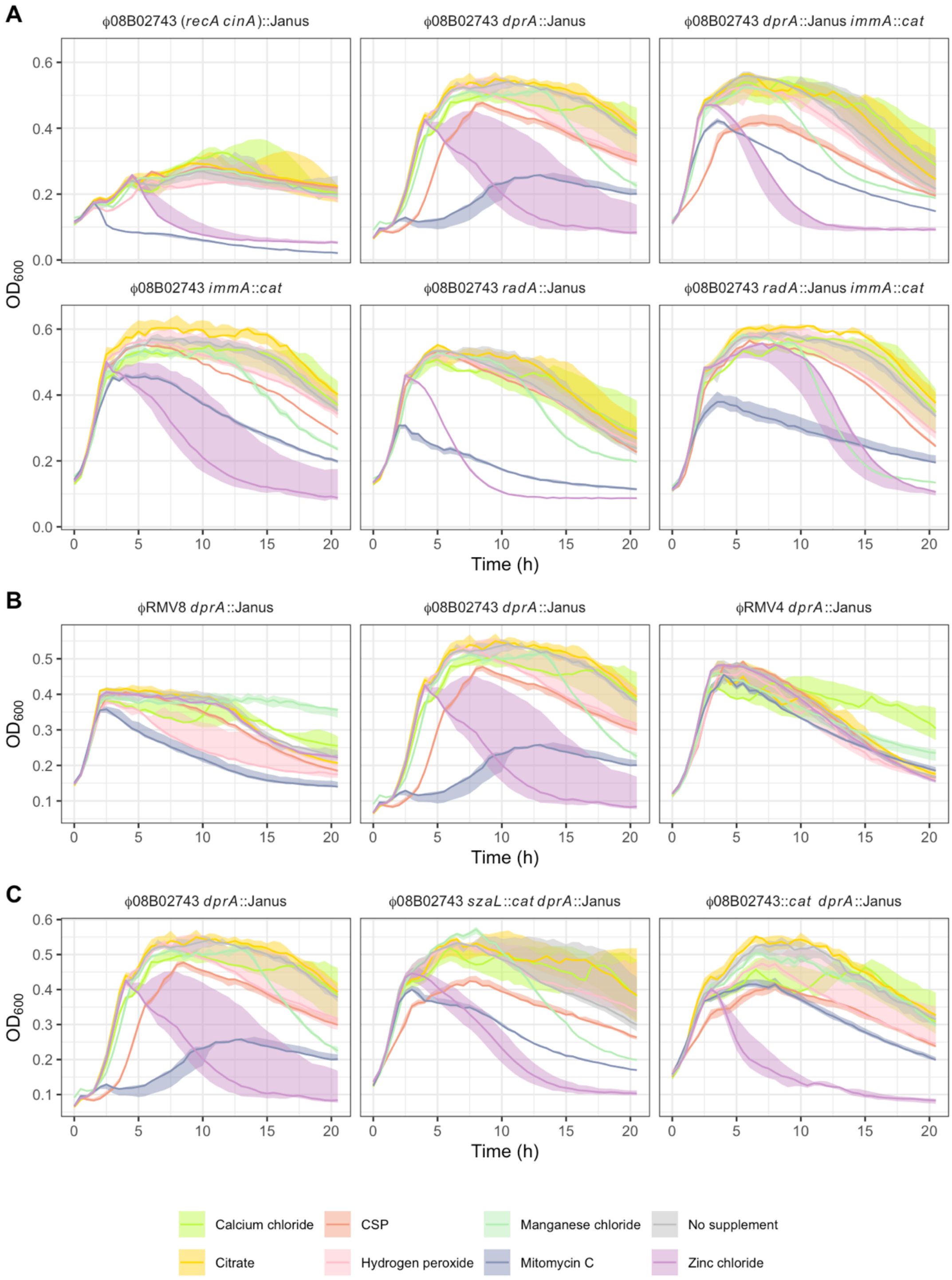


**Figure S27** Growth curves showing the effects of disrupting homologous recombination genes on the regulation of phage induction. Each panel shows the growth curves for a defined genotype in the presence of different supplements. The solid line shows the median $OD_{600}$, and the shaded ribbon shows the range of observed values across at least three biological replicates. (A) Studies of the interactions between cellular homologous recombination proteins and the ImmA prophage activator through quantifying the effects of stimuli on the growth of single and double mutants. (B) The response of prophage to DprA. (C) The interaction between DprA and ϕ08B02743.

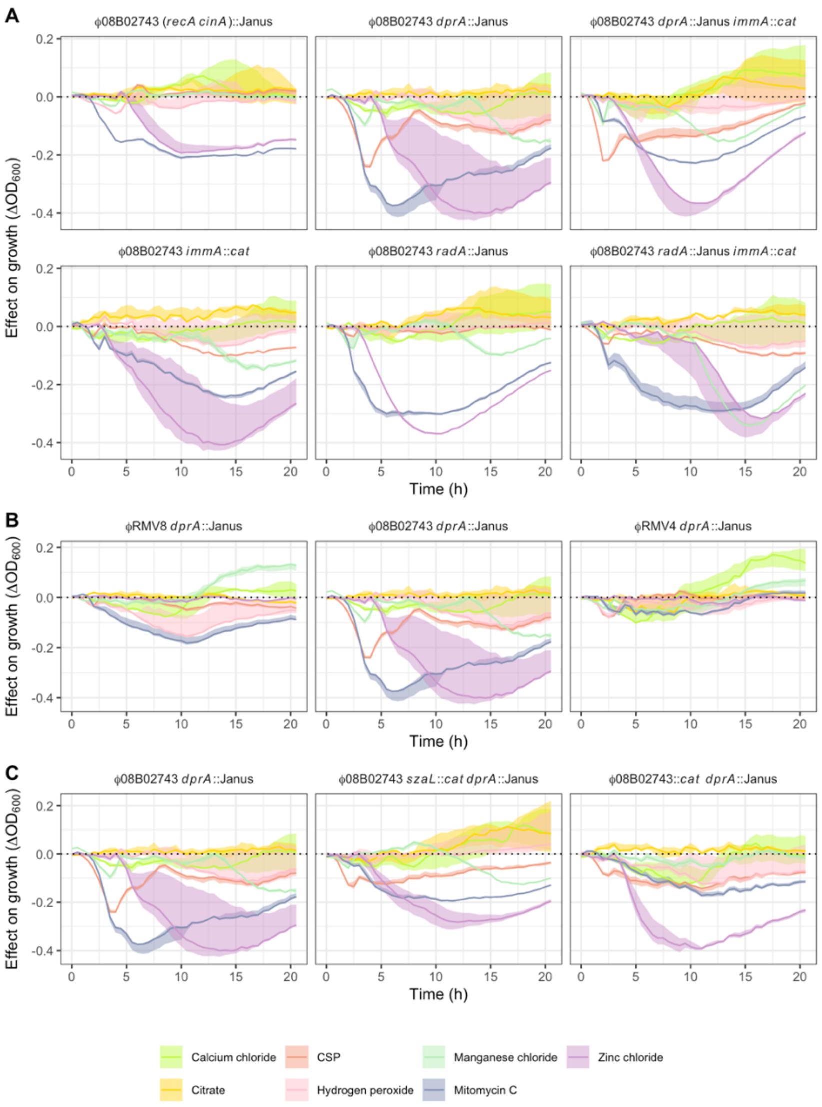


**Figure S28** Plots of relative growth showing the effects of disrupting homologous recombination genes on the regulation of phage induction. These were calculated from the data shown in Fig. S27. Each panel shows the difference between the optical density of a genotype growing in the presence of an exogenous chemical stimulus, and the median equivalent measurement from the same genotype growing in unsupplemented media, as in Fig. S15. (A) Studies of the interactions between cellular homologous recombination proteins and the ImmA prophage activator through quantifying the effects of stimuli on the growth of single and double mutants. These data show that ϕ08B02743 can be induced by CSP in the absence of DprA, in an ImmA-independent manner. (B) The response of prophage to DprA. This demonstrates the strong induction of ϕ08B02743 by CSP in the absence of DprA is not observed in ϕRMV8 or ϕRMV4. (C) The interaction between DprA and ϕ08B02743. This demonstrates the CSP-associated growth inhibition is greater in cells carrying the prophage, implying the observed growth defect is at least partially attributable to prophage activation, rather than a persistent competent state. A similar reduction in the effect of the *dprA*⁻ mutation was observed when *szaL* was disrupted, implying activation was mediated through the SzaL protein.

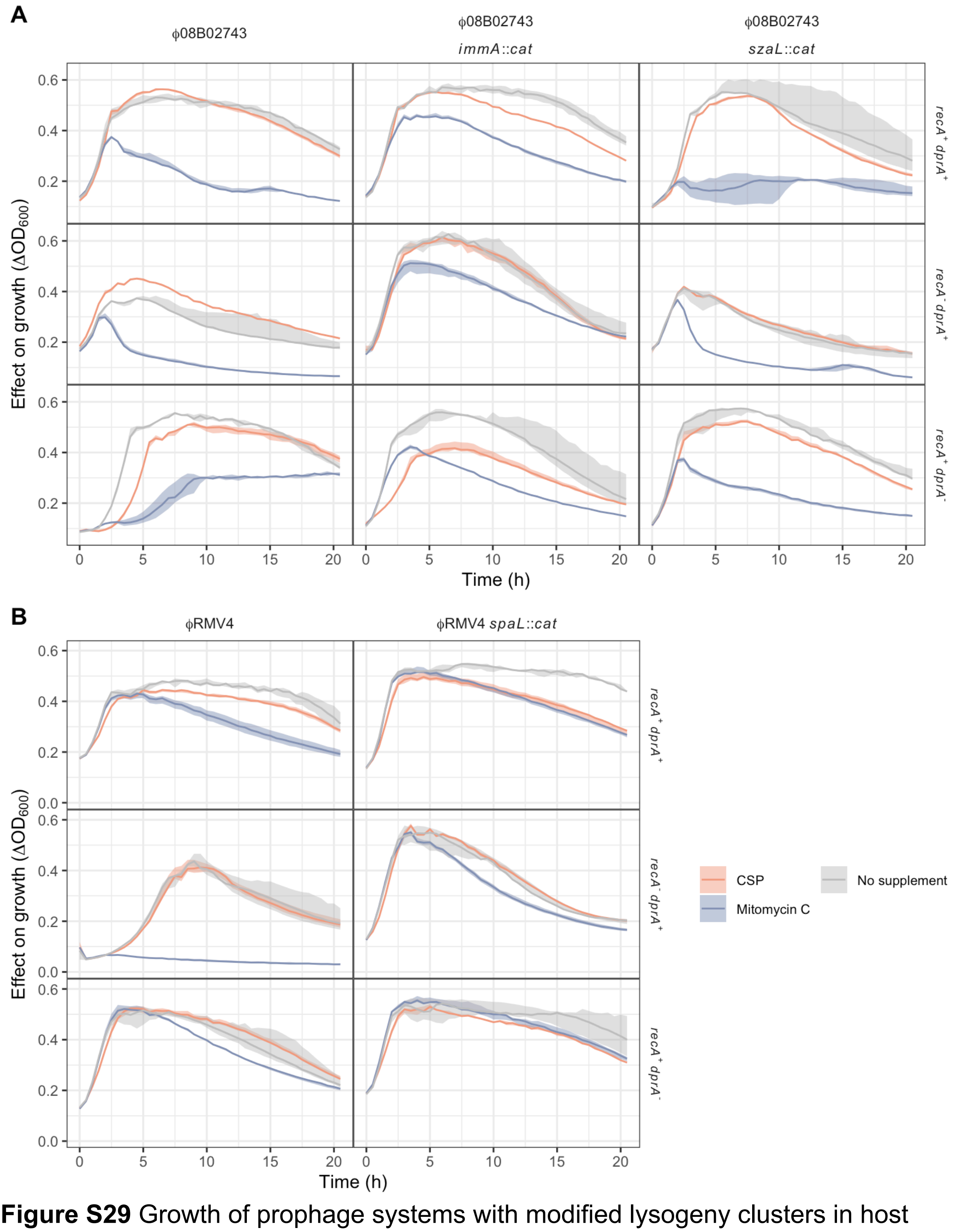


**Figure S29** Growth of prophage systems with modified lysogeny clusters in host cells in which *recA* or *dprA* has been disrupted. Experiments were conducted in unsupplemented media, and in the presence of CSP or mitomycin C. Data are shown as in Fig. 2C. (A) Growth of the ϕ08B02743 system when the prophage is intact, and when the *immA* or *szaL* regulatory genes were disrupted. These data were combined to generated Fig. 5A. (B) Growth of the ϕRMV4 system when the prophage is intact, and when the *spaL* regulatory gene was disrupted. These data were combined to generated Fig. 5D.

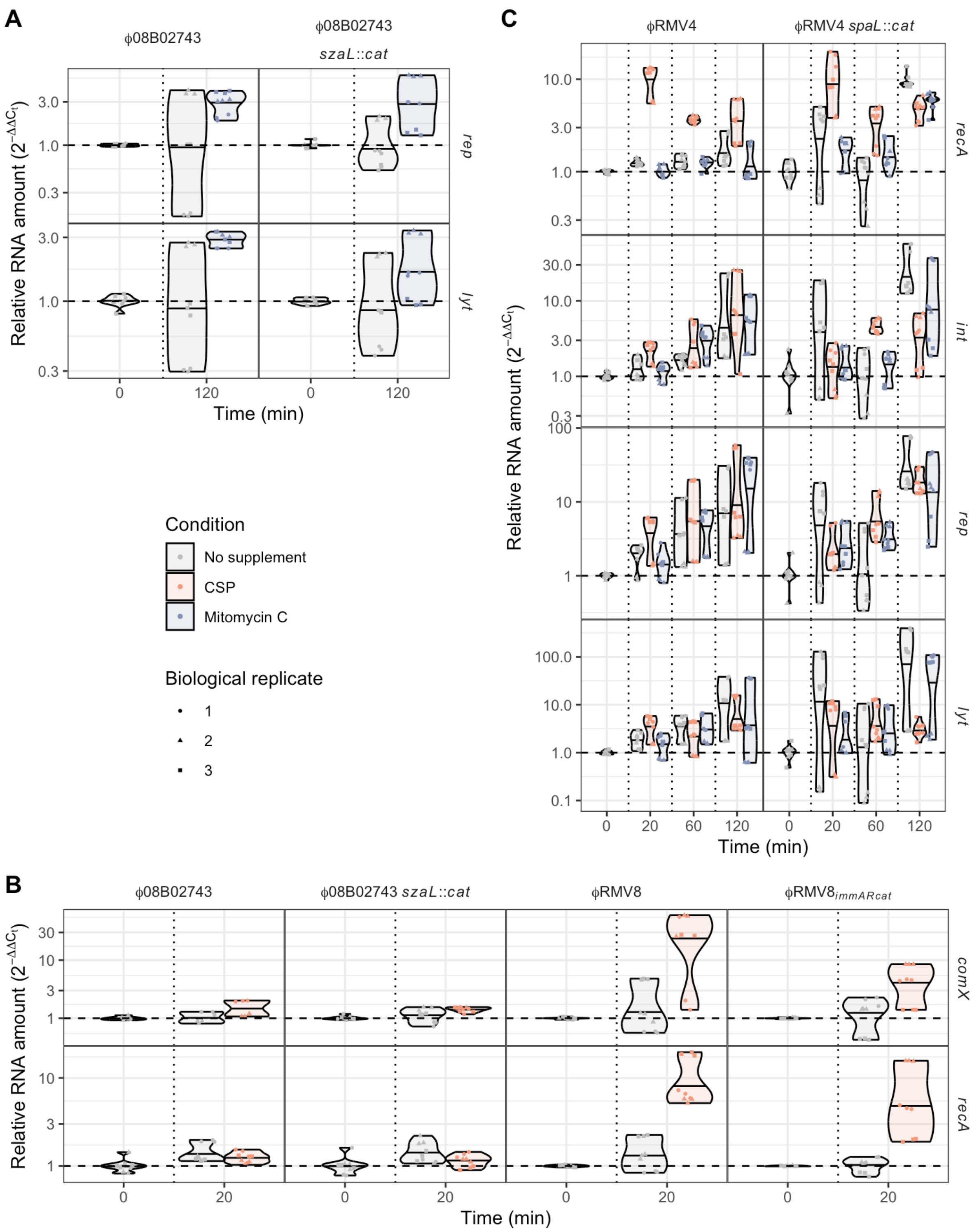


**Figure S30** Quantitative reverse transcriptase PCR assays testing the regulatory roles of the SzaL and SpaL proteins. Data are shown as in Fig. 3A. (A) The effect of the SzaL protein on the activation of ϕ08B02743 by mitomycin C, assayed using the prophage loci *rep* and *lyt*. The disruption of *szaL* by a *cat* gene did not detectably affect the activation of the prophage by mitomycin C during the time in which the virus was susceptible to deletion by transformation, despite the greater sensitivity of the *szaL*$^{-}$ mutant to MMC-induced lysis (Fig. 5A). (B) The effect of the SzaL protein on the expression of the cellular competence machinery. The *recA* gene was not induced by CSP in ϕ08B02743, but was in RMV8$_{immARcat}$, in which *szaL* was disrupted by a *cat* resistance marker. This suggested SzaL may be responsible for repressing *recA* expression. Therefore, the expression of both *recA*, and the competence regulator *comX*, were measured by quantitative reverse transcriptase PCR in genotypes differing in their prophage regulatory loci. The results show that *comX* and *recA* were not induced by CSP in ϕ08B02743 or ϕ08B02743 *szaL*::*cat*, demonstrating the protein is not necessary to prevent the induction of competence in the ϕ08B02743 prophage system. As both ϕRMV8 and $\phi RMV8_{immARcat}$ induced competence efficiently, the ImmAR proteins of ϕ08B02743 are also unlikely to cause the repression of competence induction. (C) Effect of SpaL on the regulation of ϕRMV4 genes and *recA*. Despite the substantial effect of the $spaL^-$ mutation on CSP-associated cell lysis (Fig. 5D), the similarity in the expression of the phage genes in ϕRMV4 and ϕRMV4 *spaL*::*cat* after the induction of competence suggests SpaL does not have a substantial role in transcriptional regulation within the two hour period relevant to deletion of the prophage by HR.

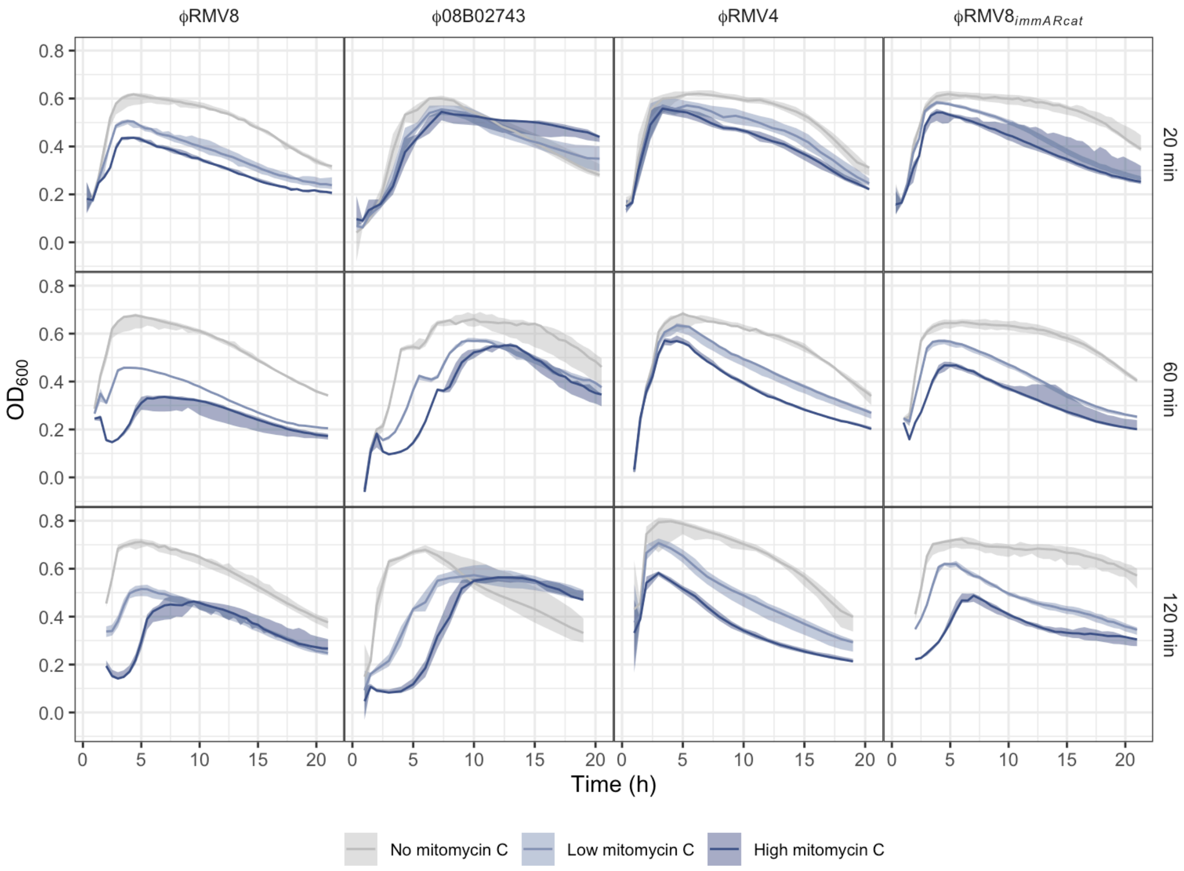


**Figure S31** Effect of different mitomycin C concentrations and pulse durations on prophage activation. The growth curves are displayed as in Fig. 2C. These are the raw data underlying the plots of the differences between growth following the mitomycin C pulses, and growth in unsupplemented media, shown in Fig. 6A.

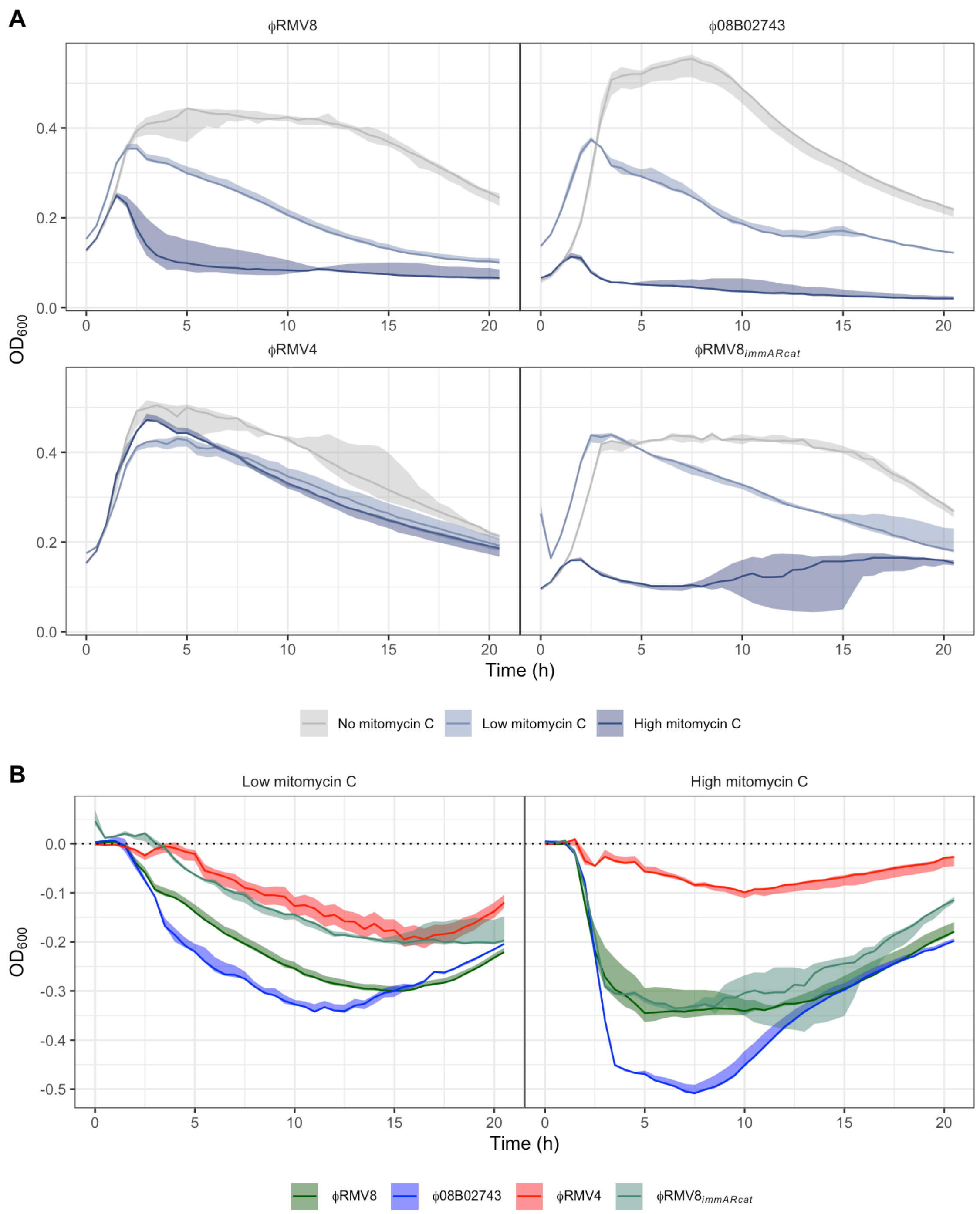


**Figure S32** Effect of different mitomycin C concentrations on the activation of the prophage systems. Each prophage system was grown in a low (5 ng μl$^{-1}$) or high (10 ng μl$^{-1}$) concentration of mitomycin C. (A) Graphs showing the growth of each prophage system in these different mitomycin C concentrations in a separate panel, displayed as in Fig. 2C. (B) The effect of mitomycin C concentration on cell density, relative to parallel growth of the systems in unsupplemented media, is shown using the data displayed in panel A, using the method described in Fig. S15. These plots demonstrate that the kinetics of prophage activation are similar across ϕRMV8, ϕ08B02743 and ϕRMV8$_{immARcat}$. Furthermore, they show that the greater activation of ϕRMV8 by a short pulse of mitomycin C is not a consequence of this system being the most sensitive to any concentration of mitomycin C.

**Table S1** Details of sequence data used to characterise the prophage-pneumococcus systems. The prophage annotations either refer to the position of the prophage sequence within the annotated whole genome sequence, or the DOI by which the relevant locus annotation may be accessed through FigShare.

| Genotype | Serotype | Global Pneumococcal Sequence Cluster | CSP Pherotype | Sequence data type | Accession code | Prophage sequence | Prophage annotation (Position in genome annotation or DOI of manually-curated prophage annotation) |
|---|---|---|---|---|---|---|---|
| *S. pneumoniae* R6 | No capsule | GPSC622 | CSP1 | Complete genome | AE007317 | - | - |
| *S. pneumoniae* ATCC 700669 | 23F | GPSC16 | CSP2 | Complete genome | FM211187 | ClPhR | 26383 - 31653 |
| *S. pneumoniae* 11930 | 23F | GPSC16 | CSP2 | Illumina draft assembly | DAUZQA010000000 | ClPhR | Fragmented |
| *S. pneumoniae* RMV8 | 6A | GPSC97 | CSP2 | Nanopore draft assembly | ERS17697889 | ϕRMV8 | 10.6084/m9.figshare.26425042 |
| *S. pneumoniae* 08B02743 | 19F | GPSC1 | CSP1 | Nanopore draft assembly | ERS17697890 | ϕ08B02743 (representative of ϕARI0460-2) | 10.6084/m9.figshare.26425042 |
| *S. pneumoniae* RMV4 | 9V | GPSC6 | CSP1 | Nanopore draft assembly | ERS17697888 | ϕRMV4 | 10.6084/m9.figshare.26425042 |
| *S. pneumoniae* RMV8$_{immARcat}$ | 6A | GPSC97 | CSP2 | Nanopore draft assembly | ERS17697887 | ϕRMV8$_{immARcat}$ | 10.6084/m9.figshare.26425066 |

**Table S2** Details of the *attB* sites identified in the *S. pneumoniae* genomes.

| Position in *S. pneumoniae* R6 | Name | Accession code of representative prophage insertion | Description of effect on coding sequences |
|---|---|---|---|
| 24066 | $attB_{OXC}$ | FQ312027 | Modifies csRNA3 |
| 24715 | $attB_{scr}$ | ERS1460212 | Partial duplication of *scr* and sequences regulating the *dut* operon |
| 231234 | $attB_{ccnB}$ | ERS1698859 | Modifies csRNA2 |
| 267259 | $attB_{ydiL}$ | ERS1021109 | Interrupts *ydiL* upstream of *folP* |
| 350515 | $attB_{cbpG}$ | ERS638944 | Disruption of protease |
| 438893 | $attB_{pyrG}$ | ERS1299787 | Within RUP element downstream of *pyrG* |
| 926553 | $attB_{ccrB}$ | ERS1813719 | Disrupts *ccrB* within PPI-1 |
| 986969 | $attB_{yjbK}$ | ERS708124 | Disrupts promoter of *yjbK* |
| 998998 | $attB_{hlpA}$ | ERS1699606 | Downstream of *hlpA* |
| 1019764 | $attB_{eno}$ | SAMEA2059239 | Downstream of *eno* |
| 1407631 | $attB_{MM1}$ | FM211187 | Into stop codon of *whiA* |
| 1465604 | $attB_{relA}$ | ERS740678 | In BOX element upstream of *relA* |
| 1624601 | $attB_{pfbA}$ | ERS2527991 | Disrupts *pfbA* |
| 1704863 | $attB_{ssbB}$ | ERS367616 | Disrupts *ssbB* |
| 1713565 | $attB_{ply}$ | ERS1699476 | Downstream of *ply* |
| 1833535 | $attB_{comYC}$ | CP002176 | Disrupts *comYC* |
| 1841795 | $attB_{tgt}$ | ERS455132 | Affects promoter of *tgt* |

**Table S3** Genotypes used in this study.

| Background | Genotype | Shortened Name | Purpose |
|---|---|---|---|
| *S. pneumoniae* R6 | R6 Δ*ivr* | | Characterising the CIPhR element. |
| | R6 Δ*ivr scr*::Janus | R6 *scr*::Janus | |
| | R6 Δ*ivr* $attL_{scr}$ | R6 $scr_{attL}$ | |
| | R6 Δ*ivr* $attL_{scr}$-restored | R6 $scr_{attB}$-restored | |
| | R6 Δ*ivr* $\Delta combox_{dut}$ | R6 $\Delta combox_{dut}$ | |
| | R6 Δ*ivr* CIPhR | R6 CIPhR | |
| *S. pneumoniae* 11930 | 11930 Δ*ivr tvr*::Janus Δ*tvr* | 11930 | Characterising the CIPhR element. |
| | 11930 Δ*ivr tvr*::Janus ΔCIPhR | 11930 ΔCIPhR | |
| | 11930 Δ*ivr tvr*::Janus ΔCIPhR::$attL_{scr}$ | 11930 $scr_{attL}$ | |
| | 11930 Δ*ivr tvr*::Janus ΔCIPhR::CIPhR | 11930 CIPhR restored | |
| *S. pneumoniae* RMV8 | RMV8 Δ*tvrR* | RMV8 $ivr^{+}$ Δ*tvrR* | Assaying the behaviour of a prophage regulated by a C1-type regulator |
| | RMV8 *int*::*cat* Δ*tvrR* | RMV8 $ivr^{+}$ Δ*tvrR int*::*cat* | |
| | RMV8 *lyt*:: *cat* Δ*tvrR* | RMV8 $ivr^{+}$ Δ*tvrR lyt*:: *cat* | |
| | RMV8 Δ*ivr* Δ*tvrR* | ϕRMV8 | |
| | RMV8 Δ*ivr* Δ*tvrR* ϕRMV8::Janus | ϕRMV8::Janus | |
| | RMV8 Δ*ivr* Δ*tvrR int*::*cat* | ϕRMV8 *int*:: *cat* | |
| | RMV8 Δ*ivr* Δ*tvrR rep*:: *cat* | ϕRMV8 *rep*:: *cat* | |
| | RMV8 Δ*ivr* Δ*tvrR lyt*:: *cat* | ϕRMV8 *lyt*:: *cat* | |
| | RMV8 Δ*ivr* Δ*tvrR int*::Janus | ϕRMV8 *int*::Janus | |
| | RMV8 Δ*ivr* Δ*tvrR int* restored | ϕRMV8 *int* restored | |

| | | | |
|---|---|---|---|
| | RMV8 Δ*ivr* Δ*tvrR* ϕRMV8::Janus *rpoB** | ϕRMV8 ϕRMV8::Janus *rpoB** | |
| | RMV8 Δ*ivr* Δ*tvrR* *recA*::Janus | ϕRMV8 *recA*::Janus | |
| | RMV8 Δ*ivr* Δ*tvrR* *int*::*cat* *recA*::Janus | ϕRMV8 *int*::*cat* *recA*::Janus | |
| | RMV8 Δ*ivr* Δ*tvrR* ϕRMV8::*ermB* | ϕRMV8 ϕRMV8::*ermB* | |
| | RMV8 Δ*ivr* Δ*tvrR* [ϕRMV8 *int*::Janus]::*ermB* | ϕRMV8 [ϕRMV8 *int*::Janus]::*ermB* | |
| | RMV8 Δ*ivr* Δ*tvrR* $recA_{lacI}$ | ϕRMV8 $recA_{lacI}$ | |
| | RMV8 Δ*ivr* Δ*tvrR* *comEA*::Janus | ϕRMV8 *comEA*::Janus | |
| *S. pneumoniae* $RMV8_{immARcat}$ | RMV8 Δ*ivr* Δ*tvrR* $c1_{\phi RMV8}$::[$immAR_{\phi 08B02743}$ *szaL*::*cat*] | $\phi RMV8_{immARcat}$ | Assaying the differences between C1-regulation and ImmAR-regulation of prophage induction. |
| | RMV8 Δ*ivr* Δ*tvrR* $c1_{\phi RMV8}$::[$immAR_{\phi 08B02743}$ *szaL*::*cat*] *int*::Janus | $\phi RMV8_{immARcat}$ *int*::Janus | |
| | RMV8 Δ*ivr* Δ*tvrR* $c1_{\phi RMV8}$::[$immAR_{\phi 08B02743}$ *szaL*::*cat*] *recA*::Janus | $\phi RMV8_{immARcat}$ *recA*::Janus | |
| | RMV8 Δ*ivr* Δ*tvrR* $c1_{\phi RMV8}$::[$immAR_{\phi 08B02743}$ *szaL*::*cat*] *dprA*::Janus | $\phi RMV8_{immARcat}$ *dprA*::Janus | |
| | RMV8 Δ*ivr* $c1_{\phi RMV8}$::[$immAR_{\phi 08B02743}$ *szaL*::*cat*] $recA_{lacI}$ | $\phi RMV8_{immARcat}$ $recA_{lacI}$ | |
| *S. pneumoniae* 08B02743 | 08B02743 | 08B02743 $ivr^+$ $tvr^+$ | Assaying the behaviour of a prophage regulated by an ImmAR-type regulator |
| | 08B02743 *int*::*cat* | 08B02743 $ivr^+$ $tvr^+$ *int*::*cat* | |
| | 08B02743 *rep*::*cat* | 08B02743 $ivr^+$ $tvr^+$ *rep*::*cat* | |
| | 08B02743 *lyt*::*cat* | 08B02743 $ivr^+$ $tvr^+$ *lyt*::*cat* | |
| | 08B02743 ϕ08B02743::Janus | 08B02743 $ivr^+$ $tvr^+$ ϕ08B02743::Janus | |

| | | | |
|---|---|---|---|
| | 08B02743 Δ*ivr* Δ*tvrR* | 08B02743 *ivr*$^-$ Δ*tvrR int*::*cat* | |
| | 08B02743 Δ*ivr* Δ*tvrR int*::*cat* | 08B02743 *ivr*$^-$ Δ*tvrR int*::*cat* | |
| | 08B02743 Δ*ivr* Δ*tvrR lyt::cat* | 08B02743 *ivr*$^-$ Δ*tvrR int*::*cat* | |
| | 08B02743 Δ*ivr* Δ*tvr* | ϕ08B02743 | |
| | 08B02743 Δ*ivr* Δ*tvr int::cat* | ϕ08B02743 *int::cat* | |
| | 08B02743 Δ*ivr* Δ*tvr rep::cat* | ϕ08B02743 *rep::cat* | |
| | 08B02743 Δ*ivr* Δ*tvr lyt::cat* | ϕ08B02743 *lyt::cat* | |
| | 08B02743 Δ*ivr* Δ*tvr recA::*Janus | ϕ08B02743 *recA::*Janus | |
| | 08B02743 Δ*ivr* Δ*tvr dprA::*Janus | ϕ08B02743 *dprA::*Janus | |
| | 08B02743 Δ*ivr* Δ*tvr* ϕ08B02743::Janus | ϕ08B02743 ϕ08B02743::Janus | |
| | 08B02743 Δ*ivr* Δ*tvr* ϕ08B02743::Janus *rpoB** | ϕ08B02743 ϕ08B02743::Janus *rpoB** | |
| | 08B02743 ϕ08B02743::Janus *dprA::cat* | ϕ08B02743 *dprA*::*cat* | |
| | 08B02743 Δ*ivr* Δ*tvr szaL::cat* | ϕ08B02743 *szaL*::*cat* | |
| | 08B02743 Δ*ivr* Δ*tvr szaL::cat recA*::Janus | ϕ08B02743 *szaL*::*cat recA*::Janus | |
| | 08B02743 Δ*ivr* Δ*tvr szaL::cat recA*::Janus | ϕ08B02743 *szaL*::*cat dprA*::Janus | |
| *S. pneumoniae* RMV4 | RMV4 Δ*ivr* Δ*tvrR* | ϕRMV4 | Assaying the behaviour of a prophage regulated by a ImmAR-type regulator |
| | RMV4 Δ*ivr* Δ*tvrR int::cat* | ϕRMV4 *int::cat* | |
| | RMV4 Δ*ivr* Δ*tvrR rep::cat* | ϕRMV4 *rep::cat* | |
| | RMV4 Δ*ivr* Δ*tvrR lyt::cat* | ϕRMV4 *lyt::cat* | |
| | RMV4 Δ*ivr* Δ*tvrR recA::*Janus | ϕRMV4 *recA::*Janus | |
| | RMV4 Δ*ivr* Δ*tvrR dprA::*Janus | ϕRMV4 *dprA::*Janus | |

| | | | |
|---|---|---|---|
| | RMV4 Δ*ivr* Δ*tvrR* ϕRMV4::Janus | ϕRMV4 ϕRMV4::Janus | |
| | RMV4 Δ*ivr* Δ*tvrR* ϕRMV4::Janus *rpoB** | ϕRMV4 ϕRMV4::Janus *rpoB** | |
| | RMV4 Δ*ivr* Δ*tvrR* *spaL::cat* | ϕRMV4 *spaL::cat* | |
| | RMV4 Δ*ivr* Δ*tvrR* *spaL::cat recA*::Janus | ϕRMV4 *spaL::cat recA::*Janus | |
| | RMV4 Δ*ivr* Δ*tvrR* *spaL::cat dprA*::Janus | ϕRMV4 *spaL::cat dprA::*Janus | |
| | RMV4 Δ*ivr* Δ*tvrR* *immAR::immAR*$_{08B02743}$-*cat* | ϕRMV4$_{immARcat}$ | |
| | RMV4 Δ*ivr* Δ*tvrR* *immAR::immAR*$_{08B02743}$-*cat recA::*Janus | ϕRMV4$_{immARcat}$ *recA::*Janus | |
| *E. coli* K-12 | *E. coli* K-12 | | Amplification of pLacI repressor |

**Table S4** Oligonucleotides used in this study.

| Purpose | Targeted locus | Forward or reverse | Homologous arm | Sequence |
|---|---|---|---|---|
| Mutant construction | *recA* | Forward | Left | GACGGAGTGACCTATGTCGTCCTTCC |
| Mutant construction | *recA* | Reverse | Left | AAGGGCCCTCTATTCTCCTACATTCTAAT |
| Mutant construction | *recA* | Forward | Right | AAGGATCCGAAGAAGCAGTGAATGAAGAA |
| Mutant construction | *recA* | Reverse | Right | GCAGAGAACCGCTGATAAACCTTGCG |
| Mutant construction | *dprA* | Forward | Left | GAATAGGTGTCATCAAAGGAGAGATTAAC |
| Mutant construction | *dprA* | Reverse | Left | TTGGGCCCTGAACCCAACATTTCCATAAT |
| Mutant construction | *dprA* | Forward | Right | TTGGATCCACTCCATTTCTTTTTCTACTC |
| Mutant construction | *dprA* | Reverse | Right | CCTATCTTGTGATTGTGCTCTTCTCTC |
| Mutant construction | $spaL_{\phi RMV4}$ | Forward | Left | TGGTCGGTGTCGTTCTTCGTTTAG |
| Mutant construction | $spaL_{\phi RMV4}$ | Reverse | Left | AAGGGCCCGATATAGCCACCAGTTTG |
| Mutant construction | $spaL_{\phi RMV4}$ | Forward | Right | AAGGATCCAAAGGCGCTACATCTCAACTA |
| Mutant construction | $spaL_{\phi RMV4}$ | Reverse | Right | TTTGGCAATTCTTCCATCCACATTGC |
| Mutant construction | $szaL_{\phi 08B02743}$ | Forward | Left | AGACTGAATTAGTCGCAGCGTTC |
| Mutant construction | $szaL_{\phi 08B02743}$ | Reverse | Left | AAGGGCCCAATAACCTCCAAAATAATAAC |
| Mutant construction | $szaL_{\phi 08B02743}$ | Forward | Right | AAGGATCCATCGTTCATAATTCGTAAAT |

| | | | | |
|---|---|---|---|---|
| Mutant construction | *szaL*$_{\phi08B02743}$ | Reverse | Right | GTATGAAGCAAGCTATTGTTGTAGGT |
| Mutant construction | Prophage$_{\phi RMV8}$ | Forward | Left | CGTAACCTTGCTGAAAAGAATCGTC |
| Mutant construction | Prophage$_{\phi RMV8}$ | Reverse | Left | AAGGGCCCCGACCTATCTTAAACAAATC |
| Mutant construction | Prophage$_{\phi RMV8}$ | Forward | Right | AAGGATCCTTTTCATAATAATCTCCCTAT |
| Mutant construction | Prophage$_{\phi RMV8}$ | Reverse | Right | TCCTGCATGATAGCTGCGCA |
| Mutant construction | Prophage$_{\phi08B02743/\phi RMV4}$ | Forward | Left | GACTTCGGGCTTGACTTTGTCA |
| Mutant construction | Prophage$_{\phi08B02743/\phi RMV4}$ | Reverse | Left | TAGGGCCCTTCTCATACTACTGAGGATA |
| Mutant construction | Prophage$_{\phi08B02743/\phi RMV4}$ | Forward | Right | TAGGATCCAAAAGGGGCAAAAGTGTCGT |
| Mutant construction | Prophage$_{\phi08B02743/\phi RMV4}$ | Reverse | Right | TGCAGCCTTTTATGCCCACCTACGCC |
| Mutant construction | *int*$_{\phi RMV8}$ | Forward | Left | TCCCCTCTCGTTCAAAAAGTAACTG |
| Mutant construction | *int*$_{\phi RMV8}$ | Reverse | Left | TAGGGCCCTGTATTCTCCTTGTTTATC |
| Mutant construction | *int*$_{\phi RMV8}$ | Forward | Right | TAGGATCCAAAAGTCCCCGCCAATT |
| Mutant construction | *int*$_{\phi RMV8}$ | Reverse | Right | CCATCTGGGATTTTCTTCCCTGAAAAAATATC |
| Mutant construction | *lyt*$_{\phi RMV8}$ | Forward | Left | GGAATGCGGTTGTTTGGTGGAAT |
| Mutant construction | *lyt*$_{\phi RMV8}$ | Reverse | Left | TAGGGCCCTCCCTATCGTCC |
| Mutant construction | *lyt*$_{\phi RMV8}$ | Forward | Right | TAGGATCCATAGAAAGGAAACTTTCTA |
| Mutant construction | *lyt*$_{\phi RMV8}$ | Reverse | Right | CACACGATGATTGAGACGCTC |

| | | | | |
|---|---|---|---|---|
| Mutant construction | *rep*$_{\phi RMV8}$ | Forward | Left | TGCTTACTGGATGACGTGGG |
| Mutant construction | *rep*$_{\phi RMV8}$ | Reverse | Left | TAGGGCCCGATTTTCAGACGATAGAAT |
| Mutant construction | *rep*$_{\phi RMV8}$ | Forward | Right | TAGGATCCTATCCATAACGAGGA |
| Mutant construction | *rep*$_{\phi RMV8}$ | Reverse | Right | CCTAGCTTTTTTCGATACCTGGTG |
| Mutant construction | *comEA* | Forward | Left | TGTGAGGGCGACACAGTTTTC |
| Mutant construction | *comEA* | Reverse | Left | AAGGGCCCCTCTTTGATTTTCTCGATA |
| Mutant construction | *comEA* | Forward | Right | AAGGATCCGACATTATCGACCATCGTGA |
| Mutant construction | *comEA* | Reverse | Right | ATAAGGGAAAAGGGATAAGTCAGCCA |
| Mutant construction | *cat* | Forward | | CAGGGCCCGAAAATTGGATAAAGTGG |
| Mutant construction | *cat* | Reverse | | TAGGATCCCTAAATCTTTAAAATGCCCCTTAAAAT |
| Mutant construction | Janus | Forward | | TTGGGCCCCCGTTTGATTTTTAATGGATAATGTG |
| Mutant construction | Janus | Reverse | | ATGGATCCCCTTTCCTTATGCTTTTGGACG |
| Mutant construction | *cinA* | Forward | Left | GGTGGAGCAGAGCTTGCTATAGAAACC |
| Mutant construction | *cinA* | Reverse | Left | AAGGGCCCGTTTCCTCCTACCTATCTATT |
| Mutant construction | *radA* | Forward | Left | AAGGGCCCCTTGGTTCGATTGACATTGAT |
| Mutant construction | *radA* | Reverse | Left | TCCTCGTAAGAAGGGCTTGGTTTT |
| Mutant construction | *radA* | Forward | Right | CGAACCGTCCGTATAGTAGATGGTT |

| | | | | |
|---|---|---|---|---|
| Mutant construction | *radA* | Reverse | Right | AAGGATCCAAAGACAAGCCAACTAATCC |
| Overlapping PCR | *immAR*$_{\Phi 08B02743}$ | Forward | Left | GAACTCCTTAGCTTGAGGC |
| Overlapping PCR | *immAR*$_{\Phi 08B02743}$ | Reverse | Left | TTAAAAGTCATTCACCTATCAAGAAGCTGATAAG |
| Overlapping PCR | *immAR*$_{\Phi 08B02743}$ | Forward | Middle | ATAGGTGAATGACTTTTAAAGAATATTTACTCAAAGC |
| Overlapping PCR | *immAR*$_{\Phi 08B02743}$ | Reverse | Middle | GTGGGGATTTTTTTGAATGGTTATTTCTTATACATGATTTC |
| Overlapping PCR | *immAR*$_{\Phi 08B02743}$ | Forward | Right | CCATTCAAAAAAATCCCCACACTCTCC |
| Overlapping PCR | *immAR*$_{\Phi 08B02743}$ | Reverse | Right | CCACCTTTCTCGTTGATAGTTATATAATC |
| Mutant construction | *int*$_{\Phi 08B02743}$ | Forward | Left | CGGACCACTTGAGATTGATAG |
| Mutant construction | *int*$_{\Phi 08B02743}$ | Reverse | Left | TA*GGGCCC*TCCATCCACATTGATTTTAACC |
| Mutant construction | *int*$_{\Phi 08B02743}$ | Forward | Right | AA*GGATCC*TTCTCATACTACTGAGATGG |
| Mutant construction | *int*$_{\Phi 08B02743}$ | Reverse | Right | TGCACTTGCGGCTTGACTTTGT |
| Mutant construction | *rep*$_{\Phi 08B02743}$ | Forward | Left | CTGCAATAATTACCCGAGCCA |
| Mutant construction | *rep*$_{\Phi 08B02743}$ | Reverse | Left | TA*GGGCCC*AGAAAGAGGGTGATGCTGCA |
| Mutant construction | *rep*$_{\Phi 08B02743}$ | Forward | Right | TA*GGATCC*CCAATCAGCGAAACCTCTCA |
| Mutant construction | *rep*$_{\Phi 08B02743}$ | Reverse | Right | AAAACGGACACACTTGAGGC |
| Mutant construction | *immA*$_{\Phi 08B02743}$ | Forward | Left | TCTCAGAAATTTACCACCAGCGCTGAC |
| Mutant construction | *immA*$_{\Phi 08B02743}$ | Reverse | Left | AAGGGCCCCATAATTACCTCCACTTACTGT |

| | | | | |
|---|---|---|---|---|
| Mutant construction | *immA*$_{\Phi 08B02743}$ | Forward | Right | AAGGATCCTAATGCTAAAAGCAGATATG |
| Mutant construction | *immA*$_{\Phi 08B02743}$ | Reverse | Right | CATGATTTCCTCACACTCAAAGTTTGGC |
| Mutant construction | *lyt*$_{\Phi 08B02743}$ | Forward | Left | ACCGTTTGGGCGTTCAACCAGTGG |
| Mutant construction | *lyt*$_{\Phi 08B02743}$ | Reverse | Left | TAGGGCCCAACCTGCTCAACGGCATTTA |
| Mutant construction | *lyt*$_{\Phi 08B02743}$ | Forward | Right | TAGGATCCAAAAGGGGCAAAAGTGTCGT |
| Mutant construction | *lyt*$_{\Phi 08B02743}$ | Reverse | Right | TGCAGCCTTTTATGCCCACCTACGCC |
| Mutant construction | *int*$_{\Phi RMV4}$ | Forward | Left | CATCAACCGAGGTCTATATGATGCCTA |
| Mutant construction | *int*$_{\Phi RMV4}$ | Reverse | Left | TAGGGCCCGTTAAAATGGGTATAGTAAAA |
| Mutant construction | *int*$_{\Phi RMV4}$ | Forward | Right | TAGGATCCTTCTCATACTACTGAAGATA |
| Mutant construction | *int*$_{\Phi RMV4}$ | Reverse | Right | GACTTCGGGCTTGACTTTGTCT |
| Mutant construction | *lyt*$_{\Phi RMV4}$ | Forward | Left | ACCGTTTGGGCGTTCAACCAGTGG |
| Mutant construction | *lyt*$_{\Phi RMV4}$ | Reverse | Left | TAGGGCCCTACCTGCTCCACGGCATTTA |
| Mutant construction | *lyt*$_{\Phi RMV4}$ | Forward | Right | TAGGATCCAAAAGGGGCAAAAGTGTCGT |
| Mutant construction | *lyt*$_{\Phi RMV4}$ | Reverse | Right | TGCAGCCTTTTATGCCCACCTACGCC |
| Mutant construction | *rep*$_{\Phi RMV4}$ | Forward | Left | AAAGCAGACACGCTTGAGGC |
| Mutant construction | *rep*$_{\Phi RMV4}$ | Reverse | Left | TAGGGCCCCCAATCAGCGAAACCTCTCA |
| Mutant construction | *rep*$_{\Phi RMV4}$ | Forward | Right | TAGGATCCAGAAAGAGAGTGATGCTGC |

| | | | | |
|---|---|---|---|---|
| Mutant construction | *rep*$_{\Phi RMV4}$ | Reverse | Right | CTGCAATAACTACCTGAGCCA |
| Mutant construction | pLac-*recA* | Forward | Left | GGATCCTCACTGCCCGCT |
| Mutant construction | pLac-*recA* | Reverse | Left | GAGAATGTAGGTGGTGAATGTGAAACCAGTAACG |
| Mutant construction | pLac-*recA* | Forward | Middle | CATTCACCACCTACATTCTCCTGTGTTTTTTTATTTTTG |
| Mutant construction | pLac-*recA* | Reverse | Middle | AAGCTTCCGTTTGATTTTTAATG |
| Mutant construction | pLac-*recA* | Forward | Middle | AAGCTTTACAGTTTATTCTTGAC |
| Mutant construction | pLac-*recA* | Reverse | Middle | TTTTCGCCATAGATCCATTTGCCTC |
| Mutant construction | pLac-*recA* | Forward | Right | TGGATCTATGGCGAAAAAACCAAAAAAATTAG |
| Mutant construction | pLac-*recA* | Reverse | Right | TTCATCTTTGTAAGAATACCAAG |
| Mutant construction | *scr* | Forward | Left | CAGCTAGTCTTGAGCAATTGAAAC |
| Mutant construction | *scr* | Reverse | Left | AAGGGCCCTTATTTTTTCCGTCTATTTC |
| Mutant construction | *scr* | Forward | Right | AAGGATCCATGAAAATTCGTGGTTTTGA |
| Mutant construction | *scr* | Reverse | Right | ACTTAGACATACATTCTAGAAACCGAGA |
| Mutant construction | Modified *attL*$_{scr}$ | Forward | | AAGGGCCCATAGTGGAGCAACAGTTCTGC |
| Mutant construction | Modified *attL*$_{scr}$ | Reverse | | AAGGATCCTTAAATTGTGCCCCTCGTC |
| Mutant construction | Modified *attL*$_{scr}$ | Reverse | | AAGGATCCTGCTCTTTTTTTCGTGCTTTT |
| Mutant construction | *reg*$_{\Phi RMV8}$ | Forward | Left | CTTTTTGACCATTTCTTACCAGGTA |

| | | | | |
|---|---|---|---|---|
| Mutant construction | *reg*$_{ΦRMV8}$ | Reverse | Left | AAGGGCCCGATTATTCACCCCTTGTTT |
| Mutant construction | *reg*$_{ΦRMV8}$ | Forward | Right | AAGGATCCGCATAGCATTCTCGATTTT |
| Mutant construction | *reg*$_{ΦRMV8}$ | Reverse | Right | CATTAGCATAGTTCAGAACTTTTTG |
| Mutant construction | Distal$_{ΦRMV8}$ | Forward | Left | GCACACTTATGATGATGACTTCATC |
| Mutant construction | Distal$_{ΦRMV8}$ | Reverse | Left | AAGGGCCCTCTTCACCTCAATCAAAGT |
| Mutant construction | Distal$_{ΦRMV8}$ | Forward | Right | AAGGATCCAAATAGGGTGGGCGGTTGG |
| Mutant construction | Distal$_{ΦRMV8}$ | Reverse | Right | ATACTGGTTCAGGTAAGATTTTGTTAAC |
| Mutant construction | Distal$_{Φ08B02743}$ | Forward | Left | GTTAGGGTTCTCAGTCATTCCAATCAA |
| Mutant construction | Distal$_{Φ08B02743}$ | Reverse | Left | AAGGGCCCACTTTCAAAGTAAACTTTTAC |
| Mutant construction | Distal$_{Φ08B02743}$ | Forward | Right | AAGGATCCATGAGAAATTCGAATGACAG |
| Mutant construction | Distal$_{Φ08B02743}$ | Reverse | Right | AACGGTTTCTTCAACCCATTGATTTTTA |
| Mutant construction | *reg*$_{ΦRMV4}$ | Forward | Left | CCGACAACCTCAGCCACTTCAT |
| Mutant construction | *reg*$_{ΦRMV4}$ | Reverse | Left | AAGGGCCCAAAAACTCCTCCTTTCTAT |
| Mutant construction | *reg*$_{ΦRMV4}$ | Forward | Right | AAGGATCCTCGAGCGTGACTTTTTTTGA |
| Mutant construction | *reg*$_{ΦRMV4}$ | Reverse | Right | TCTTCTAGTTTTGTTAGAGGAATCCCTC |
| Quantitative PCR | *rpoA* | Forward | | CACGAGCAGGTTCCACTTGA |
| Quantitative PCR | *rpoA* | Reverse | | TGGTCGTGGATATGTACCTGC |

| | | | | |
|---|---|---|---|---|
| Quantitative PCR | *int*$_{ΦRMV8}$ | Forward | | GCCAGAGTGATATCCCTGCT |
| Quantitative PCR | *int*$_{ΦRMV8}$ | Reverse | | CGCTGTCGTTCTGAGTGATG |
| Quantitative PCR | *rep*$_{ΦRMV8}$ | Forward | | CTGAGAATGAGCGAGAGAGGA |
| Quantitative PCR | *rep*$_{ΦRMV8}$ | Reverse | | GGCTCTTACCTGTTCCAGCT |
| Quantitative PCR | *lyt*$_{ΦRMV8}$ | Forward | | GGTGCAGCCTTATCGACAAG |
| Quantitative PCR | *lyt*$_{ΦRMV8}$ | Reverse | | GGACCTACTTGCATGACACG |
| Quantitative PCR | *reg*$_{ΦRMV8}$ | Forward | | ATCCCTACTTCCCCTCTCGT |
| Quantitative PCR | *reg*$_{ΦRMV8}$ | Reverse | | ACAGTGTCATAGTTCCCATCGT |
| Quantitative PCR | *int*$_{Φ08B02743/ΦRMV4}$ | Forward | | ACTCCCCAGGCAAGAAATCA |
| Quantitative PCR | *int* $_{Φ08B02743/ΦRMV4}$ | Reverse | | GCCTCTTGAAGAAAACGCCT |
| Quantitative PCR | *rep*$_{Φ08B02743}$ | Forward | | GTGGTTTGACAGGAAGATTCGA |
| Quantitative PCR | *rep*$_{Φ08B02743}$ | Reverse | | GTATTGGCCCGTGATGTTCC |
| Quantitative PCR | *lyt*$_{Φ08B02743/ΦRMV4}$ | Forward | | GGTGCAGCCTTATCGACAAG |
| Quantitative PCR | *lyt*$_{Φ08B02743/ΦRMV4}$ | Reverse | | CCAACATCCCAACTTCCGTT |
| Quantitative PCR | *immA*$_{Φ08B02743}$ | Forward | | AAGCTAACGTCATGGCCTCT |
| Quantitative PCR | *immA*$_{Φ08B02743}$ | Reverse | | TAACGAGGTCAATAGCCGCT |
| Quantitative PCR | *rep*$_{ΦRMV4}$ | Forward | | GAGTCAAATCGGTGGCGTAC |

| | | | | |
|---|---|---|---|---|
| Quantitative PCR | *rep*$_{\Phi RMV4}$ | Reverse | | AATCTTCCCGTCAACCCACT |
| Quantitative PCR | *scr* left | Forward | | CAGTTCTGCGTGAAGCGG |
| Quantitative PCR | *scr* conserved | Reverse | | AGAGCACACAATTCAAATCGCT |
| Quantitative PCR | *scr full* | Reverse | | CGAAACCAATTCAAAACCACGA |
| Quantitative PCR | *scr*$_{attL}$ *modified* | Reverse | | CCCTCGTCTGCCCCTTTT |
| Quantitative PCR | *recA* | Forward | | GAACATGCCCTTGATCCAGC |
| Quantitative PCR | *recA* | Reverse | | CATACGAGCCTGCAAACCAA |
| Quantitative PCR | *comEA* | Forward | | AGGCTCTGGTTTACGTTCCT |
| Quantitative PCR | *comEA* | Reverse | | TGTCCTGAGCTCGTTTTCCT |
| Quantitative PCR | Φ08B02743 *attB*/*attL* | Forward | | CTGCTGTCAACAACGCTATCA |
| Quantitative PCR | Φ08B02743 *attP*/*attL* | Reverse | | GGGGCAAAAGTGTCGTAAATCT |
| Quantitative PCR | Φ08B02743 *attP*/*attR* | Forward | | GGATATGGAGGATAAACTGGTCA |
| Quantitative PCR | Φ08B02743 *attB*/*attR* | Reverse | | ATAGTCTCAGCACCCCTCTG |
| Quantitative PCR | ΦRMV4 *attB*/*attR* | Reverse | | ATAGCCTCAGCACCCCTCTG |
| Quantitative PCR | ΦRMV8 *attB*/*attL* | Forward | | AAAAGGCTAATCGTTGGGAAATT |
| Quantitative PCR | ΦRMV8 *attP*/*attL* | Reverse | | AATATCAAGGGTTTAGGCGCT |
| Quantitative PCR | ΦRMV8 *attP*/*attR* | Forward | | GGCTCGGTCATGCAAAACTT |

| | | | | |
|---|---|---|---|---|
| Quantitative PCR | ΦRMV8 *attB*/*attR* | Reverse | | CCAACGATACCTGCTGTCA |
| ClPhR excision | 11930_ClPhR_left | Forward | | CAGCTAGTCTTGAGCAATTGAAAC |
| ClPhR excision | 11930_ClPhR_right | Reverse | | ACTTAGACATACATTCTAGAAACCGAGA |
